# Present and future distribution of bat hosts of sarbecoviruses: implications for conservation and public health

**DOI:** 10.1101/2021.12.09.471691

**Authors:** Renata L. Muylaert, Tigga Kingston, Jinhong Luo, Maurício Humberto Vancine, Nikolas Galli, Colin J. Carlson, Reju Sam John, Maria Cristina Rulli, David T. S. Hayman

## Abstract

Global changes in response to human encroachment into natural habitats and carbon emissions are driving the biodiversity extinction crisis and increasing disease emergence risk. Host distributions are one critical component to identify areas at risk of spillover, and bats act as reservoirs of diverse viruses. We developed a reproducible ecological niche modelling pipeline for bat hosts of SARS-like viruses (subgenus *Sarbecovirus*), given that since SARS-CoV-2 emergence several closely-related viruses have been discovered and sarbecovirus-host interactions have gained attention. We assess sampling biases and model bats’ current distributions based on climate and landscape relationships and project future scenarios. The most important predictors of species distribution were temperature seasonality and cave availability. We identified concentrated host hotspots in Myanmar and projected range contractions for most species by 2100. Our projections indicate hotspots will shift east in Southeast Asia in >2 °C hotter locations in a fossil-fueled development future. Hotspot shifts have implications for conservation and public health, as loss of population connectivity can lead to local extinctions, and remaining hotspots may concentrate near human populations.

## Introduction

Major current and future global changes pose a severe risk to biodiversity and human survival [1]. Global climate change and human encroachment into natural habitats is simultaneously driving the biodiversity extinction crisis and increasing disease emergence risk [2]. Climate and land cover change will alter the range distribution of species [3], an important, poorly defined predictor of zoonotic (animal to human) disease risk [4,5], and the direction and magnitude of range shifts are not estimated for many species, leaving the impacts on their viral interactions uncertain [6,7].

In early 2020, genomic analysis identified the severe acute respiratory syndrome coronavirus 2 (SARS-CoV-2), responsible for the coronavirus disease 2019 (COVID-19) pandemic. SARS-CoV-2 is closely related to viruses present in the intermediate horseshoe bat *Rhinolophus affinis* (virus RaTG13, sampled from the Yunnan province of China in 2013 [8]), and the following bats captured in northern Lao PDR in 2020: *Rhinolophus malayanus* (virus RmYN02), *Rhinolophus marshalli* (virus BANAL-236), and *Rhinolophus pusillus* (virus BANAL-103) [9]). These viruses, like SARS-CoV-2, are part of the *Sarbecovirus* subgenus, and belong to the *Betacoronavirus* genus (family *Coronaviridae*, subfamily *Orthocoronavirinae*). In the year since SARS-CoV-2 was described, 41 potential *Betacoronavirus* hosts have been reported [10]. Knowing hosts and where hosts are gives insights into emerging disease origins and clues on future risk [11,12].

Bats comprise ∼20% of global mammal taxonomic diversity, with more than 1435 species described [13], among 6400 known mammals. This diversity likely contributes to the viral diversity in bats [14,15], including some that have emerged as pathogenic in people [16,17], such as sarbecoviruses [12]. Bats have multiple functions in ecosystem dynamics as pollinators, seed dispersers and insect predators [18]. However, over a fifth of species are Threatened or Near Threatened with extinction (IUCN 2021) and drivers of changes in bat distributions, such as land use change, likely contribute both to population declines and the simultaneous increase in infectious disease emergence risk [11,19]. Improved range estimates can, therefore, support conservation strategies and understanding disease risk.

Among factors influencing bat distributions, suitable climatic limits and karst are critical for many species [20–22]. The presence of native habitat, especially dense forest is also vital for many species [3,23]. Maintaining such habitats has important implications for conservation and potentially viral transmission through changes in species interactions and survival probability [7]. For example, host dispersal among vampire bats, and specifically males, has facilitated rabies spread in Peru [24] and sympatry has led to host shifts among bat coronaviruses [25,26].

There are, however, knowledge gaps ranging from bat distributional ecology to their behaviour, immunity, and physiology [15]. Attempts to estimate bat sarbecovirus-host spatial distributions have included modelling near current distributions for Rhinolophidae in Southeast Asia [27] and filtering their expected areas of habitat [12]. However, estimating species distributions with future projections can help understand their conservation status, inform land use planning to avoid conflicts [28], generate better models for estimating the risk of emerging novel pathogens, and allow targeted infectious disease surveillance.

The rapid increase in bat data after the COVID-19 pandemic provides opportunities to better understand bats’ distributional ecology, but may bring sampling biases in areas where surveillance has been greatest [11,29,30]. Avoiding misprediction is essential, and we have the opportunity to update ecological niche models with the help of big data, reproducible tools and open science. We need adequate inferences regarding bat species distributions from the current period projected to proximate future scenarios, so we can establish guidelines for how to transition from the current trajectory of biodiversity loss and pandemic risk to a more sustainable future [1].

Here, we use ecological niche models to assess the potential distribution of bat hosts of *Sarbecovirus* to identify: a) What is the availability and spatial coverage of data for inferring the distribution of bats known to host sarbecoviruses?; b) How are *Sarbecovirus* bat hosts distributions affected by climate, karst and forest amount in the near current and future scenarios?; and c) Where are current and future areas with high species richness (hotspots) of *Sarbecovirus* bat hosts? Finally, we share a dynamic pipeline, considering the inevitable addition of new data on host species in the future.

## Methods

### Target species and occurrence data

All analyses were in R 4.1.0 [31] (Figure S1). Code is provided in a GitHub repository (https://github.com/renatamuy/dynamic) and data in Zenodo (TBD, https://drive.google.com/drive/folders/1kBAi4eIGYRLXF7HLBc7QjpqyH4mHBEuu?usp=sharing). We spatially predicted the occurrence of all known *Sarbecovirus* hosts regardless of the first viral detection location using Ecological Niche Models (ENMs) and approximating them to species distribution models (SDMs). We compiled host data (Table S1) from published articles, preprints, and NCBI (National Center for Biotechnology Information) accession numbers cross-checked in Virion v0.2.1 [32] with Genbank references. Bat hosts of *Sarbecovirus* viruses were: (1) explicitly named in the reference; and (2) the source of viruses or viral fragments of *Sarbecovirus* or synonyms for SARS-related coronaviruses.

Then, we mined bat host species occurrences in September 2021 from: Darkcides v1 [22], Global Biodiversity Information Facility (GBIF), Berkeley Ecoinformatics Engine (Ecoengine), Vertnet, Integrated Digitized Biocollections (IDigBio), iNaturalist, Obis, Vertnet, and data compiled for previous publications [11,33]. We filtered all data sources, and with iNaturalist kept only ‘research quality grade’ data. We performed data mining using a custom loop through spocc function [21]. We only kept records from 1970 onwards, and records from before 2000 comprised only 4% of records. All points from DarkCideS (N= 1351) are from 2000’s onwards.

### Sampling bias assessment

To reduce spatial sampling bias due to uneven and undersampling [34,35], we performed a series of filtering steps, removing data concentrated in political centroids of countries, provinces, national capitals, and centroids for GBIF headquarters and museums. Duplicate coordinates for the same species and points in permanent water bodies and oceans were excluded with clean_coordinates from CoordinateCleaner 2.0-18 [36] and cc_outl used for removing geographic outliers, defined by the interquartile range. Because the number of points can define outliers differently and lead to information loss for rare data, we set 20 as the minimum value for correction. We then visually inspected points in QGIS 3.10.7.

We inspected points for taxonomic and expert range consistency based on IUCN Red List evaluations and the Handbook of the Mammals of the World when an IUCN polygon was not available [37]. We used IUCN Red list habitat and population trend information to discuss our findings. For species complexes, the broader range matching the viral detection information was considered to later intersect with input data points. Species complexes like *H. pomona* and *H. ruber* were treated as *lato sensu* across their distributions. For *H. pomona*, we considered the broad distribution for *H. gentilis* as a mask for filtering occurrences for the IUCN-intersected input. *H. ruber* occurrences were kept within and outside their matching IUCN polygons, since range and taxonomy may need review. Occurrences of *Rhinolophus cornutus* in continental areas from DarkCides were not considered, as they refer to *R. pusillus* in recent assessments [38].

Final preprocessing point thinning was made in ENMTML [39], using a 13 km approximated search cell radius size. We did not perform further environmental filtering as the largest gains in model performance using those filters are for 10 points or fewer [40] and we only ran models with more than 40 points, as is good practice [41].

We assessed accessibility bias for the final thinned data (*Sarbecovirus* host presence regardless of species) considering the distance of occurrence points from cities, rivers, roads and airports through sampbias [42], which estimates how sampling rates (a Poisson sampling process with rate λ) vary as a function of proximity bias to drivers, generating a sampling coverage metric driven by accessibility bias. We assume that areas with low sampling coverage driven by accessibility bias should be more investigated, especially if SDMs predict a suitable habitat for a high numbers of species. After generating the layers including estimated sampling rates (input parameters in Table S2), we visualized which highly species-rich areas predicted by our ensembles have high values of estimated sampling proportions with a bivariate choropleth map.

### Accessible area and spatial restriction

Accessible areas where species may disperse were defined by ecoregion extents [43] of locations where each species occurred (IUCN polygon intersected and non-intersected). This extent was used for geographically and environmentally constrained background point sampling [44] with the default ratio between presence and background points around a 50 km buffer from occurrence points. Data partitioning for training:testing was 75:25% by split sample. We included dispersal capability in the model ensembles using the *a posteriori* method ‘OBR’ for SDMs. This method was coupled within the accessible areas for spatial restriction of the final maps, as it performed well in virtual species tests [45], reducing overprediction without increasing omission errors. Because the ‘OBR’ method [45] is not applied to future projections as we cannot restrict them based on observed future occurrences, we compared all projected ranges within the accessible area in future scenarios as being the maximum limit for dispersal. We define range as the area where the species most likely occurs (estimated occupied area) driven by the environmental covariates used.

### Covariates and workflow

We defined environmental variables that are important drivers of target species distributions in the current geographic space, which can also be projected into the future. Based on the focal species’ ecology, we chose selected climatic (annual precipitation, precipitation seasonality, annual mean temperature, temperature seasonality) and landscape variables (karst and primary forest cover) as covariates (Table S3).

Habitats used by each species were extracted using rredlist v0.7 (Figure S2). We selected forest cover as the main land cover variable to avoid correlation and error inflation due to limited data. Furthermore, most of our target species benefit from forest physiognomies (Figure S2). Forest habitat was calculated as a proportion from LUH2 sensu [46] for near current and future scenarios at 0.25, so we resampled all other layers through bilinear interpolation to match this resolution. Distance to cave or karst is the only static variable. We calculated the minimum euclidean distance (km) from karst or cave and averaged it within the working resolution grid. Distances were calculated in QGIS 3.10.7 after warping layers to Mercator metric projection Datum WGS84 and then reprojected back to the geographic system with Datum WGS84.

Climate predictors were downloaded from WorldClim 2.1 [47] with 10 arc-minutes spatial resolution. We selected covariates with Pearson’s product-moment correlation value |r| < 0.70 [48] across all covariates prior to modelling to avoid collinearity (Figure S3). From 19 near current (1970-2000) bioclimatic variables, we selected annual mean temperature (bio_1), temperature seasonality (standard deviation ×100, bio_4), annual precipitation (bio_12), and precipitation seasonality (coefficient of variation, bio_15).

We built ENM ensembles using ‘MXS’ and ‘MXD’ maximum entropy algorithms. We selected these algorithms based on experience and performance [45,49,50]. We used consensus ensemble maps including the suitability values weighted by TSS (weighted mean of True Skill Statistics values) and report performance using TSS, but also report Boyce discrimination values [51]. Each model algorithm was replicated 10 times through the ‘bootstrap’ term in ENMTML. Despite being called bootstrap, this method applies a split sampling method. To evaluate model performance, we randomized occurrence data into 75%:25% train:test samples to calculate the TSS [34] for each model. Models with TSS higher than 0.5 were considered as performing above that expected by chance [52]. We weighted the ensembles based on model performance and used weighted TSS value differences for selecting the most realistic maps between IUCN intersected and non-IUCN-intersected data. Threshold values were calculated to transform each model’s predictions to presence or absence of each species, using ‘MAX_TSS’, the threshold at which the sum of the sensitivity and specificity is the highest [53]. After generating species maps, we calculated host richness as the sum of presences of each host per pixel in the binary maps, producing a map of host– species taxonomic richness. We derived zonal statistics based on the host species map for the country’s shapefile, calculating the maximum predicted host richness per country using administrative regions from Natural Earth (https://www.naturalearthdata.com/). We compared estimated taxonomic richness maps generated from the two sets of models to check for convergent patterns using Pearson’s product-moment correlation. We ran models for two sets of filtered and thinned datasets: 1) IUCN-intersected polygon data; and 2) non-IUCN polygon intersected data. We compared performance values between those two sets of ensembles to infer improvement in performance. Correlative variable contribution was inspected throughout each set of model algorithms.

### Hotspots and future projections

We downloaded SSP (Shared Socioeconomic Pathways) scenarios for Coupled Model Intercomparison Project Phase 6 (CMIP6) at 10 arc-minutes spatial resolution. We present results for projections using the SSP 245 and SSP 585, focusing on SSP585 for our results. SSPs represent baseline paths with varying human behaviour reflected in actions changing land cover, and coupled with gas emissions they provide future global change scenarios. Scenario SSP 585 means 8.5 radiative forcing level coupled with an SSP5 (Fossil-fueled Development), which translates in a pessimistic scenario, for instance [54]. The SSP 245 is known as a middle-of-the-road scenario. Here, we present results for BCC-CSM2-MR (Beijing Climate Center Climate System Model) and CanESM5 (Canadian Earth System Model version 5) GCMs for 2021-2040, 2041-2060, 2061-2080, 2081-2100 periods. We report values for SSP585 and BCC-CSM2-MR for 2100 in our maps, as a more extreme, but possible future scenario. We include results for all scenarios, periods and GCMs in the supplements. Our pipeline easily incorporates all combinations of SSPs, periods and GCMs, depending on computational power. We resampled all future rasters to 0.25 dd using the bilinear method. Area calculations of range shifts and shifts in range overlap were made for each consensus binary map. Hotspots were simply defined as pixels with the greatest predicted species richness. The hotspots’ centroid in the present and future were calculated to describe changes in climate and their locations. To investigate if hotspots were getting physically hotter we inspected the distribution of richness values and average temperature in the present and by 2100 for SSP585 and SSP245.

## Results

Sarbecoviruses were reported from 35 bat species (Table S1). We could model 17 species using IUCN-intersected data, and 23 with non-intersected data (Figure S4). From the 23 species, six could only be modelled without intersecting their points with IUCN data; nine species did not show improvement in TSS values after intersecting them with IUCN ranges, while eight had small improvements after cropping occurrences within IUCN range limits (Table S4).

The maps show three focal areas of suitability across species; one each in Western Europe, Indochina, and Central Africa (Figure S5, S6). IUCN delimitation for occurrence inclusion does not improve model performance for more than 10% added value in TSS in most cases (Table S4). Richness maps for the two datasets were highly positively correlated (|r| 0.955; p-value <0.0001), indicating agreement between richness hotspots for IUCN-intersected data and data gathered inside and outside IUCN polygons. Therefore, we report the non-IUCN intersected datasets. Overall, species with smaller ranges had fewer filtered points (Figure S7).

Environmental covariates affected *Sarbecovirus* bat hosts differently, with temperature seasonality (n = 12) and karst (n = 5) the top ranked variables for most species, and other variables varied in contribution, with differing responses considering the correlative covariate importance (Figure S8).

The highest number of bat species (i.e. host hotspots) in the present and future projections occurred in Southeast Asia (Figure 1; Table 1). The highest values were in Myanmar (13 species), then China, Lao PDR, Thailand, and Vietnam (12 species). Area changes are visible in SSP 585 in 2100, and highest richness values are less continuous in the future due to projected species losses (Figure S9).

**Table 1.** Highest national maximum *Sarbecovirus* host richness values (i.e. hotspots) predicted through SDMs.

| Country | Subregion | Hosts | Median sampling rate<br>(Mean +/- SD) |
| --- | --- | --- | --- |
| Myanmar | Southeast Asia | 13 | 0.033 (0.053 +/- 0.061) |
| China | East Asia | 12 | 0.024 (0.051 +/- 0.064) |
| Lao PDR | Southeast Asia | 12 | 0.052 (0.063 +/- 0.041) |
| Thailand | Southeast Asia | 12 | 0.087 (0.102 +/- 0.067) |
| Vietnam | Southeast Asia | 12 | 0.086 (0.107 +/- 0.074) |
| Cambodia | Southeast Asia | 8 | 0.112 (0.127 +/- 0.076) |
| India | South Asia | 8 | 0.069 (0.09 +/- 0.078) |
| France | West Europe | 8 | 0.128 (0.144 +/- 0.076) |
| Italy | South Europe | 8 | 0.146 (0.152 +/- 0.079) |
| Malaysia | Southeast Asia | 8 | 0.051 (0.074 +/- 0.072) |

**Figure 1.**
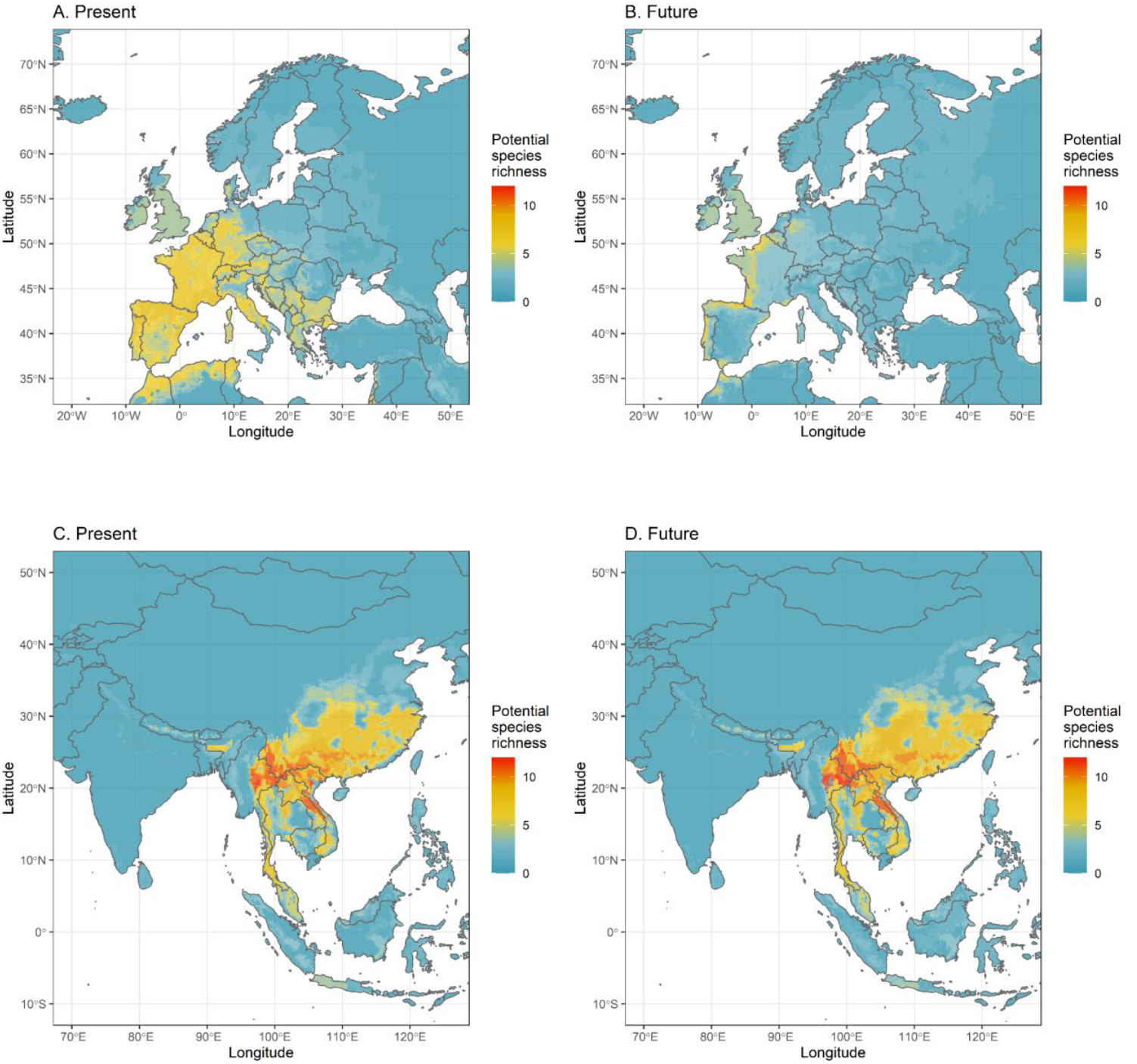
Present (A, C) and future (B, D) distribution of *Sarbecovirus* bat host species richness, mostly peaking in Europe and Southeast Asia. Projections for *Sarbecovirus* bat host species richness consider the period 2080-2100 (SSP585 scenario, BCC-CSM2-MR global circulation model).

From the top 10 countries for maximum potential species richness in a pixel, Italy had the highest estimated sampling rate. Figure 2 shows the interaction between estimated *Sarbecovirus* bat hosts richness with estimated sampling rates. We highlight areas where the number of species is high and sampling proportion low as future priorities for data collection. Sampling rates are mostly correlated with the distance from roads (Figure S10). Overall the estimated sampling rates through accessibility were low for individual locations (min=0, median=0.013, mean=0.045, 3rd q.=0.065, max=0.49), even when species are present by ENMs (min=0, median=0.08, mean=0.098, 3rd q.=0.141, max=0.446). In Europe, sampling rates show high accessibility bias and better coverage of hosts than other regions, but regions in Southeast Asia are also well sampled, especially eastern coastal areas (purple in Figure 2). Highest values for richness were estimated for areas with low sampling rates (Figure S11).

**Figure 2.**
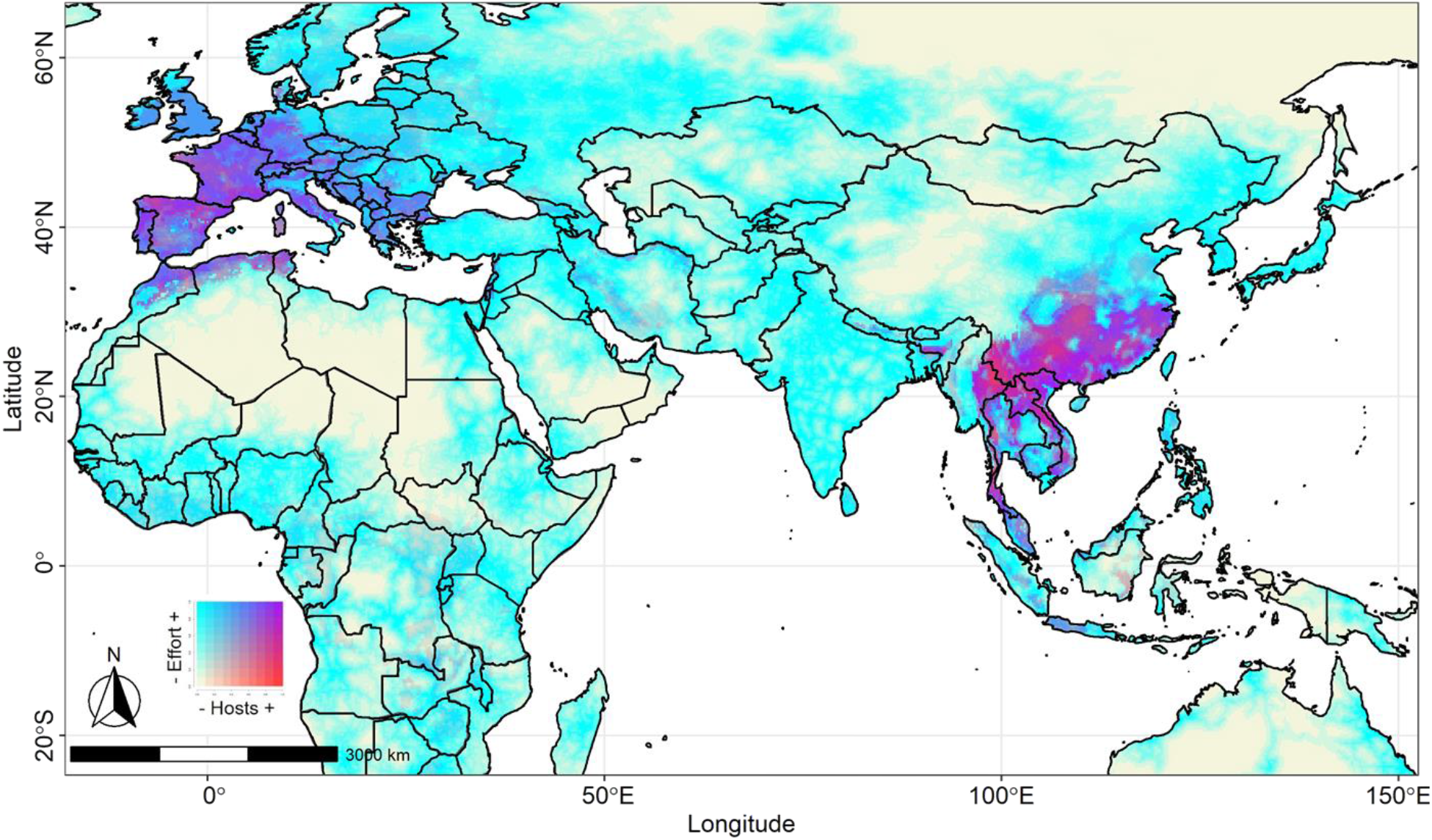
A choropleth bivariate map showing the near current distribution of reported *Sarbecovirus* bat host species and estimated sampling rate calculated for the filtered dataset, according to potential drivers of residual accessibility bias. Areas in red signal high numbers of *Sarbecovirus* hosts, but estimated lower sampling rates driven by accessibility.

Highest host richness values decline in future SSP585 projections (Figure 3). Average temperature of current hotspots - the few where 13 species are present - is 20.6 °C, increasing to 22.7 °C in future hotspots (SSP585, BCC-CSM2-MR in 2100). The hotspot centroid in Southeast Asia is predicted to shift from Myanmar into eastern forests regardless of GCM used. The predicted centroids for highest richness shift from in Kat Ku, Myanmar, to denser forest surroundings to the east, only 42 km apart, considering BCC-CSM2-MR, or more distantly 373 km further east if we consider the CanESM5 GCM (in the east of Ban Ka Kiak, Laos) by 2100 using SSP585. Scenarios project increased species richness in locations (Figure 3) where temperature is also higher.

**Figure 3.**
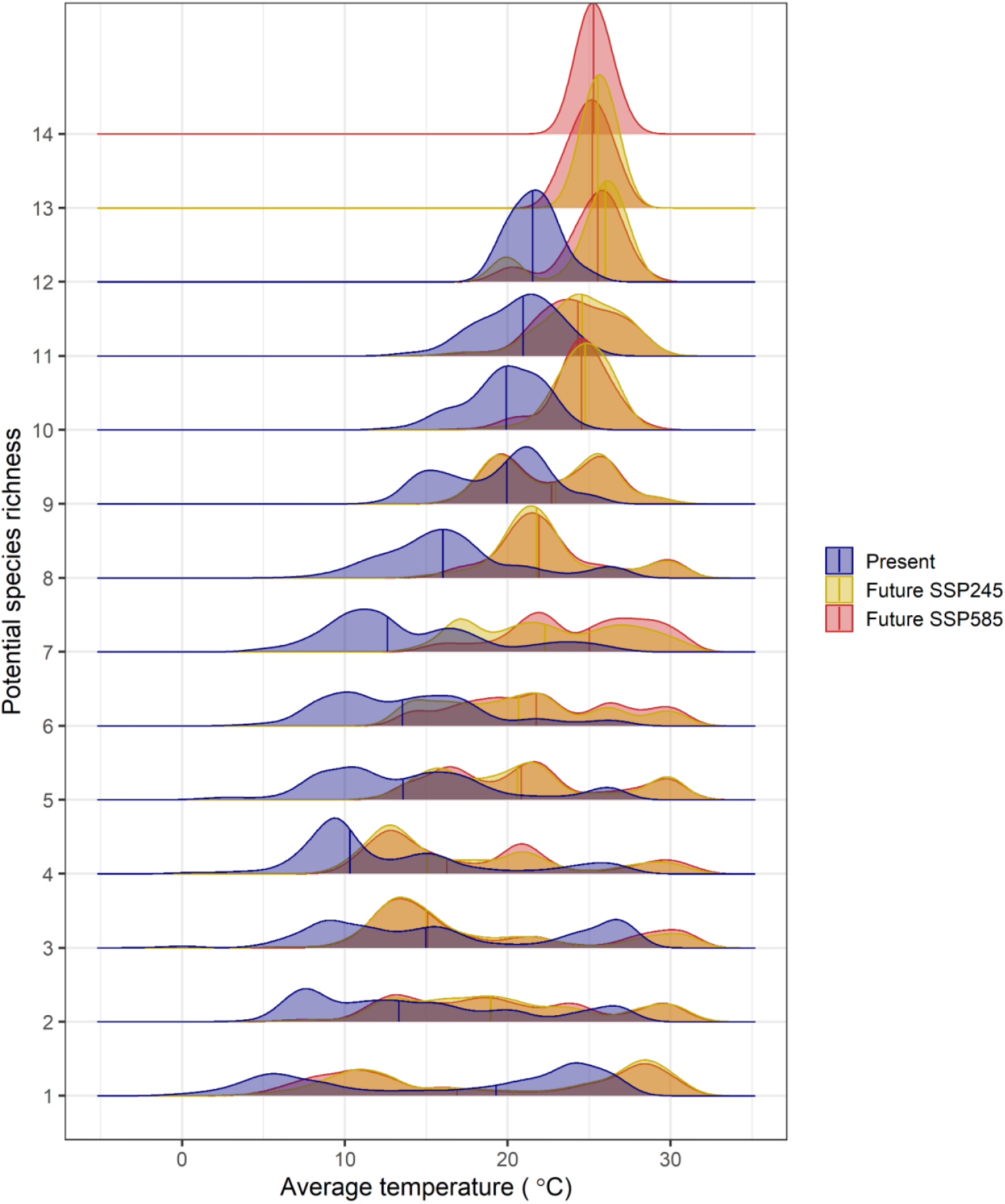
Hotspots of *Sarbecovirus* hosts will be hotter and more concentrated in the future. SSP585 shows a higher maximum of 14 species in the future, whereas SSP245 projects 14 species on only one occasion, considering BCC-CSM2-MR. Vertical lines represent the median.

Potential ranges for all periods, scenarios, and GCMs used are in Table S5. Several species showed resilience, such as *Hipposideros armiger, H. galeritus* and *H. larvatus*. However, some species with large ranges (Figure S12), such as *Rhinolophus ferrumequinum*, and *Rhinolophus affinis*, are predicted to suffer range contractions (Table S5, Table S6). Population trend data showed that many species do not have a current evaluation (Table S7). Considering the most extreme global warming scenario (SSP585), most species will suffer range contractions (N=17, 74%), while six may gain area (N=6), with species overlap decreasing (Figure S13, Table S6). For SSP245, fewer, but still more than half the species will suffer range contraction (N=14, 61%). In terms of total area overlaps by 2100, there is higher overlap in SSP245 in comparison to SSP585 (Figure S14), however it varies with species. Overall, projections show 180.57 million km2 of cumulative area overlap in SSP585 versus 187.26 million km2 of cumulative area overlap in SSP245 considering BCC-CSM2-MR. From species modeled, most (N=16, 70%) agreed in the range shift trend across SSPs. Most differences pointed to range contraction in SSP585, while there was a slight expansion in potential range in SSP245 (Table S5).

## Discussion

Human-driven habitat change—including through global warming—will alter species distributions, and as a consequence species interactions. Species interactions can have far reaching effects in ecological communities. Parasites and infections that animals carry are often overlooked yet key components of ecological systems [55–57] and host distribution changes may redistribute and alter disease emergence risk. Here, we developed SDMs to determine the drivers of *Sarbecovirus* bat host distributions, identify hotspots of host species richness and model changes in distributions and hotspots under future climate scenarios. For the species modelled, temperature seasonality and karst were important determinants of geographic distributions, and we identified host hotspots in Europe and Asia. We projected how these will change under future climate change scenarios, identifying species’ ranges will often contract and fragment and shift hotspots, along with identifying where sampling bias is greatest due to distance from roads.

Our focal species are insectivorous bats with varying geographical ranges and sensitivity to habitat disturbance [58]. Species responses to climate change can be complex [59,60]. Though some species are resilient (Figure S12), *Sarbecovirus* bat hosts are impacted by forest quality and cave area disturbance [3], and our projections highlight their sensitivity to increased temperatures, so we assume forest amount and proximity to karst and rocky areas and climate will be crucial in driving their future distributions. Species differing environment responses reinforce the need to evaluate species-specific differences, even within the same genus. For example, *R. sinicus* lives in montane forests [61], yet *R. affinis* can live in lowland forest, dry forest and disturbed areas [58]. *R. ferrumequinum*, a species ranging from Europe and Northwest Africa to Asia, hibernates during winter in caves, but this varies across the range and with age and sex [58]. Most modelled species are associated with caves (Figure S8), with karst availability varying in importance, but frequently among the most important factors. How karst will change due to mining and land conversion is unclear. Metal mining and limestone quarrying are increasingly threatening karst habitats [62] that bats depend upon [3]. We did not model this potential change, but it may reduce suitable habitat and fragment populations.

Primary remaining forest habitat will likely be refugia for many species. However, the average temperature in the current host diversity hotspots in SE Asia is 20.6 °C, increasing to 22.7 °C under SSP585. With hotspots getting hotter, most sarbecovirus hosts’ ranges will contract in the future, following the expected pattern for Southeast Asian bats [63]. Host diversity hotspots will shift to more climatically stable areas where shrinking remaining primary forests remain (Figure S9). Suitable areas are lost in northern areas, especially on the China borders. These changes reduce species richness with time in both scenarios used.

We chose a pessimistic carbon emissions SSP585 scenario as an example here, but it is a likely scenario [64] and considered a possible future, though less likely according to recent reports [65]. However, there is high convergence between SSP245 and SSP585 hotspot projections (Figure 3), though SSP585 concentrates more species. In fact SSP245 is, to some extent, less extreme with fewer range contractions, whereas species are projected to become more spatially concentrated, especially in SSP585, probably due to a refugia effect [66] since there will be less suitable habitat. We predict slight range gains for 2015-2040, possibly due to forest regrowth, which is more likely if international initiatives for reducing deforestation and nature-based solutions succeed [67,68]. After 2040, in the high carbon emission scenario, we predict suitable habitat loss for most species and a shift in the hotspot centroids from Kat Ku, Myanmar, to denser forest surroundings to the east towards Laos regardless of the global circulation used.

Our analyses highlight the dynamic and uneven nature of data acquisition. Our pipeline can be easily updated to include new or changing data. Ongoing viral discovery will almost certainly add to the list of bat species testing positive for sarbecoviruses, and many bat species, particularly the Rhinolophidae and Hipposideridae in Africa [69–71] are in need of taxonomic revision, and bat species continue to be discovered and described. Revisions could rapidly lead to taxonomic changes and changes in conservation assessments. Similar rapid changes happen with viral surveillance, particularly for bat hosts [72,73]. Also, new species discoveries can indicate or alter sampling bias, since remote areas may be undersampled. More intensive sampling in species-rich effort gap areas can reduce biases. Despite these challenges, after data curation to reduce sampling bias and autocorrelation, we could still model most species but identified important classic shortfalls [74]. The smallest-ranged bats in our dataset did not reach our modelling criteria (Figure S7) because of data gaps, and these gaps are likely larger for *Sarbecovirus* ecology in their natural hosts (from classic Wallacean to Eltonian shortfalls)[15]. Our estimated bat sampling rates were partially driven by accessibility, where effort gaps were mostly correlated with the distance from roads (Figure S10). By providing a surface of estimated sampling rate, we provide a more realistic scenario for prioritizing sampling of the focal species, as uneven sampling is one of the most common violations of assumption in distribution models [75]. Initiatives to prioritize specific bat viral sampling have been based on phylogeny, expert range definition and viral sharing probabilities [10,76]. We suggest areas with high estimated diversity of hosts and low estimated sampling rates should be a priority for *Sarbecovirus* host studies in the future, such as Indochina and China hotspots (areas highlighted in red, Figure 2) [77].

There are conservation implications of our findings. Range contractions are predicted for several species, even for species using variable habitats, such as *Rhinolophus pearsonii* (Table S4, Figure S5). Most species’ populations are currently declining (N=8) or have unknown population trends (N=20, Table S6). Local species loss predictions were almost twice the number of maximum gain values (Figure S9). Along with climate change mitigation, strategies for maintaining landscape-level habitat connectivity will allow populations to reach refugia and lower extinction risk. This could be done by developing landscape connectivity surfaces that maximize diversity hotspot extensions, with monitoring effective dispersal through genetic [78,79] and population [80].

Our results identify broad regions where bats reported positive for sarbecoviruses most probably occur and co-occur. These hotspots coincide, but are not restricted only to Rhinolophidae diversity hotspots previously reported [81] and to hotspots of mammal vulnerability to climate change [82]. Projections suggest that hundreds of new future viral sharing events may occur in Southeast Asia [7]. Novel interactions may be of concern for species survival as pathogens could spread more easily in vulnerable wild populations, which could facilitate epizootics and panzootics [83]. The role of bats as putative reservoirs of different zoonosis-causing agents must be interpreted with care, though [84]. Sarbecoviruses circulating in horseshoe bats might be directly infectious for humans [85,86], or infect other species prior to people [17,87]. Thus, bat presences may act as one component of hazard in risk assessments using ecosystem perspectives and multiple drivers [88]. Increased incidence of zoonoses is more likely through human-mediated change of the environment, including climate [89]. Changes in host hotspots may alter disease risk when other changes to human, intermediate domestic or wildlife populations take place [12].

Our future projections assume models using present data will perform adequately. However, our models do not account for biotic components that also interfere with suitability, so we are limited to inferences of distribution derived from landscape and climate drivers. Projections will therefore need validation with new data and new predictions. Related, small changes may be relevant for local health and conservation initiatives, and coronavirus hosts are a focus of increasing research [77]. Studies represent spatio-temporal snapshots of nature: data change as new hosts are identified [10], host distributions revised, and remote sensing of their drivers updated. We provide a pipeline ready for the inevitable addition of new bat hosts (e.g. [9,73]), which could also be applied for inferring the distribution of potential intermediate hosts for sarbecoviruses. As we affect climate, biodiversity and ecosystems in real-time, we can validate and update our predictions (forecasting). Beyond refining distributional ecology, more work on host characterization will improve our understanding of the role of bats as reservoirs of coronaviruses [77]. Here, we could predict range contractions for most species of bats hosts of sarbecoviruses in response to global changes in climate and forest cover, along with host hotspot shifts. Further evaluations will help inform climate change vulnerability assessments [90] and future integrative data-modeling steps, in addition to communication of processes involving bats that could benefit One Health [91] and Nature-based solutions projects.

## Acknowledgments

Massey University subscription to New Zealand eScience Infrastructure (NeSi, #03262), Te Pūnaha Matatini (Mahia Te Mahi workshop, RLM), Dan Becker and the Verena Consortium (viralemergence.org) for database curation and Paolo D’Odorico for comments.

## References

1. IPBES. 2020 Workshop Report on Biodiversity and Pandemics of the Intergovernmental Platform on Biodiversity and Ecosystem Services.

2. Allen T, Murray KA, Zambrana-Torrelio C, Morse SS, Rondinini C, Di Marco M, Breit N, Olival KJ, Daszak P. 2017 Global hotspots and correlates of emerging zoonotic diseases. Nat. Commun. 8, 1124.

3. Frick WF, Kingston T, Flanders J. 2020 A review of the major threats and challenges to global bat conservation. Ann. N. Y. Acad. Sci. 1469, 5–25.

4. Olival KJ, Hosseini PR, Zambrana-Torrelio C, Ross N, Bogich TL, Daszak P. 2017 Host and viral traits predict zoonotic spillover from mammals. Nature 546, 646–650.

5. Carlson CJ et al. 2021 The future of zoonotic risk prediction. Philos. Trans. R. Soc. Lond. B Biol. Sci. (doi:10.1098/rstb.2020.0358)

6. Phelps KL, Hamel L, Alhmoud N, Ali S, Bilgin R, Sidamonidze K, Urushadze L, Karesh W, Olival KJ. 2019 Bat Research Networks and Viral Surveillance: Gaps and Opportunities in Western Asia. Viruses 11. (doi:10.3390/v11030240)

7. Carlson CJ, Albery GF, Merow C, Trisos CH, Zipfel CM, Eskew EA, Olival KJ, Ross N, Bansal S. 2021 Climate change will drive novel cross-species viral transmission. bioRxiv., 2020.01.24.918755. (doi:10.1101/2020.01.24.918755)

8. Zhou P et al. 2020 A pneumonia outbreak associated with a new coronavirus of probable bat origin. Nature. 579, 270–273. (doi:10.1038/s41586-020-2012-7)

9. Temmam S et al. 2021 Coronaviruses with a SARS-CoV-2-like receptor-binding domain allowing ACE2-mediated entry into human cells isolated from bats of Indochinese peninsula. (doi:10.21203/rs.3.rs-871965/v1)

10. Becker DJ et al. 2020 Predicting wildlife hosts of betacoronaviruses for SARS-CoV-2 sampling prioritization. bioRxiv., 2020.05.22.111344. (doi:10.1101/2020.05.22.111344)

11. Rulli MC, D’Odorico P, Galli N, Hayman DTS. 2021 Land-use change and the livestock revolution increase the risk of zoonotic coronavirus transmission from rhinolophid bats. Nature Food 2, 409–416.

12. Sánchez CA, Li H, Phelps KL, Zambrana-Torrelio C, Wang L-F, Olival KJ, Daszak P. 2021 A strategy to assess spillover risk of bat SARS-related coronaviruses in Southeast Asia. medRxiv, 2021.09.09.21263359.

13. Simmons NB, Cirranello AL. 2021 Bat Species of the World: A taxonomic and geographic database. Bat Species of the World: A taxonomic and geographic database. See https://batnames.org/ (accessed on 17 August 2021).

14. Hayman DTS. 2016 Bats as Viral Reservoirs. Annual Review of Virology. 3, 77–99. (doi:10.1146/annurev-virology-110615-042203)

15. Letko M, Seifert SN, Olival KJ, Plowright RK, Munster VJ. 2020 Bat-borne virus diversity, spillover and emergence. Nat. Rev. Microbiol. 18, 461–471.

16. Andersen KG, Rambaut A, Lipkin WI, Holmes EC, Garry RF. 2020 The proximal origin of SARS-CoV-2. Nat. Med. 26, 450–452.

17. Boni MF, Lemey P, Jiang X, Lam TT-Y, Perry BW, Castoe TA, Rambaut A, Robertson DL. 2020 Evolutionary origins of the SARS-CoV-2 sarbecovirus lineage responsible for the COVID-19 pandemic. Nature Microbiology 5, 1408–1417.

18. Kunz TH, de Torrez EB, Bauer D, Lobova T, Fleming TH. 2011 Ecosystem services provided by bats. Ann. N. Y. Acad. Sci. 1223, 1–38.

19. Wilkinson DA, Marshall JC, French NP, Hayman DTS. 2018 Habitat fragmentation, biodiversity loss and the risk of novel infectious disease emergence. J. R. Soc. Interface 15. (doi:10.1098/rsif.2018.0403)

20. Tanalgo KC, Tabora JAG, Ac. H. 2018 Bat cave vulnerability index (BCVI): A holistic rapid assessment tool to identify priorities for effective cave conservation in the tropics. Ecol. Indic. 89, 852–860.

21. Furey NM, Racey PA. 2016 Conservation Ecology of Cave Bats. In Bats in the Anthropocene: Conservation of Bats in a Changing World, pp. 463–500. Springer, Cham.

22. Tanalgo KC et al. 2021 DarkCideS 1.0, a global database for bats in karsts and caves. Authorea Preprints (doi:10.22541/au.163578759.92395202/v1)

23. Kingston T. 2010 Research priorities for bat conservation in Southeast Asia: a consensus approach. Biodivers. Conserv. 19, 471–484.

24. Streicker DG et al. 2016 Host–pathogen evolutionary signatures reveal dynamics and future invasions of vampire bat rabies. Proc. Natl. Acad. Sci. U. S. A. 113, 10926–10931.

25. Cui J et al. 2007 Evolutionary Relationships between Bat Coronaviruses and Their Hosts. Emerg. Infect. Dis. 13, 1526.

26. Willoughby AR, Phelps KL, PREDICT Consortium, Olival KJ. 2017 A Comparative Analysis of Viral Richness and Viral Sharing in Cave-Roosting Bats. Diversity 9, 35.

27. Zhou H et al. 2021 Identification of novel bat coronaviruses sheds light on the evolutionary origins of SARS-CoV-2 and related viruses. Cell 184, 4380–4391.e14.

28. Kitratporn N, Takeuchi W. 2019 Spatiotemporal Distribution of Human–Elephant Conflict in Eastern Thailand: A Model-Based Assessment Using News Reports and Remotely Sensed Data. Remote Sensing 12, 90.

29. Latinne A et al. 2020 Origin and cross-species transmission of bat coronaviruses in China. Nat. Commun. 11, 1–15.

30. Lin X, Wang W, Hao Z, Wang Z, Guo W, Guan Q. 2017 Extensive diversity of coronaviruses in bats from China. Virology 507, 1–10.

31. R Core Team. 2020 R: A Language and Environment for Statistical Computing.

32. Carlson CJ et al. 2021 The Global Virome in One Network (VIRION): an atlas of vertebrate-virus associations. bioRxiv., 2021.08.06.455442. (doi:10.1101/2021.08.06.455442)

33. Luo J, Jiang T, Lu G, Wang L, Wang J, Feng J. 2013 Bat conservation in China: should protection of subterranean habitats be a priorityã Oryx 47, 526–531.

34. Hughes AC, Orr MC, Ma K, Costello MJ, Waller J, Provoost P, Yang Q, Zhu C, Qiao H. 2021 Sampling biases shape our view of the natural world. Ecography

35. Fisher-Phelps M, Cao G, Wilson RM, Kingston T. 2017 Protecting bias: Across time and ecology, open-source bat locality data are heavily biased by distance to protected area. Ecol. Inform. 40, 22–34.

36. Zizka A et al. 2019 CoordinateCleaner : Standardized cleaning of occurrence records from biological collection databases. Methods in Ecology and Evolution. 10, 744–751. (doi:10.1111/2041-210x.13152)

37. Wilson D, Mittermeier R, editors. 2019 Handbook of the Mammals of the World. Barcelona: Springer.

38. Fuwen WEI et al. 2021 Catalogue of mammals in China(2021). Shou Lei Xue Bao 41, 487.

39. In press. ENMTML: An R package for a straightforward construction of complex ecological niche models. See https://paperpile.com/app/p/eab49d31-560a-0e13-b8f5-5ff92def0cf7 (accessed on 7 October 2021).

40. Varela S, Anderson RP, García-Valdés R, Fernández-González F. 2014 Environmental filters reduce the effects of sampling bias and improve predictions of ecological niche models. Ecography., no–no. (doi:10.1111/j.1600-0587.2013.00441.x)

41. Proosdij ASJ, Sosef MSM, Wieringa JJ, Raes N. 2016 Minimum required number of specimen records to develop accurate species distribution models. Ecography. 39, 542–552. (doi:10.1111/ecog.01509)

42. Zizka A, Antonelli A, Silvestro D. 2021 sampbias, a method for quantifying geographic sampling biases in species distribution data. Ecography. 44, 25–32. (doi:10.1111/ecog.05102)

43. Olson DM et al. 2001 Terrestrial Ecoregions of the World: A New Map of Life on Earth: A new global map of terrestrial ecoregions provides an innovative tool for conserving biodiversity. Bioscience 51, 933–938.

44. Lobo JM, Jimenez-Valverde A, Hortal J. 2010 The uncertain nature of absences and their importance in species distribution modelling. Ecography 33, 103–114.

45. Mendes P, Velazco SJE, de Andrade AFA, de Marco P. 2020 Dealing with overprediction in species distribution models: How adding distance constraints can improve model accuracy. Ecol. Modell. 431, 109180.

46. Vale MM, Lima-ribeiro MS, Rocha C. 2021 View of Global land-use and land-cover data for ecologists: Historical, current, and future scenarios. Biodiversity Informatics, 28–38.

47. Fick SE, Hijmans RJ. 2017 WorldClim 2: new 1-km spatial resolution climate surfaces for global land areas. International Journal of Climatology. 37, 4302–4315. (doi:10.1002/joc.5086)

48. Dormann CF et al. 2013 Collinearity: a review of methods to deal with it and a simulation study evaluating their performance. Ecography. 36, 27–46. (doi:10.1111/j.1600-0587.2012.07348.x)

49. Qiao H, Soberón J, Peterson AT. 2015 No silver bullets in correlative ecological niche modelling: Insights from testing among many potential algorithms for niche estimation. Methods Ecol. Evol. 6, 1126–1136.

50. Duan R-Y, Kong X-Q, Huang M-Y, Fan W-Y, Wang Z-G. 2014 The Predictive Performance and Stability of Six Species Distribution Models. PLoS One 9. (doi:10.1371/journal.pone.0112764)

51. Boyce MS, Vernier PR, Nielsen SE, Schmiegelow FKA. 2002 Evaluating resource selection functions. Ecol. Modell. 157, 281–300.

52. Lawson CR, Hodgson JA, Wilson RJ, Richards SA. 2014 Prevalence, thresholds and the performance of presence-absence models. Methods Ecol. Evol. 5, 54–64.

53. Liu C, White M, Newell G. 2013 Selecting thresholds for the prediction of species occurrence with presence-only data. J. Biogeogr. 40, 778–789.

54. In press. Climate Change 2021: The Physical Science Basis. See https://www.ipcc.ch/report/sixth-assessment-report-working-group-i/ (accessed on 22 September 2021).

55. Rigaud T, Perrot-Minnot M-J, Brown MJF. 2010 Parasite and host assemblages: embracing the reality will improve our knowledge of parasite transmission and virulence. Proceedings of the Royal Society B: Biological Sciences (doi:10.1098/rspb.2010.1163)

56. Albery GF et al. 2021 The science of the host–virus network. Nature Microbiology 6, 1483–1492.

57. Lafferty KD et al. 2008 Parasites in food webs: the ultimate missing links. Ecol. Lett. 11. (doi:10.1111/j.1461-0248.2008.01174.x)

58. Hutson AM, Mickleburgh SP, Racey PA. 2001 Microchiropteran bats: global status survey and conservation action plan. Switzerland and Cambridge, UK: IUCN, Gland.

59. Chen I-C, Hill JK, Ohlemüller R, Roy DB, Thomas CD. 2011 Rapid Range Shifts of Species Associated with High Levels of Climate Warming. Science (doi:10.1126/science.1206432)

60. Di Marco M, Pacifici M, Maiorano L, Rondinini C. 2021 Drivers of change in the realised climatic niche of terrestrial mammals. Ecography 44, 1180–1190.

61. Luo X, Liang L, Liu Z, Wang J, Huang T, Geng D, Chen B. 2020 Habitat Suitability Evaluation of the Chinese Horseshoe Bat (R. sinicus) in the Wuling Mountain Area Based on MAXENT Modelling. Pol. J. Environ. Stud. 29, 1263–1273.

62. 2021 Surge in global metal mining threatens vulnerable ecosystems. Glob. Environ. Change 69, 102303.

63. Hughes AC, Satasook C, Bates PJJ, Bumrungsri S, Jones G. 2012 The projected effects of climatic and vegetation changes on the distribution and diversity of Southeast Asian bats. Glob. Chang. Biol. 18, 1854–1865.

64. Schwalm CR, Glendon S, Duffy PB. 2020 RCP8.5 tracks cumulative CO2emissions. Proceedings of the National Academy of Sciences. 117, 19656–19657. (doi:10.1073/pnas.2007117117)

65. Hausfather Z, Peters GP. 2020 Emissions – the ‘business as usual’ story is misleading. Nature 577, 618–620.

66. Morelli TL et al. 2016 Managing Climate Change Refugia for Climate Adaptation. PLoS One 11, e0159909.

67. Estoque RC, Ooba M, Avitabile V, Hijioka Y, DasGupta R, Togawa T, Murayama Y. 2019 The future of Southeast Asia’s forests. Nat. Commun. 10, 1829.

68. Lechner AM et al. 2020 Challenges and considerations of applying nature-based solutions in low- and middle-income countries in Southeast and East Asia. Blue-Green Systems 2, 331–351.

69. Ith S et al. 2016 Geographical Variation of Rhinolophus affinis (Chiroptera: Rhinolophidae) in the Sundaic Subregion of Southeast Asia, including the Malay Peninsula, Borneo and Sumatra. acta 18, 141–161.

70. Demos TC, Webala PW, Goodman SM, Kerbis Peterhans JC, Bartonjo M, Patterson BD. 2019 Molecular phylogenetics of the African horseshoe bats (Chiroptera: Rhinolophidae): expanded geographic and taxonomic sampling of the Afrotropics. BMC Evol. Biol. 19, 1–14.

71. Vallo P, Guillén-Servent A, Benda P, Pires DB, Koubek P. 2008 Variation of mitochondrial DNA in the Hipposideros caffer complex (Chiroptera: Hipposideridae) and its taxonomic implications. acta 10, 193–206.

72. Gibb R, Albery GF, Mollentze N, Eskew EA, Brierley L, Ryan SJ, Seifert SN, Carlson CJ. 2021 Mammal virus diversity estimates are unstable due to accelerating discovery effort. bioRxiv., 2021.08.10.455791. (doi:10.1101/2021.08.10.455791)

73. Wu Z et al. 2021 A comprehensive survey of bat sarbecoviruses across China for the origin tracing of SARS-CoV and SARS-CoV-2. Research Square. (doi:10.21203/rs.3.rs-885194/v1)

74. Hortal J, de Bello F, Diniz-Filho JAF, Lewinsohn TM, Lobo JM, Ladle RJ. 2015 Seven Shortfalls that Beset Large-Scale Knowledge of Biodiversity. Annu. Rev. Ecol. Evol. Syst. 46, 523–549.

75. 2021 Want to model a species nicheã A step-by-step guideline on correlative ecological niche modelling. Ecol. Modell. 456, 109671.

76. Grange ZL et al. 2021 Ranking the risk of animal-to-human spillover for newly discovered viruses. Proc. Natl. Acad. Sci. U. S. A. 118. (doi:10.1073/pnas.2002324118)

77. Ruiz-Aravena M et al. 2021 Ecology, evolution and spillover of coronaviruses from bats. Nat. Rev. Microbiol., 1–16.

78. Baguette M, Blanchet S, Legrand D, Stevens VM, Turlure C. 2013 Individual dispersal, landscape connectivity and ecological networks. Biol. Rev. Camb. Philos. Soc. 88, 310–326.

79. 2018 Landscape resistance influences effective dispersal of endangered golden lion tamarins within the Atlantic Forest. Biol. Conserv. 224, 178–187.

80. Finch D, Corbacho DP, Schofield H, Davison S, Wright PGR, Broughton RK, Mathews F. 2020 Modelling the functional connectivity of landscapes for greater horseshoe bats Rhinolophus ferrumequinum at a local scale. Landsc. Ecol. 35, 577–589.

81. Wang Q, Chen H, Shi Y, Hughes AC, Liu WJ, Jiang J, Gao GF, Xue Y, Tong Y. 2021 Tracing the origins of SARS-CoV-2: lessons learned from the past. Cell Res. 31, 1139–1141.

82. Pacifici M, Visconti P, Rondinini C. 2018 A framework for the identification of hotspots of climate change risk for mammals. Glob. Chang. Biol. 24, 1626–1636.

83. Sweeny AR, Albery GF, Becker DJ, Eskew EA, Carlson CJ. 2021 Synzootics. J. Anim. Ecol. (doi:10.1111/1365-2656.13595)

84. Shapiro JT et al. 2021 Setting the Terms for Zoonotic Diseases: Effective Communication for Research, Conservation, and Public Policy. Viruses 13. (doi:10.3390/v13071356)

85. Wang N et al. 2018 Serological Evidence of Bat SARS-Related Coronavirus Infection in Humans, China. Virol. Sin. 33, 104.

86. Zheng BJ, Guan Y, Wong KH, Zhou J, Wong KL, Young BWY, Lu LW, Lee SS. 2004 SARS-related Virus Predating SARS Outbreak, Hong Kong. Emerg. Infect. Dis. 10, 176.

87. WHO. 2021 WHO-convened global study of origins of SARS-CoV-2: China Part. See https://www.who.int/publications/i/item/who-convened-global-study-of-origins-of-sars-cov-2-china-part (accessed on 22 September 2021).

88. Gibb R, Franklinos LHV, Redding DW, Jones KE. 2020 Ecosystem perspectives are needed to manage zoonotic risks in a changing climate. BMJ 371. (doi:10.1136/bmj.m3389)

89. Lu M, Wang X, Ye H, Wang H, Qiu S, Zhang H, Liu Y, Luo J, Feng J. 2021 Does public fear that bats spread COVID-19 jeopardize bat conservationã Biol. Conserv. 254, 108952.

90. Santos MJ, Smith AB, Dekker SC, Eppinga MB, Leitão PJ, Moreno-Mateos D, Morueta-Holme N, Ruggeri M. 2021 The role of land use and land cover change in climate change vulnerability assessments of biodiversity: a systematic review. Landsc. Ecol. (doi:10.1007/s10980-021-01276-w)

91. Decker DJ, Evensen DTN, Siemer WF, Leong KM, Riley SJ, Wild MA, Castle KT, Higgins CL. 2010 Understanding Risk Perceptions to Enhance Communication about Human-Wildlife Interactions and the Impacts of Zoonotic Disease. ILAR J. 51, 255–261.

