## Supplements for "Present and future distribution of bat hosts of sarbecoviruses: implications for conservation and public health": supplemental_material_biorxv.pdf

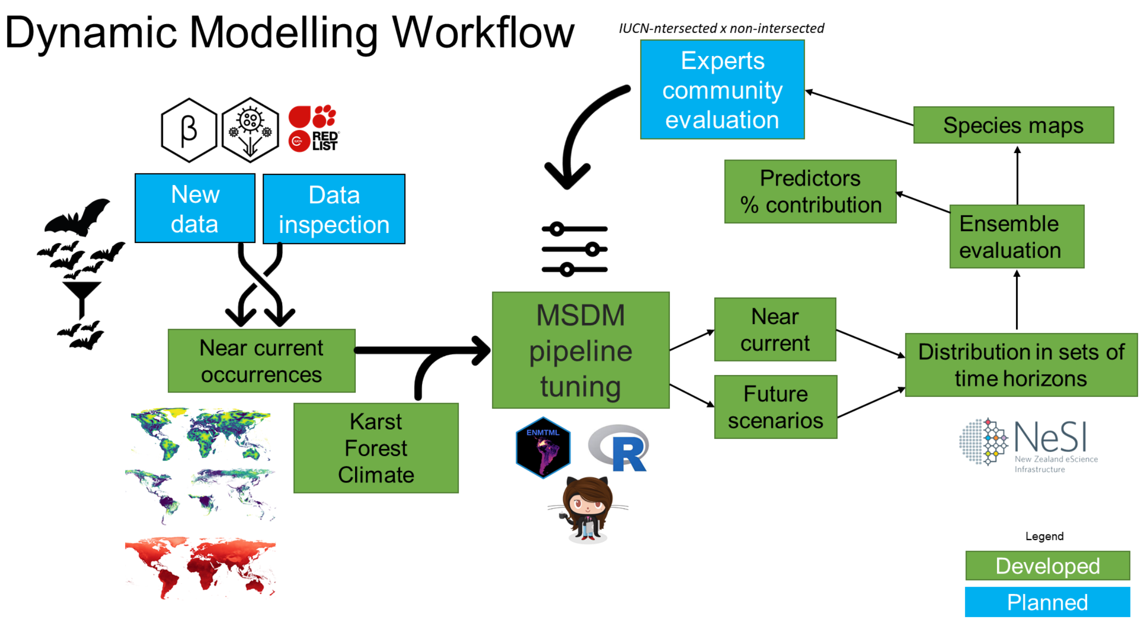


**Figure S1.** Dynamic modelling workflow for species distribution models developed here based on open science and open data initiatives.

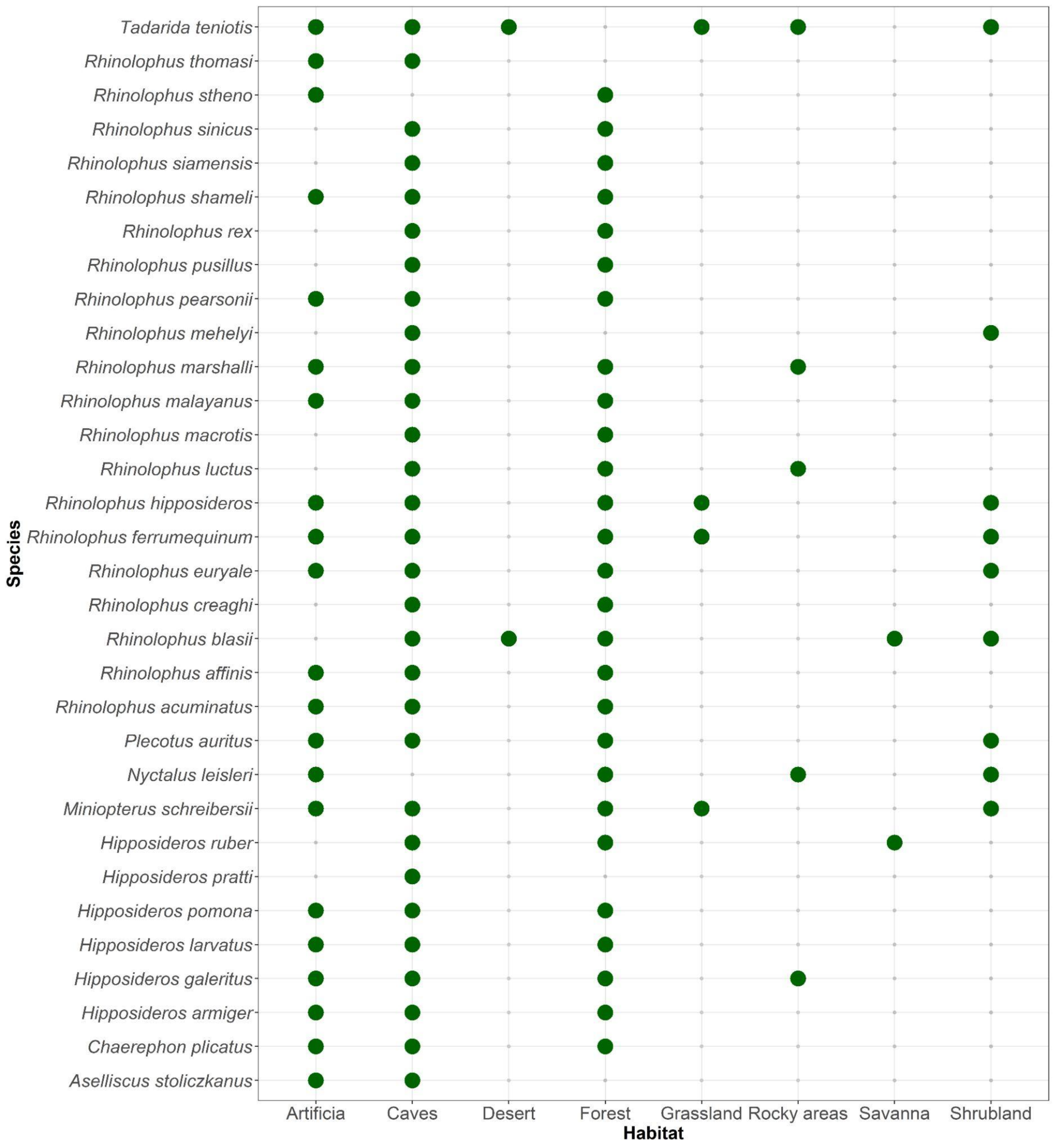


**Figure S2.** Occupied habitats per species as listed in the IUCN Red List Assessments for each species (IUCN 2021).

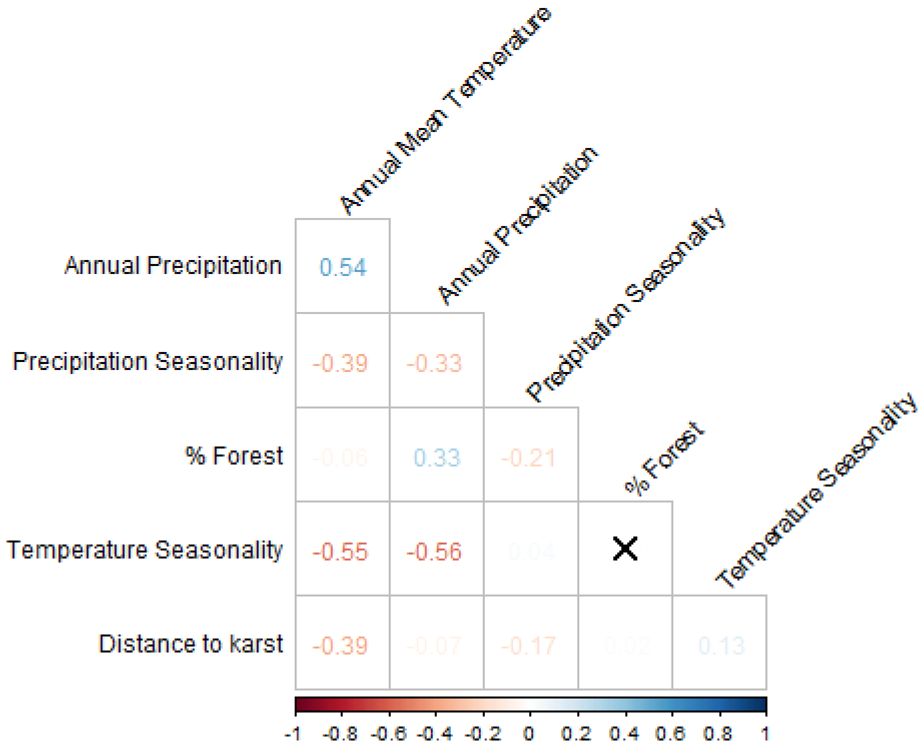


**Figure S3.** Correlation plot of the selected covariates used in species distribution models for sarbecovirus bat hosts. We considered those covariates as having low correlation among each other as they all presented absolute correlation values lower than 0.7. Black cross means non-significant correlation from a Pearson's correlation test.

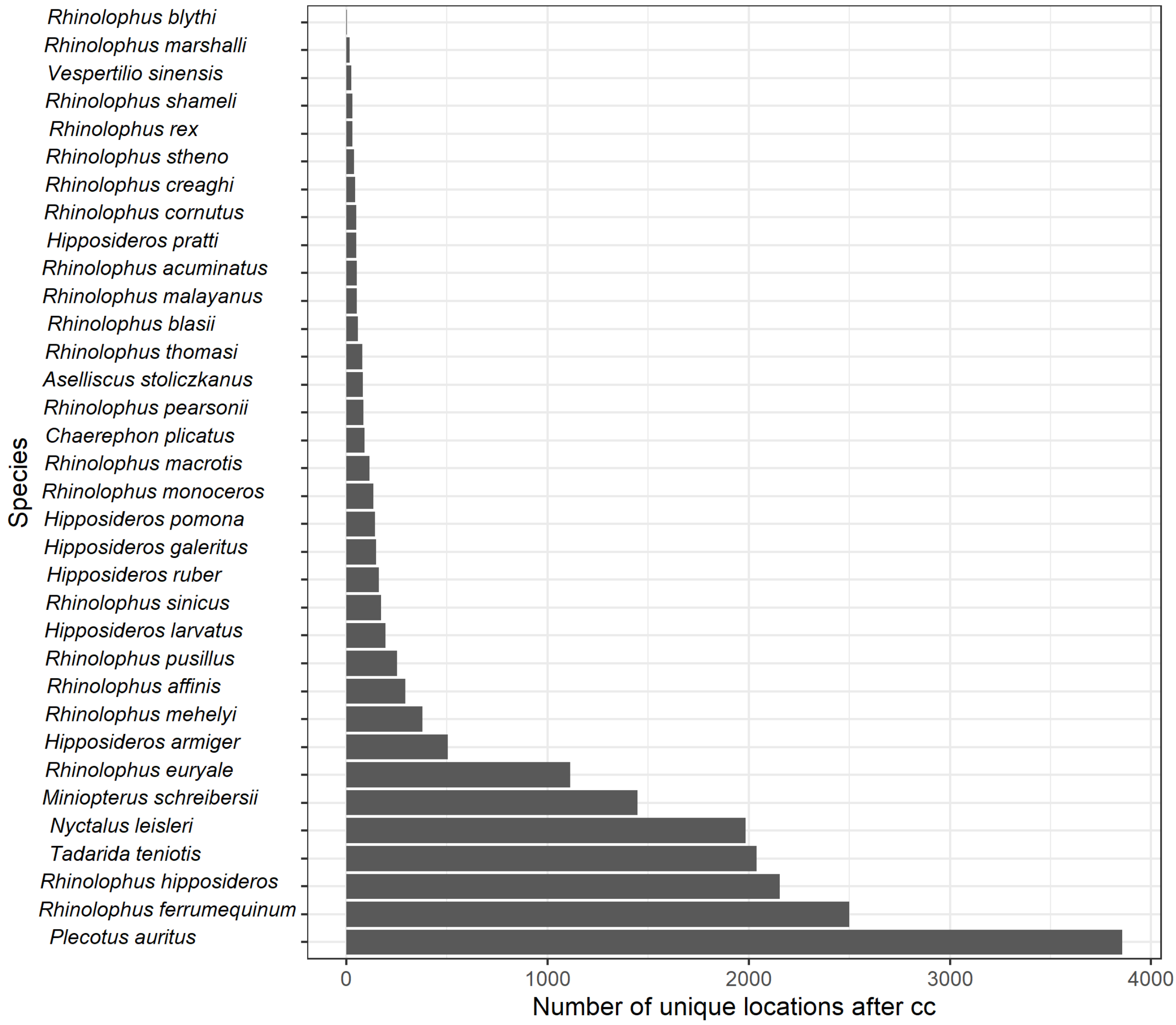


**Figure S4.** Number of unique occurrences of bat hosts of Sarbecoviruses after the cleaning process prior to thinning and prior to intersecting with IUCN polygons. In the x axes, ‘cc’ means 'after the coordinate cleaning process'.

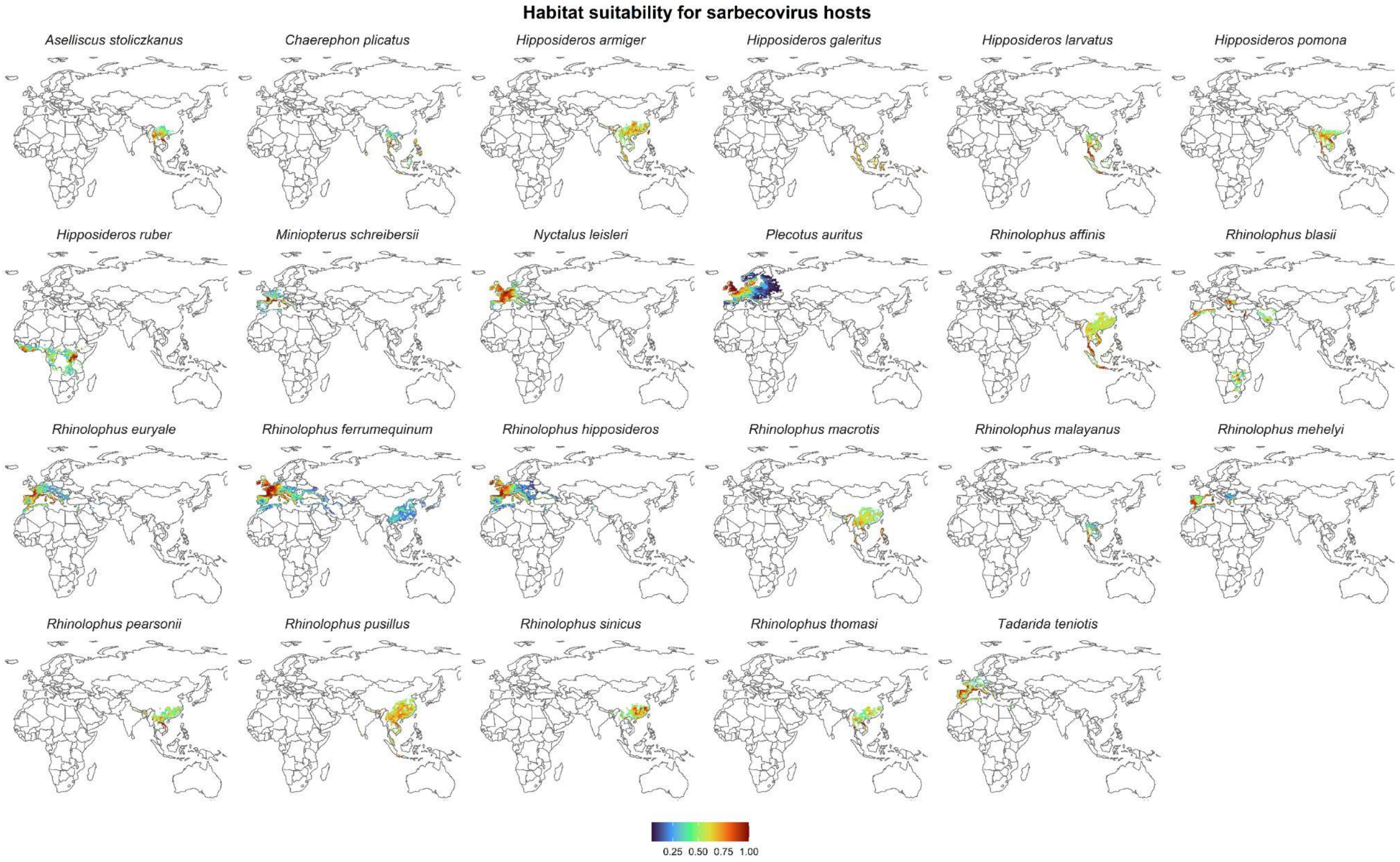


**Figure S5.** Habitat suitability maps for non-IUCN intersected data.

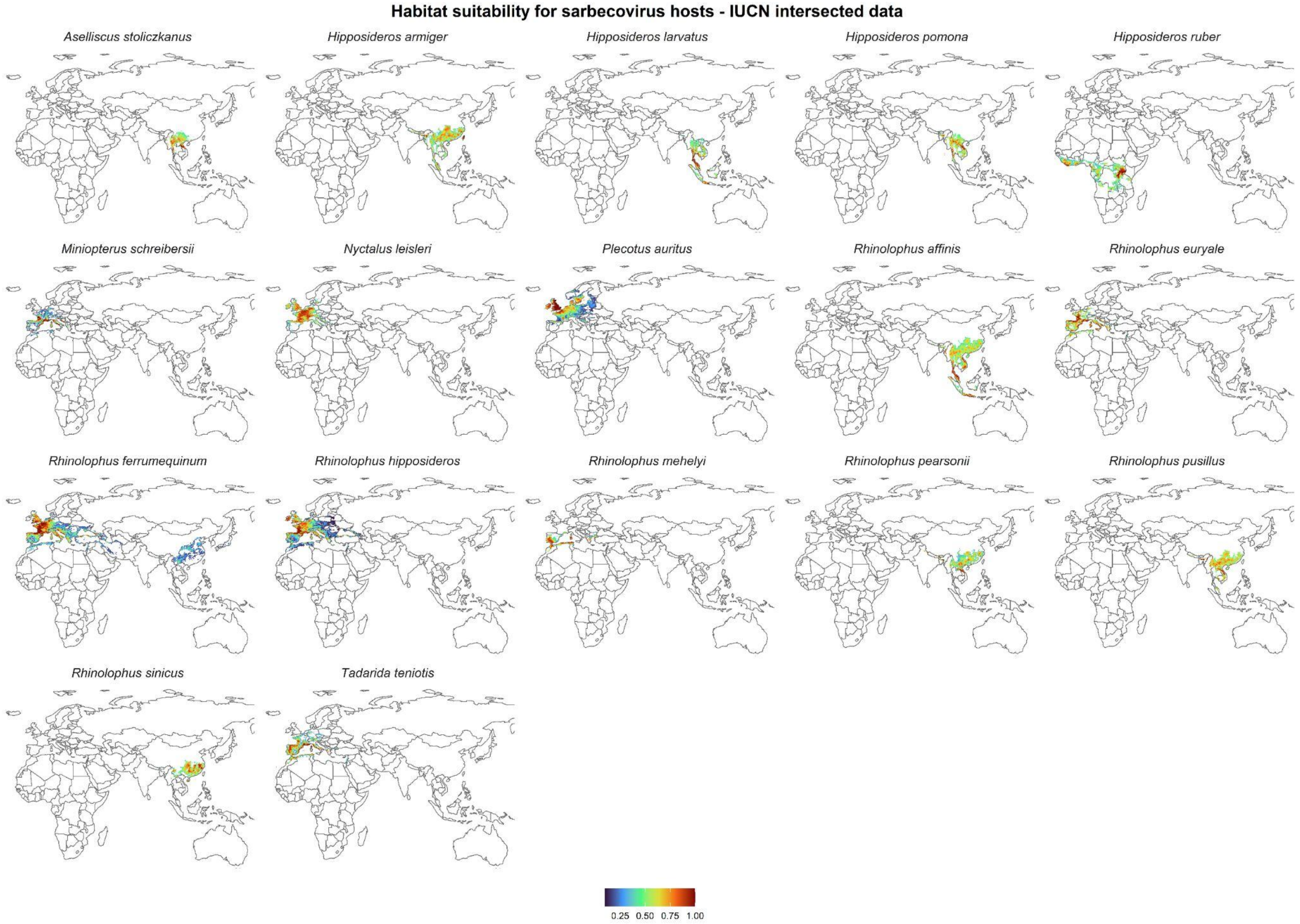


**Figure S6.** Habitat suitability maps for IUCN-intersected data.

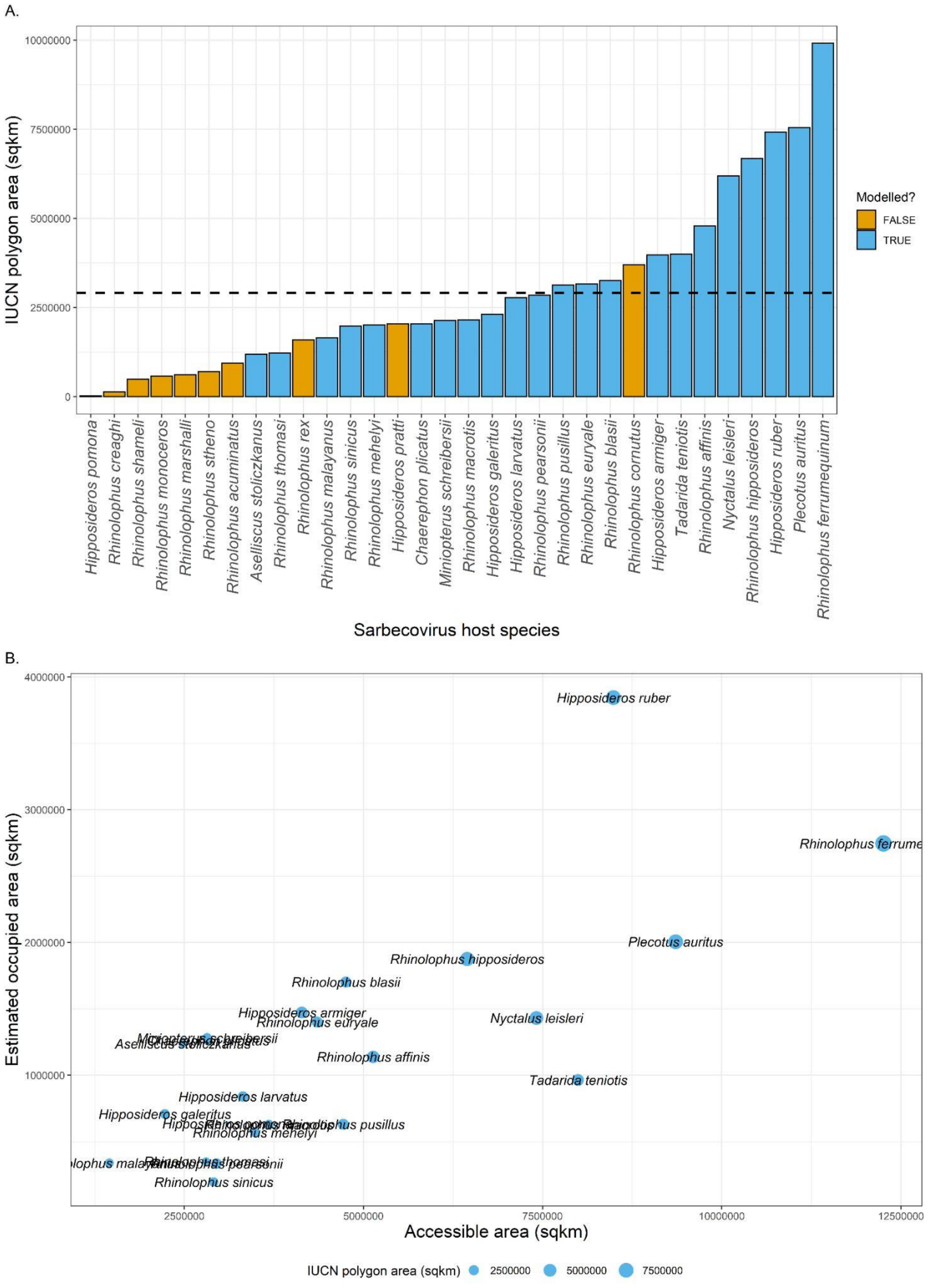


**Figure S7.** A. Distributions of species with smaller expected ranges (IUCN polygons) are more frequently challenging to model due to fewer data points. The dashed black line displays the average range from IUCN polygons. Spatially restricted modelled range. B. Accessible area is defined as the Olson bioregions where species occurred and their estimated occurrence area.

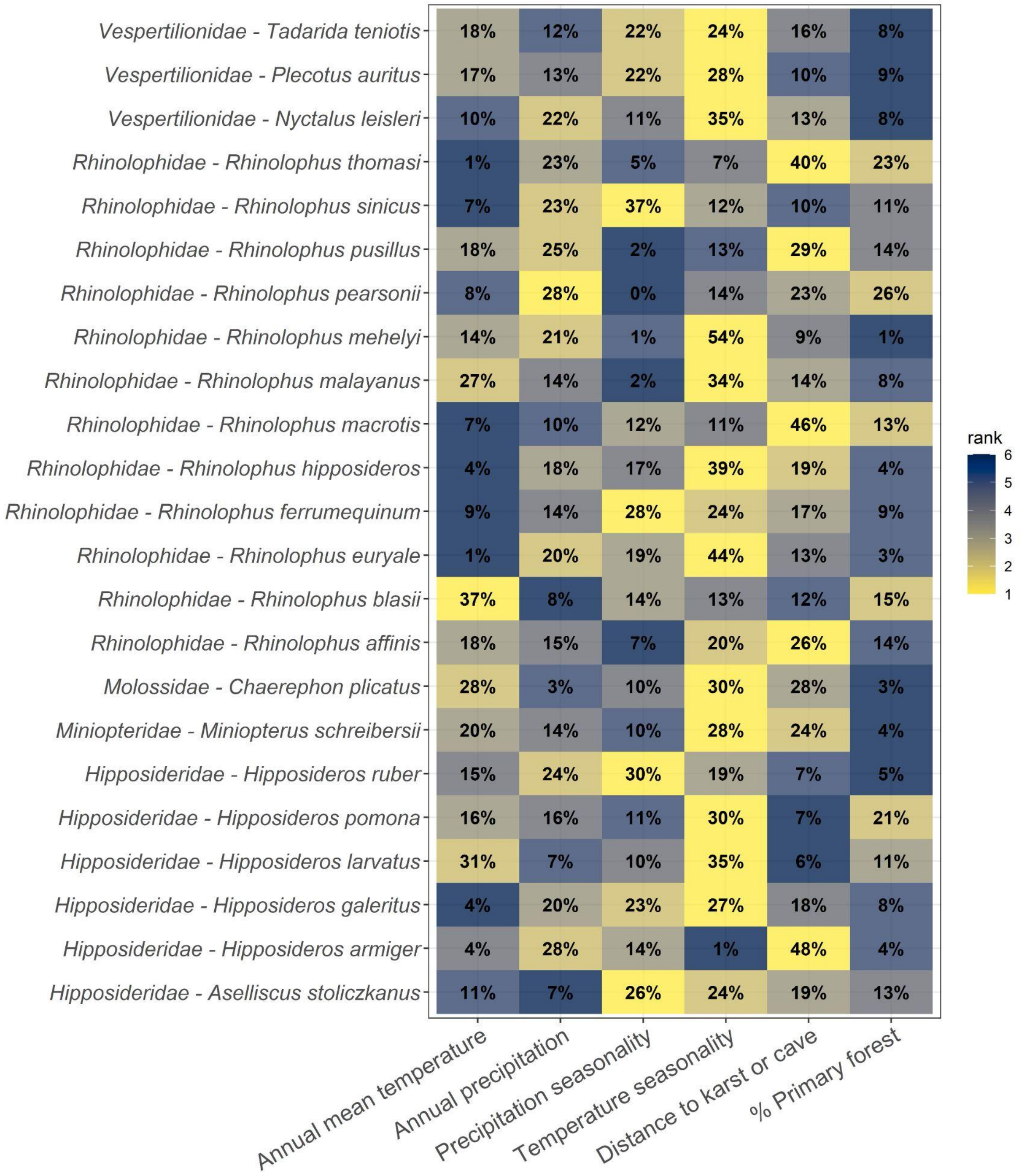


**Figure S8.** A Variable importance (rounded proportion for zero decimals and rank for comparison) for bat species modelled.

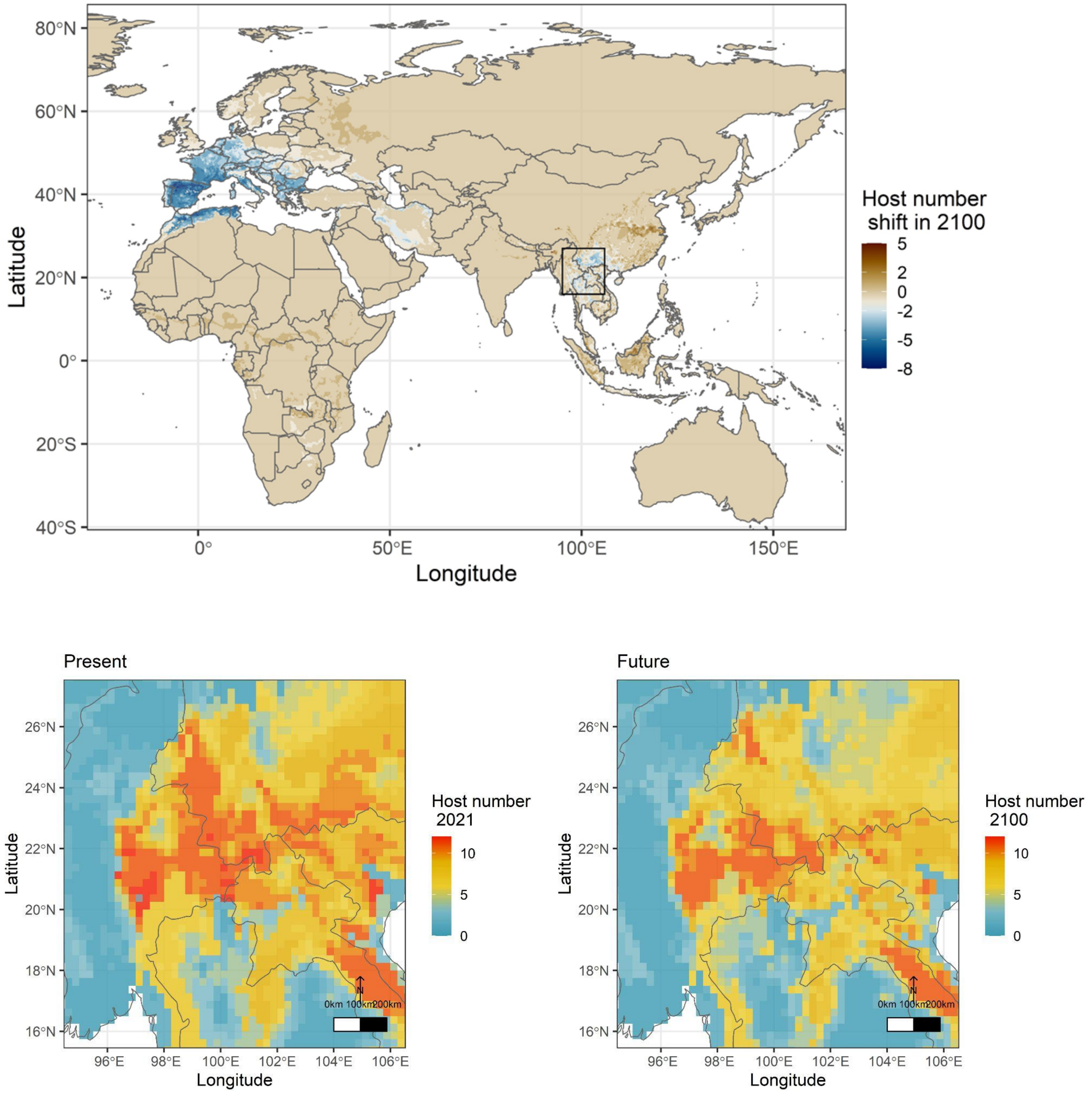


**Figure S9.** Species shifts considering SSP585 in 2100 with insets of the hotspots of Sarbecovirus bat hosts. Hotspots are more fragmented in the future (SSP585, BCC-CSM2-MR).

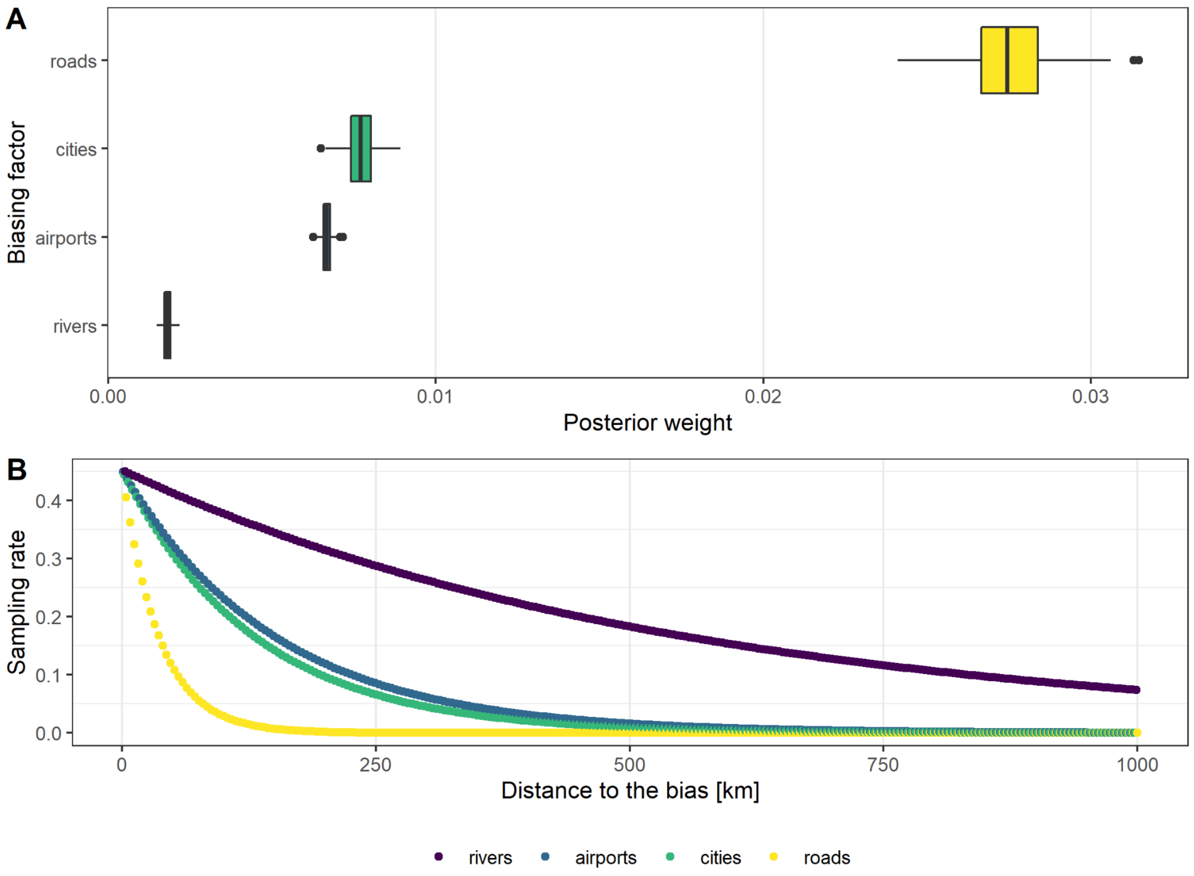


**Figure S10.** Residual accessibility bias on sarbecovirus hosts considering roads, cities, airports and rivers. A. Biasing factor posterior weight per bias type. B. Estimated sampling rate decay as a distance to bias source.

C.

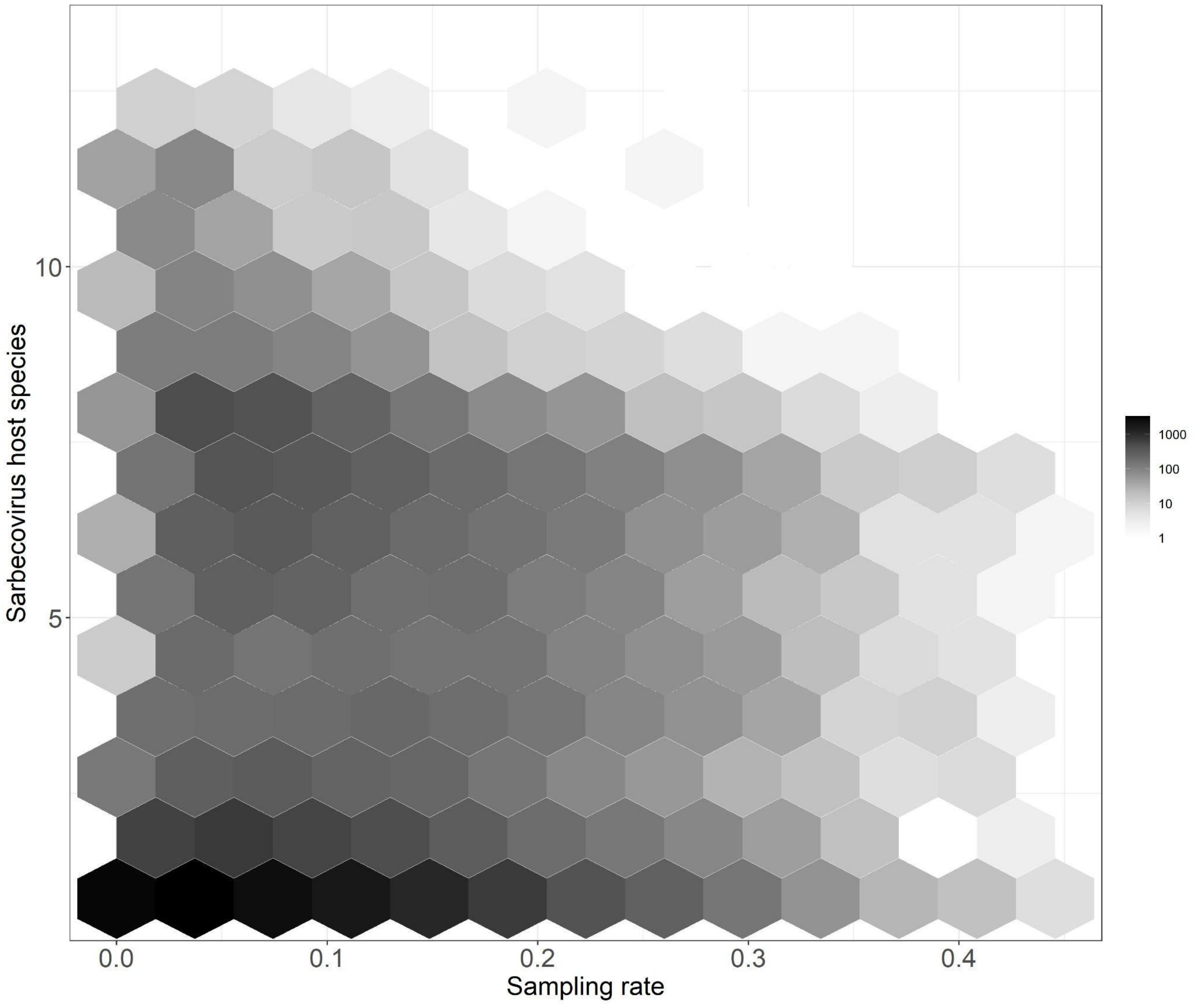


**Figure S11.** Pixel count of richness values and sampling rates across our study region.

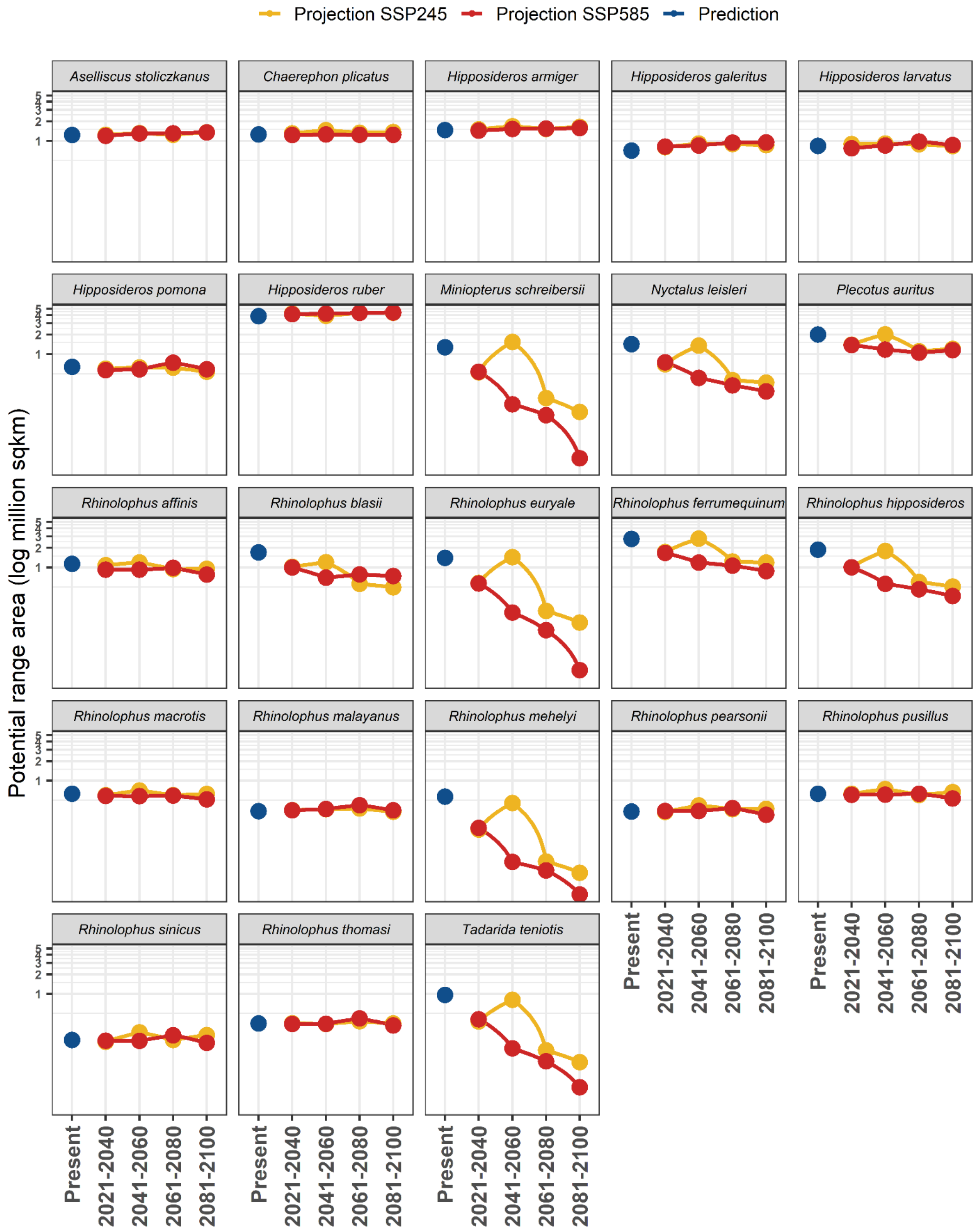


**Figure S12.** Potential range in the current model ensemble's predictions (blue) and future projections for each species according to SSP585 (red) and SSP245 (yellow) projections considering BCC-CSM2-MR.

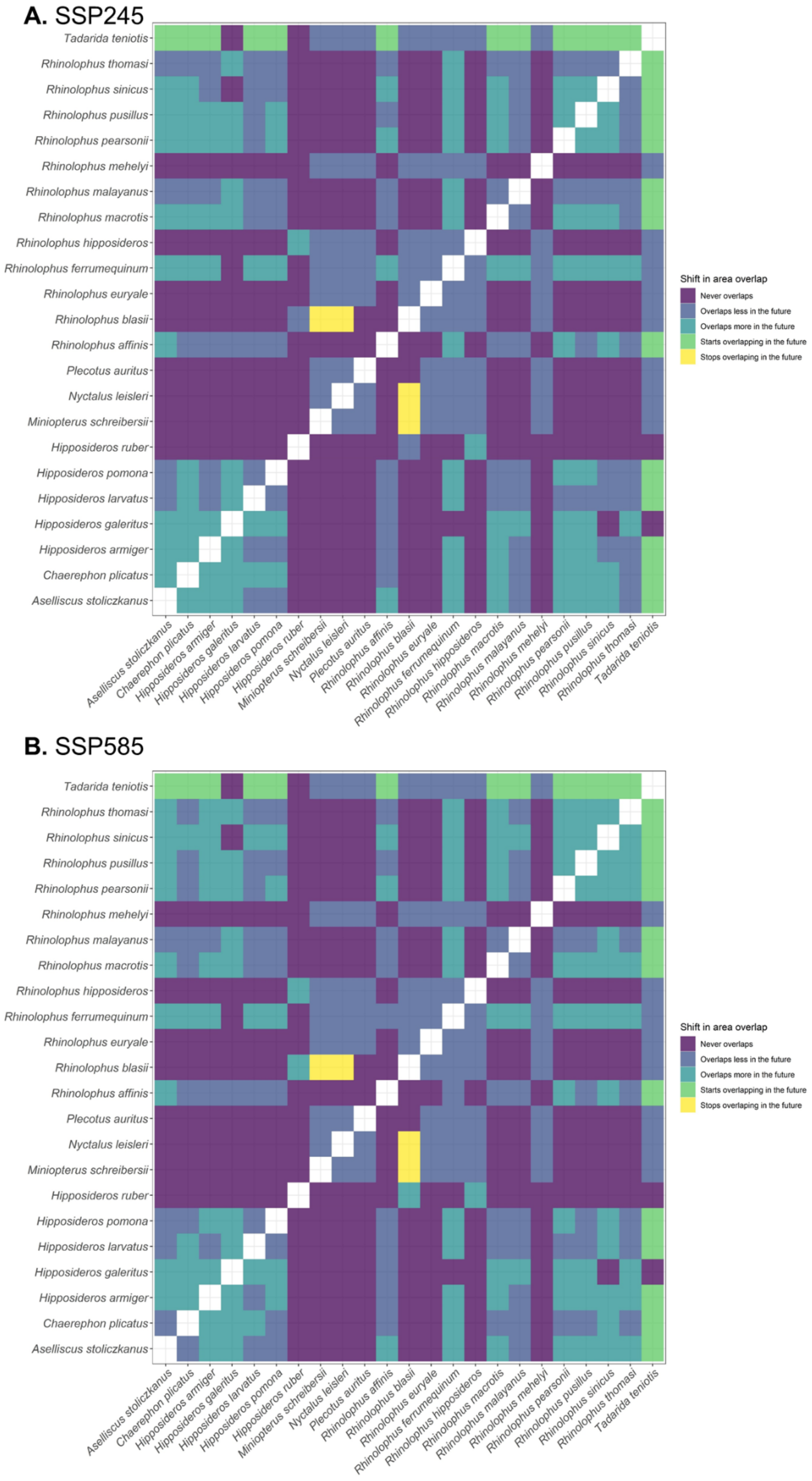


**Figure S13.** Overlap shifts in the future, considering the scenario SSP245 in 2100 (A) and SSP585 in 2100 (B) in comparison to the present.

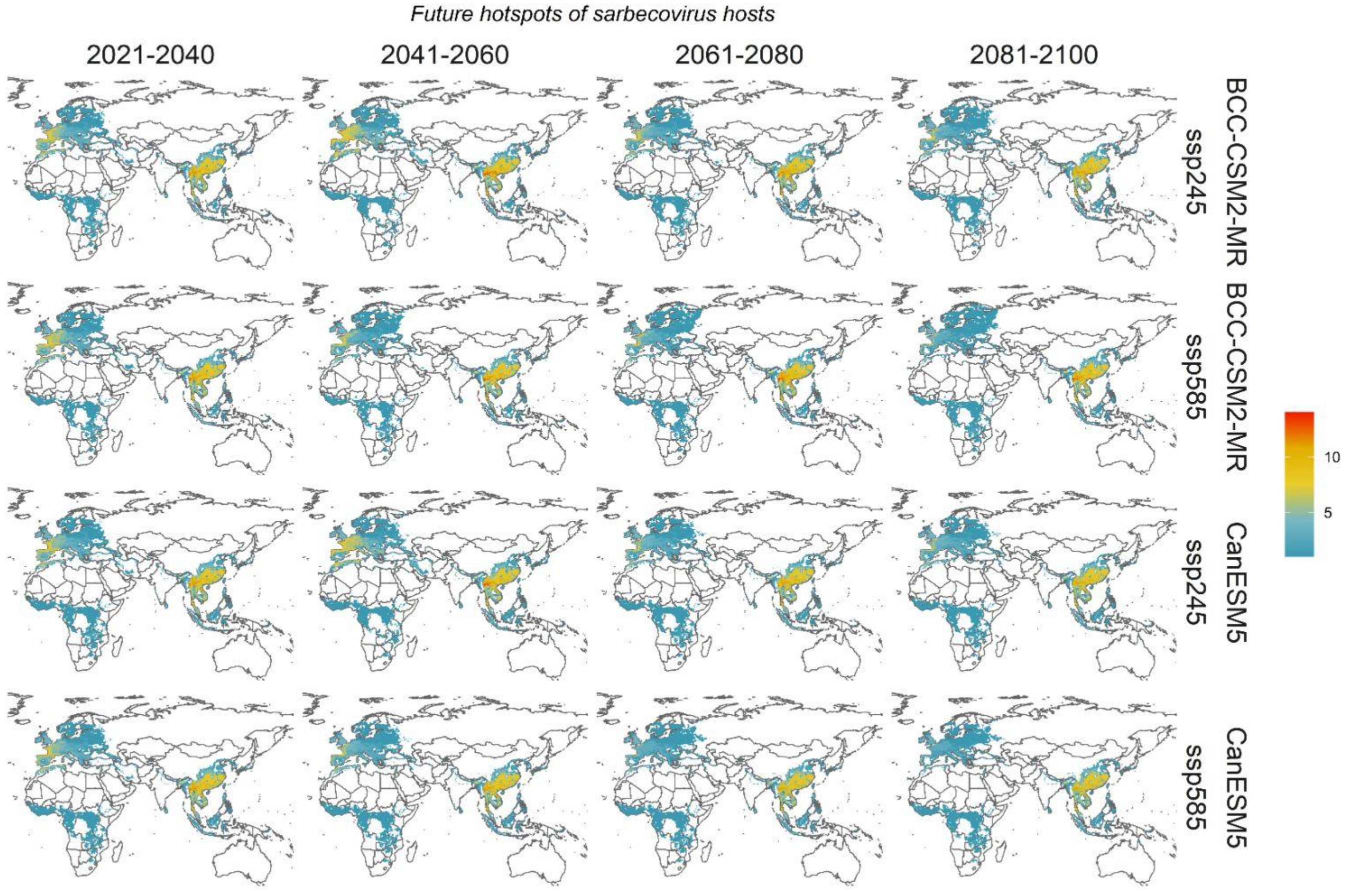


**Figure S14.** Future potential distribution of sarbecovirus bat hosts. Distribution across all periods, emission scenarios (SSP), and GCMs in the future.

**Table S1.** Target species for ecological niche models and for improving models estimating the risk of pathogen emergence. Sarbecoviruses have been detected in individuals of these species. * species not modelled. DOI: Digital object identifier for the references.

| Bat host of sarbecovirus | Family | DOI(s) |
|---|---|---|
| *Aselliscus stoliczkanus* | Hipposideridae | 10.1371/journal.ppat.1006698; 10.1038/s41467-020-17687-3 |
| *Hipposideros armiger* | Hipposideridae | 10.1093/ve/vex012 |
| *Hipposideros galeritus* | Hipposideridae | www.researchsquare.com/article/rs-885194/v1 |
| *Hipposideros larvatus* | Hipposideridae | doi.org/10.1093/ve/vex012 |
| *Hipposideros pomona (gentilis)* | Hipposideridae | www.nature.com/articles/s41467-020-17687-3 |
| *Hipposideros pratti* | Hipposideridae | www.nature.com/articles/s41467-020-17687-3 |
| *Hipposideros ruber* | Hipposideridae | doi.org/10.1093/ve/vex012 |
| *Miniopterus schreibersii* | Miniopteridae | 10.1016/j.virol.2017.03.019 |
| *Chaerephon plicatus* | Molossidae | 10.3201/eid1906.121648; 10.1038/ismej.2015.138 |
| *Tadarida teniotis* | Molossidae | 10.1007/s11262-018-1614-8. |
| *Rhinolophus acuminatus* | Rhinolophidae | 10.1038/s41467-021-21240-1 |
| *Rhinolophus affinis* | Rhinolophidae | 10.1128/JVI.00631-14; https://doi.org/10.1038/s41586-020-2012-7 |
| *Rhinolophus blasii* | Rhinolophidae | 10.1128/JVI.00650-10 |
| *Rhinolophus blythi* | Rhinolophidae | www.tandfonline.com/doi/full/10.1080/22221751.2021.1964925 |
| *Rhinolophus cornutus* | Rhinolophidae | 10.1093/database/bau021 |
| *Rhinolophus creaghi** | Rhinolophidae | doi.org/10.1093/ve/vex012 |
| *Rhinolophus euryale* | Rhinolophidae | 10.1128/JVI.00650-10 |

| | | |
|---|---|---|
| *Rhinolophus ferrumequinum* | Rhinolophidae | 10.1093/ve/vex012; 10.1371/journal.ppat.1006698 |
| *Rhinolophus hipposideros* | Rhinolophidae | 10.1007/s00705-010-0612-5 |
| *Rhinolophus luctus* | Rhinolophidae | www.researchsquare.com/article/rs-885194/v1 |
| *Rhinolophus macrotis* | Rhinolophidae | 10.1126/science.1118391 |
| *Rhinolophus malayanus* | Rhinolophidae | 10.1371/journal.ppat.1006698; doi.org/10.1016/j.cell.2021.06.008 |
| *Rhinolophus marshalli* | Rhinolophidae | 10.21203/rs.3.rs-871965/v1 |
| *Rhinolophus mehelyi* | Rhinolophidae | 10.1128/JVI.00650-10 |
| *Rhinolophus monoceros* | Rhinolophidae | 10.1111/zph.12271 |
| *Rhinolophus pearsonii* | Rhinolophidae | 10.1016/j.virol.2017.03.019 |
| *Rhinolophus pusillus* | Rhinolophidae | 10.1016/j.virol.2017.03.019; 10.1038/s41467-020-17687-3 |
| *Rhinolophus rex* | Rhinolophidae | 10.21203/rs.3.rs-885194/v1 |
| *Rhinolophus shameli* | Rhinolophidae | doi.org/10.1016/j.meegid.2016.11.029. |
| *Rhinolophus siamensis* | Rhinolophidae | www.researchsquare.com/article/rs-885194/v1 |
| *Rhinolophus sinicus* | Rhinolophidae | doi.org/10.1038/s41467-020-17687-3 |
| *Rhinolophus stheno* | Rhinolophidae | 10.1016/j.cell.2021.06.008 |
| *Rhinolophus thomasi* | Rhinolophidae | 10.1093/ve/vex012 |
| *Nyctalus leisleri* | Vespertilionidae | 10.1128/JVI.00650-10 |
| *Plecotus auritus* | Vespertilionidae | 10.1007/s11262-018-1614-8 |

**Table S2.** Input terms for sampling bias analysis ran for sarbecovirus hosts (target species).

| Function term | Description | Value |
|---|---|---|
| res | Resolution | 0.25 |
| buffer | Buffer around the area for accounting for bias coming from neighbourhood structures | 2 |
| mcmc_burnin | number of iterations for the Markov chain Monte Carlo (MCMC) | 5000 |
| mcmc_iteractions | burn-in for the MCMC | 50000 |

**Table S3.** Covariates used in the dynamic pipeline for modelling present and future distribution of bat hosts of Sarbecovirus.

| Layer source | Description | Spatial resolution processed (dd) | Spatial resolution reported | Time period | Website | References |
|---|---|---|---|---|---|---|
| Distance to WHYMAP_WOKAM cave or karst composite. | We retrieved the karst layer from the “Karstifiable rocks” layer of the World Karst Aquifer Map (WOKAM) by WHYMAP1. This layer includes carbonatic rocks and evaporites. To include other geological formations possibly characterized by caves, we extracted observed cave locations from OpenStreetMap using the Overpass API. We used the Overpass Turbo website (https://overpass-turbo.eu) to query for worldwide cave locations. While this resulting cave locations layer is clearly affected by some bias deriving from OSM’s citizen-based mapping approach, it complements the karst layer with non-karst cave locations, such as coastal, volcanic, and artificial caves. To obtain an intersection of the two sources while keeping a transparent procedure, we computed the | 0.25 | ~1km | Static | NA | This work; Chen, Z., Goldscheider, N., Auler, A., Bakalowicz, M., Broda, S., Drew, D., Hartmann, J., Jiang, G., Moosdorf, N., Richts, A., Stevanovic, Z., Veni, G., Dumont, A., Aureli, A., Clos, P., Krombholz, M. (2017). World Karst Aquifer Map (WHYMAP WOKAM). BGR, IAH, KIT, UNESCO, doi: 10.25928/b2.21_sfkq-r406. |

| | | | | | | |
|---|---|---|---|---|---|---|
| | Euclidean distance in km from both karst formations and caves, and selected, for each pixel, the minimum of these two values. The layers were warped onto a WGS84 - global Mercator projection to calculate the distances, and then reset to the original geographic WGS84 reference system. | | | | | |
| Primary forest cover | Habitat - % forest | 0.25 | 0.25 | nearly current and future | LUH2 from LULC<br><br>CMIP6_Land_Use_Harmonization_primf_2015 | |
| bio_1 | Annual Mean Temperature | 0.25 | 10 min | nearly current 1970-2000 and future | http://worldclim.org/version2 | WorldClim v02 Fick, S.E. and R.J. Hijmans, 2017. WorldClim 2: new 1km spatial resolution climate surfaces for global land areas. International Journal of Climatology 37 (12): 4302-4315. |

| | | | | | | |
|---|---|---|---|---|---|---|
| bio_2 | Mean Diurnal Range (Mean of monthly (max temp - min temp)) | 0.25 | 10 min | nearly current 1970-2000 and future | http://worldclim.org/version3 | WorldClim v02 Fick, S.E. and R.J. Hijmans, 2017. WorldClim 2: new 1km spatial resolution climate surfaces for global land areas. International Journal of Climatology 37 (12): 4302-4315. |
| bio_3 | Isothermality (BIO2/BIO7) (×100) | 0.25 | 10 min | nearly current 1970-2000 and future | http://worldclim.org/version4 | WorldClim v02 Fick, S.E. and R.J. Hijmans, 2017. WorldClim 2: new 1km spatial resolution climate surfaces for global land areas. International Journal of Climatology 37 (12): 4302-4315. |
| bio_4 | Temperature Seasonality (standard deviation ×100) | 0.25 | 10 min | nearly current 1970-2000 and future | http://worldclim.org/version5 | WorldClim v02 Fick, S.E. and R.J. Hijmans, 2017. WorldClim 2: new 1km spatial resolution climate surfaces for global land areas. |

| | | | | | | |
|---|---|---|---|---|---|---|
| | | | | | | International Journal of Climatology 37 (12): 4302-4315. |
| bio_5 | Max Temperature of Warmest Month | 0.25 | 10 min | nearly current 1970-2000 and future | http://worldclim.org/version6 | WorldClim v02 Fick, S.E. and R.J. Hijmans, 2017. WorldClim 2: new 1km spatial resolution climate surfaces for global land areas. International Journal of Climatology 37 (12): 4302-4315. |
| bio_6 | Min Temperature of Coldest Month | 0.25 | 10 min | nearly current 1970-2000 and future | http://worldclim.org/version7 | WorldClim v02 Fick, S.E. and R.J. Hijmans, 2017. WorldClim 2: new 1km spatial resolution climate surfaces for global land areas. International Journal of Climatology 37 (12): 4302-4315. |

| | | | | | | |
|---|---|---|---|---|---|---|
| bio_7 | Temperature Annual Range (BIO5-BIO6) | 0.25 | 10 min | nearly current 1970-2000 and future | http://worldclim.org/version8 | WorldClim v02 Fick, S.E. and R.J. Hijmans, 2017. WorldClim 2: new 1km spatial resolution climate surfaces for global land areas. International Journal of Climatology 37 (12): 4302-4315. |
| bio_8 | Mean Temperature of Wettest Quarter | 0.25 | 10 min | nearly current 1970-2000 and future | http://worldclim.org/version9 | WorldClim v02 Fick, S.E. and R.J. Hijmans, 2017. WorldClim 2: new 1km spatial resolution climate surfaces for global land areas. International Journal of Climatology 37 (12): 4302-4315. |
| bio_9 | Mean Temperature of Driest Quarter | 0.25 | 10 min | nearly current 1970-2000 and future | http://worldclim.org/version10 | WorldClim v02 Fick, S.E. and R.J. Hijmans, 2017. WorldClim 2: new 1km spatial resolution climate surfaces for global land areas. |

| | | | | | | |
|---|---|---|---|---|---|---|
| | | | | | | International Journal of Climatology 37 (12): 4302-4315. |
| bio_10 | Mean Temperature of Warmest Quarter | 0.25 | 10 min | nearly current 1970-2000 and future | http://worldclim.org/version11 | WorldClim v02 Fick, S.E. and R.J. Hijmans, 2017. WorldClim 2: new 1km spatial resolution climate surfaces for global land areas. International Journal of Climatology 37 (12): 4302-4315. |
| bio_11 | Mean Temperature of Coldest Quarter | 0.25 | 10 min | nearly current 1970-2000 and future | http://worldclim.org/version12 | WorldClim v02 Fick, S.E. and R.J. Hijmans, 2017. WorldClim 2: new 1km spatial resolution climate surfaces for global land areas. International Journal of Climatology 37 (12): 4302-4315. |

| | | | | | | |
|---|---|---|---|---|---|---|
| bio_12 | Annual Precipitation | 0.25 | 10 min | nearly current 1970-2000 and future | http://worldclim.org/version13 | WorldClim v02 Fick, S.E. and R.J. Hijmans, 2017. WorldClim 2: new 1km spatial resolution climate surfaces for global land areas. International Journal of Climatology 37 (12): 4302-4315. |
| bio_13 | Precipitation of Wettest Month | 0.25 | 10 min | nearly current 1970-2000 and future | http://worldclim.org/version14 | WorldClim v02 Fick, S.E. and R.J. Hijmans, 2017. WorldClim 2: new 1km spatial resolution climate surfaces for global land areas. International Journal of Climatology 37 (12): 4302-4315. |
| bio_14 | Precipitation of Driest Month | 0.25 | 10 min | nearly current 1970-2000 and future | http://worldclim.org/version15 | WorldClim v02 Fick, S.E. and R.J. Hijmans, 2017. WorldClim 2: new 1km spatial resolutionclimate surfaces for global land areas. |

| | | | | | | |
|---|---|---|---|---|---|---|
| | | | | | | International Journal of Climatology 37 (12): 4302-4315. |
| bio_15 | Precipitation Seasonality (Coefficient of Variation) | 0.25 | 10 min | nearly current 1970-2000 and future | http://worldclim.org/version16 | WorldClim v02 Fick, S.E. and R.J. Hijmans, 2017. WorldClim 2: new 1km spatial resolution climate surfaces for global land areas. International Journal of Climatology 37 (12): 4302-4315. |
| bio_16 | Precipitation of Wettest Quarter | 0.25 | 10 min | nearly current 1970-2000 and future | http://worldclim.org/version17 | WorldClim v02 Fick, S.E. and R.J. Hijmans, 2017. WorldClim 2: new 1km spatial resolution climate surfaces for global land areas. International Journal of Climatology 37 (12): 4302-4315. |

| | | | | | | |
|---|---|---|---|---|---|---|
| bio_17 | Precipitation of Driest Quarter | 0.25 | 10 min | nearly current 1970-2000 and future | http://worldclim.org/version18 | WorldClim v02 Fick, S.E. and R.J. Hijmans, 2017. WorldClim 2: new 1km spatial resolution climate surfaces for global land areas. International Journal of Climatology 37 (12): 4302-4315. |
| bio_18 | Precipitation of Warmest Quarter | 0.25 | 10 min | nearly current 1970-2000 and future | http://worldclim.org/version19 | WorldClim v02 Fick, S.E. and R.J. Hijmans, 2017. WorldClim 2: new 1km spatial resolution climate surfaces for global land areas. International Journal of Climatology 37 (12): 4302-4315. |
| bio_19 | Precipitation of Coldest Quarter | 0.25 | 10 min | nearly current 1970-2000 and future | http://worldclim.org/version20 | WorldClim v02 Fick, S.E. and R.J. Hijmans, 2017. WorldClim 2: new 1km spatial resolution climate surfaces for global land areas. |

International Journal
of Climatology
37 (12):
4302-4315.

**Table S4.** Ensemble forecast performance for each species, considering TSS values, Boyce index and their respective standard deviation values (SD). *Positive values mean % TSS improvement after intersecting with IUCN ranges ('_i'). NA= not available (species with less than 40 unbiased occurrences after intersecting with IUCN polygon). n = number of occurrences.

| Species | n | TSS | TSS_SD | Boyce | Boyce_SD | n_i | TSS_i | TSS_SD_i | Boyce_i | Boyce_SD_i | pct_tss_improved_after_i |
|---|---|---|---|---|---|---|---|---|---|---|---|
| *Rhinolophus pusillus* | 180 | 0.86 | 0.03 | 0.87 | 0.07 | 116 | 0.65 | 0.06 | 0.74 | 0.11 | -24.23 |
| *Hipposideros ruber* | 92 | 0.76 | 0.07 | 0.66 | 0.11 | 85 | 0.62 | 0.05 | 0.29 | 0.22 | -18.01 |
| *Rhinolophus mehelyi* | 176 | 0.89 | 0.04 | 0.65 | 0.12 | 125 | 0.77 | 0.06 | 0.62 | 0.22 | -13.38 |
| *Rhinolophus sinicus* | 121 | 0.83 | 0.05 | 0.85 | 0.07 | 105 | 0.73 | 0.04 | 0.82 | 0.08 | -11.85 |
| *Rhinolophus euryale* | 468 | 0.85 | 0.02 | 0.66 | 0.14 | 448 | 0.79 | 0.03 | 0.81 | 0.09 | -6.48 |
| *Hipposideros armiger* | 215 | 0.85 | 0.02 | 0.91 | 0.05 | 206 | 0.83 | 0.02 | 0.77 | 0.12 | -1.82 |
| *Plecotus auritus* | 1043 | 0.93 | 0.01 | 0.93 | 0.03 | 961 | 0.92 | 0.01 | 0.92 | 0.03 | -1.36 |
| *Nyctalus leisleri* | 626 | 0.91 | 0.01 | 0.87 | 0.06 | 572 | 0.90 | 0.01 | 0.92 | 0.04 | -1.08 |

| | | | | | | | | | | | |
|---|---|---|---|---|---|---|---|---|---|---|---|
| *Rhinolophus affinis* | 183 | 0.81 | 0.04 | 0.95 | 0.02 | 155 | 0.81 | 0.05 | 0.84 | 0.14 | -0.39 |
| *Rhinolophus hipposideros* | 727 | 0.90 | 0.02 | 0.91 | 0.07 | 707 | 0.90 | 0.01 | 0.79 | 0.12 | 0.25 |
| *Rhinolophus ferrumequinum* | 961 | 0.91 | 0.01 | 0.94 | 0.06 | 886 | 0.92 | 0.01 | 0.86 | 0.08 | 1.01 |
| *Aselliscus stoliczkanus* | 55 | 0.81 | 0.10 | 0.72 | 0.20 | 50 | 0.82 | 0.10 | 0.56 | 0.27 | 1.18 |
| *Miniopterus schreibersii* | 289 | 0.86 | 0.04 | 0.89 | 0.06 | 247 | 0.87 | 0.03 | 0.86 | 0.04 | 1.33 |
| *Tadarida teniotis* | 569 | 0.88 | 0.02 | 0.90 | 0.08 | 534 | 0.92 | 0.02 | 0.87 | 0.11 | 4.95 |
| *Hipposideros pomona* | 75 | 0.68 | 0.11 | 0.72 | 0.18 | 63 | 0.72 | 0.09 | 0.54 | 0.28 | 5.05 |
| *Rhinolophus pearsonii* | 66 | 0.79 | 0.03 | 0.71 | 0.11 | 52 | 0.84 | 0.08 | 0.58 | 0.16 | 6.47 |
| *Hipposideros larvatus* | 99 | 0.70 | 0.05 | 0.49 | 0.16 | 91 | 0.83 | 0.10 | 0.79 | 0.14 | 19.94 |

Six species that could only be modelled without intersecting their points with IUCN data

| | | | | | | | | | | | |
|---|---|---|---|---|---|---|---|---|---|---|---|
| *Rhinolophus blasii* | 47 | 0.82 | 0.11 | 0.57 | 0.31 | #N/A | #N/A | #N/A | #N/A | #N/A | #N/A |
| *Rhinolophus malayanus* | 41 | 0.80 | 0.09 | 0.50 | 0.19 | #N/A | #N/A | #N/A | #N/A | #N/A | #N/A |
| *Rhinolophus thomasi* | 52 | 0.80 | 0.12 | 0.53 | 0.33 | #N/A | #N/A | #N/A | #N/A | #N/A | #N/A |
| *Chaerephon plicatus* | 43 | 0.79 | 0.07 | 0.47 | 0.16 | #N/A | #N/A | #N/A | #N/A | #N/A | #N/A |
| *Rhinolophus macrotis* | 79 | 0.76 | 0.04 | 0.67 | 0.20 | #N/A | #N/A | #N/A | #N/A | #N/A | #N/A |
| *Hipposideros galeritus* | 56 | 0.54 | 0.13 | 0.44 | 0.26 | #N/A | #N/A | #N/A | #N/A | #N/A | #N/A |

**Table S5.** Current potential ranges and shifts per species considering SSP245 and SSP585 BCC-CSM2-MR by 20100. All area values are given in square kilometers.

| Species | Family | Potential range | Future range SSP 585 | shift SSP 585 | Future range SSP 245 BCC | shift SSP 245 |
|---|---|---|---|---|---|---|
| *Aselliscus stoliczkanus* | Hipposiderida e | 1237714.40 | 1354674.22 | expansion | 1360481.41 | expansion |
| *Hipposideros armiger* | Hipposiderida e | 1472900.56 | 1575768.92 | expansion | 1639022.97 | expansion |
| *Hipposideros galeritus* | Hipposiderida e | 708237.78 | 947935.47 | expansion | 860848.81 | expansion |
| *Hipposideros larvatus* | Hipposiderida e | 840365.42 | 866014.11 | expansion | 832310.19 | contraction |
| *Hipposideros pomona* | Hipposiderida e | 636663.45 | 588897.54 | contraction | 535044.21 | contraction |
| *Hipposideros ruber* | Hipposiderida e | 3845211.87 | 4380332.98 | expansion | 4330393.50 | expansion |
| *Miniopterus schreibersii* | Miniopteridae | 1281327.42 | 25114.56 | contraction | 129159.98 | contraction |
| *Chaerephon plicatus* | Molossidae | 1262708.06 | 1244615.17 | contraction | 1369702.85 | expansion |
| *Tadarida teniotis* | Molossidae | 964636.56 | 36796.23 | contraction | 89358.43 | contraction |
| *Rhinolophus affinis* | Rhinolophidae | 1138906.15 | 774616.08 | contraction | 955821.49 | contraction |
| *Rhinolophus blasii* | Rhinolophidae | 1702219.57 | 740211.72 | contraction | 495957.91 | contraction |
| *Rhinolophus euryale* | Rhinolophidae | 1400870.88 | 26370.68 | contraction | 141671.43 | contraction |
| *Rhinolophus ferrumequinum* | Rhinolophidae | 2746040.19 | 879230.53 | contraction | 1185782.66 | contraction |
| *Rhinolophus hipposideros* | Rhinolophidae | 1876065.73 | 362834.71 | contraction | 507249.92 | contraction |
| *Rhinolophus macrotis* | Rhinolophidae | 628397.27 | 514795.32 | contraction | 624512.98 | contraction |
| *Rhinolophus malayanus* | Rhinolophidae | 338876.83 | 351169.85 | expansion | 333958.98 | contraction |
| *Rhinolophus mehelyi* | Rhinolophidae | 568970.21 | 17856.66 | contraction | 38414.11 | contraction |
| *Rhinolophus pearsonii* | Rhinolophidae | 334870.83 | 299161.94 | contraction | 368905.17 | expansion |
| *Rhinolophus pusillus* | Rhinolophidae | 630659.07 | 534844.10 | contraction | 665682.75 | expansion |
| *Rhinolophus sinicus* | Rhinolophidae | 195968.99 | 176958.90 | contraction | 233653.64 | expansion |
| *Rhinolophus thomasi* | Rhinolophidae | 351126.40 | 328172.47 | contraction | 351413.54 | expansion |
| *Nyctalus leisleri* | Vespertilionida e | 1431618.78 | 267786.90 | contraction | 365373.35 | contraction |
| *Plecotus auritus* | Vespertilionida e | 2006582.48 | 1153915.41 | contraction | 1206608.02 | contraction |

**Table S6.** Potential range for Sarbecovirus host species per GCM per algorithm, per scenario and per period.

| Potential range sqkm | binomial | facet | period | scenario | gcm | accessible area sqkm | Potential range million sqkm |
|---|---|---|---|---|---|---|---|
| 1237714.399 | *Aselliscus stoliczkanus* | Present | Present | #N/A | Present | 2476390 | 1.237714 |
| 1262708.061 | *Chaerephon plicatus* | Present | Present | #N/A | Present | 2825780 | 1.262708 |
| 1472900.561 | *Hipposideros armiger* | Present | Present | #N/A | Present | 4137359 | 1.472901 |
| 708237.7781 | *Hipposideros galeritus* | Present | Present | #N/A | Present | 2227205 | 0.708238 |
| 840365.4218 | *Hipposideros larvatus* | Present | Present | #N/A | Present | 3315892 | 0.840365 |
| 636663.4465 | *Hipposideros pomona* | Present | Present | #N/A | Present | 3100912 | 0.636663 |
| 3845211.865 | *Hipposideros ruber* | Present | Present | #N/A | Present | 8487199 | 3.845212 |
| 1281327.424 | *Miniopterus schreibersii* | Present | Present | #N/A | Present | 2816421 | 1.281327 |
| 1431618.785 | *Nyctalus leisleri* | Present | Present | #N/A | Present | 7413692 | 1.431619 |
| 2006582.48 | *Plecotus auritus* | Present | Present | #N/A | Present | 9355015 | 2.006582 |
| 1138906.151 | *Rhinolophus affinis* | Present | Present | #N/A | Present | 5133788 | 1.138906 |
| 1702219.571 | *Rhinolophus blasii* | Present | Present | #N/A | Present | 4751972 | 1.70222 |
| 1400870.884 | *Rhinolophus euryale* | Present | Present | #N/A | Present | 4353786 | 1.400871 |

| | | | | | | | |
|---|---|---|---|---|---|---|---|
| 2746040.192 | *Rhinolophus ferrumequinum* | Present | Present | #N/A | Present | 12254734 | 2.74604 |
| 1876065.726 | *Rhinolophus hipposideros* | Present | Present | #N/A | Present | 6445160 | 1.876066 |
| 628397.2666 | *Rhinolophus macrotis* | Present | Present | #N/A | Present | 3677114 | 0.628397 |
| 338876.8274 | *Rhinolophus malayanus* | Present | Present | #N/A | Present | 1452518 | 0.338877 |
| 568970.2082 | *Rhinolophus mehelyi* | Present | Present | #N/A | Present | 3479949 | 0.56897 |
| 334870.8282 | *Rhinolophus pearsonii* | Present | Present | #N/A | Present | 2947268 | 0.334871 |
| 630659.0663 | *Rhinolophus pusillus* | Present | Present | #N/A | Present | 4716619 | 0.630659 |
| 195968.9857 | *Rhinolophus sinicus* | Present | Present | #N/A | Present | 2907524 | 0.195969 |
| 351126.4045 | *Rhinolophus thomasi* | Present | Present | #N/A | Present | 2802593 | 0.351126 |
| 964636.5554 | *Tadarida teniotis* | Present | Present | #N/A | Present | 7994697 | 0.964637 |
| 1242349.045 | *Aselliscus stoliczkanus* | BCC-CSM2-MR_ssp245_2021-2040 | 2021-2040 | ssp245 | BCC-CSM2-MR | 2476390 | 1.242349 |
| 1309464.86 | *Chaerephon plicatus* | BCC-CSM2-MR_ssp245_2021-2040 | 2021-2040 | ssp245 | BCC-CSM2-MR | 2825780 | 1.309465 |
| 1510871.879 | *Hipposideros armiger* | BCC-CSM2-MR_ssp245_2021-2040 | 2021-2040 | ssp245 | BCC-CSM2-MR | 4137359 | 1.510872 |
| 800573.2324 | *Hipposideros galeritus* | BCC-CSM2-MR_ssp245_2021-2040 | 2021-2040 | ssp245 | BCC-CSM2-MR | 2227205 | 0.800573 |
| 894113.7166 | *Hipposideros larvatus* | BCC-CSM2-MR_ssp245_2021-2040 | 2021-2040 | ssp245 | BCC-CSM2-MR | 3315892 | 0.894114 |

| | | | | | | | |
|---|---|---|---|---|---|---|---|
| 597743.8134 | *Hipposideros pomona* | BCC-CSM2-MR_ssp245_2021-2040 | 2021-2040 | ssp245 | BCC-CSM2-MR | 3100912 | 0.597744 |
| 4205880.221 | *Hipposideros ruber* | BCC-CSM2-MR_ssp245_2021-2040 | 2021-2040 | ssp245 | BCC-CSM2-MR | 8487199 | 4.20588 |
| 527311.5906 | *Miniopterus schreibersii* | BCC-CSM2-MR_ssp245_2021-2040 | 2021-2040 | ssp245 | BCC-CSM2-MR | 2816421 | 0.527312 |
| 696519.7658 | *Nyctalus leisleri* | BCC-CSM2-MR_ssp245_2021-2040 | 2021-2040 | ssp245 | BCC-CSM2-MR | 7413692 | 0.69652 |
| 1398240.069 | *Plecotus auritus* | BCC-CSM2-MR_ssp245_2021-2040 | 2021-2040 | ssp245 | BCC-CSM2-MR | 9355015 | 1.39824 |
| 1087913.39 | *Rhinolophus affinis* | BCC-CSM2-MR_ssp245_2021-2040 | 2021-2040 | ssp245 | BCC-CSM2-MR | 5133788 | 1.087913 |
| 1024227.748 | *Rhinolophus blasii* | BCC-CSM2-MR_ssp245_2021-2040 | 2021-2040 | ssp245 | BCC-CSM2-MR | 4751972 | 1.024228 |
| 576178.7155 | *Rhinolophus euryale* | BCC-CSM2-MR_ssp245_2021-2040 | 2021-2040 | ssp245 | BCC-CSM2-MR | 4353786 | 0.576179 |
| 1729248.263 | *Rhinolophus ferrumequinum* | BCC-CSM2-MR_ssp245_2021-2040 | 2021-2040 | ssp245 | BCC-CSM2-MR | 12254734 | 1.729248 |
| 1002500.142 | *Rhinolophus hipposideros* | BCC-CSM2-MR_ssp245_2021-2040 | 2021-2040 | ssp245 | BCC-CSM2-MR | 6445160 | 1.0025 |
| 599568.0945 | *Rhinolophus macrotis* | BCC-CSM2-MR_ssp245_2021-2040 | 2021-2040 | ssp245 | BCC-CSM2-MR | 3677114 | 0.599568 |
| 353745.002 | *Rhinolophus malayanus* | BCC-CSM2-MR_ssp245_2021-2040 | 2021-2040 | ssp245 | BCC-CSM2-MR | 1452518 | 0.353745 |

| | | | | | | | |
|---|---|---|---|---|---|---|---|
| 177806.5308 | *Rhinolophus mehelyi* | BCC-CSM2-MR_ssp245_2021-2040 | 2021-2040 | ssp245 | BCC-CSM2-MR | 3479949 | 0.177807 |
| 330709.1194 | *Rhinolophus pearsonii* | BCC-CSM2-MR_ssp245_2021-2040 | 2021-2040 | ssp245 | BCC-CSM2-MR | 2947268 | 0.330709 |
| 629600.6279 | *Rhinolophus pusillus* | BCC-CSM2-MR_ssp245_2021-2040 | 2021-2040 | ssp245 | BCC-CSM2-MR | 4716619 | 0.629601 |
| 183671.412 | *Rhinolophus sinicus* | BCC-CSM2-MR_ssp245_2021-2040 | 2021-2040 | ssp245 | BCC-CSM2-MR | 2907524 | 0.183671 |
| 354041.1929 | *Rhinolophus thomasi* | BCC-CSM2-MR_ssp245_2021-2040 | 2021-2040 | ssp245 | BCC-CSM2-MR | 2802593 | 0.354041 |
| 377734.9854 | *Tadarida teniotis* | BCC-CSM2-MR_ssp245_2021-2040 | 2021-2040 | ssp245 | BCC-CSM2-MR | 7994697 | 0.377735 |
| 1317184.876 | *Aselliscus stoliczkanus* | BCC-CSM2-MR_ssp245_2041-2060 | 2041-2060 | ssp245 | BCC-CSM2-MR | 2476390 | 1.317185 |
| 1469092.962 | *Chaerephon plicatus* | BCC-CSM2-MR_ssp245_2041-2060 | 2041-2060 | ssp245 | BCC-CSM2-MR | 2825780 | 1.469093 |
| 1680928.477 | *Hipposideros armiger* | BCC-CSM2-MR_ssp245_2041-2060 | 2041-2060 | ssp245 | BCC-CSM2-MR | 4137359 | 1.680928 |
| 912779.8597 | *Hipposideros galeritus* | BCC-CSM2-MR_ssp245_2041-2060 | 2041-2060 | ssp245 | BCC-CSM2-MR | 2227205 | 0.91278 |
| 912034.8283 | *Hipposideros larvatus* | BCC-CSM2-MR_ssp245_2041-2060 | 2041-2060 | ssp245 | BCC-CSM2-MR | 3315892 | 0.912035 |
| 625530.9503 | *Hipposideros pomona* | BCC-CSM2-MR_ssp245_2041-2060 | 2041-2060 | ssp245 | BCC-CSM2-MR | 3100912 | 0.625531 |

| | | | | | | | |
|---|---|---|---|---|---|---|---|
| 3900497.725 | *Hipposideros ruber* | BCC-CSM2-MR_ssp245_2041-2060 | 2041-2060 | ssp245 | BCC-CSM2-MR | 8487199 | 3.900498 |
| 1542172.095 | *Miniopterus schreibersii* | BCC-CSM2-MR_ssp245_2041-2060 | 2041-2060 | ssp245 | BCC-CSM2-MR | 2816421 | 1.542172 |
| 1351831.355 | *Nyctalus leisleri* | BCC-CSM2-MR_ssp245_2041-2060 | 2041-2060 | ssp245 | BCC-CSM2-MR | 7413692 | 1.351831 |
| 2024303.67 | *Plecotus auritus* | BCC-CSM2-MR_ssp245_2041-2060 | 2041-2060 | ssp245 | BCC-CSM2-MR | 9355015 | 2.024304 |
| 1194480.797 | *Rhinolophus affinis* | BCC-CSM2-MR_ssp245_2041-2060 | 2041-2060 | ssp245 | BCC-CSM2-MR | 5133788 | 1.194481 |
| 1199657.187 | *Rhinolophus blasii* | BCC-CSM2-MR_ssp245_2041-2060 | 2041-2060 | ssp245 | BCC-CSM2-MR | 4751972 | 1.199657 |
| 1442923.471 | *Rhinolophus euryale* | BCC-CSM2-MR_ssp245_2041-2060 | 2041-2060 | ssp245 | BCC-CSM2-MR | 4353786 | 1.442923 |
| 2781081.748 | *Rhinolophus ferrumequinum* | BCC-CSM2-MR_ssp245_2041-2060 | 2041-2060 | ssp245 | BCC-CSM2-MR | 12254734 | 2.781082 |
| 1793885.481 | *Rhinolophus hipposideros* | BCC-CSM2-MR_ssp245_2041-2060 | 2041-2060 | ssp245 | BCC-CSM2-MR | 6445160 | 1.793885 |
| 707051.0018 | *Rhinolophus macrotis* | BCC-CSM2-MR_ssp245_2041-2060 | 2041-2060 | ssp245 | BCC-CSM2-MR | 3677114 | 0.707051 |
| 363630.6792 | *Rhinolophus malayanus* | BCC-CSM2-MR_ssp245_2041-2060 | 2041-2060 | ssp245 | BCC-CSM2-MR | 1452518 | 0.363631 |
| 452516.9024 | *Rhinolophus mehelyi* | BCC-CSM2-MR_ssp245_2041-2060 | 2041-2060 | ssp245 | BCC-CSM2-MR | 3479949 | 0.452517 |

| | | | | | | | |
|---|---|---|---|---|---|---|---|
| 417319.3994 | *Rhinolophus pearsonii* | BCC-CSM2-MR_ssp245_2041-2060 | 2041-2060 | ssp245 | BCC-CSM2-MR | 2947268 | 0.417319 |
| 744248.1518 | *Rhinolophus pusillus* | BCC-CSM2-MR_ssp245_2041-2060 | 2041-2060 | ssp245 | BCC-CSM2-MR | 4716619 | 0.744248 |
| 257231.7185 | *Rhinolophus sinicus* | BCC-CSM2-MR_ssp245_2041-2060 | 2041-2060 | ssp245 | BCC-CSM2-MR | 2907524 | 0.257232 |
| 345506.1447 | *Rhinolophus thomasi* | BCC-CSM2-MR_ssp245_2041-2060 | 2041-2060 | ssp245 | BCC-CSM2-MR | 2802593 | 0.345506 |
| 811467.5218 | *Tadarida teniotis* | BCC-CSM2-MR_ssp245_2041-2060 | 2041-2060 | ssp245 | BCC-CSM2-MR | 7994697 | 0.811468 |
| 1245863.534 | *Aselliscus stoliczkanus* | BCC-CSM2-MR_ssp245_2061-2080 | 2061-2080 | ssp245 | BCC-CSM2-MR | 2476390 | 1.245864 |
| 1333308.781 | *Chaerephon plicatus* | BCC-CSM2-MR_ssp245_2061-2080 | 2061-2080 | ssp245 | BCC-CSM2-MR | 2825780 | 1.333309 |
| 1529052.225 | *Hipposideros armiger* | BCC-CSM2-MR_ssp245_2061-2080 | 2061-2080 | ssp245 | BCC-CSM2-MR | 4137359 | 1.529052 |
| 894056.6989 | *Hipposideros galeritus* | BCC-CSM2-MR_ssp245_2061-2080 | 2061-2080 | ssp245 | BCC-CSM2-MR | 2227205 | 0.894057 |
| 890634.9158 | *Hipposideros larvatus* | BCC-CSM2-MR_ssp245_2061-2080 | 2061-2080 | ssp245 | BCC-CSM2-MR | 3315892 | 0.890635 |
| 621025.6654 | *Hipposideros pomona* | BCC-CSM2-MR_ssp245_2061-2080 | 2061-2080 | ssp245 | BCC-CSM2-MR | 3100912 | 0.621026 |
| 4318386.723 | *Hipposideros ruber* | BCC-CSM2-MR_ssp245_2061-2080 | 2061-2080 | ssp245 | BCC-CSM2-MR | 8487199 | 4.318387 |

| | | | | | | | |
|---|---|---|---|---|---|---|---|
| 210809.2368 | *Miniopterus schreibersii* | BCC-CSM2-MR_ssp245_2061-2080 | 2061-2080 | ssp245 | BCC-CSM2-MR | 2816421 | 0.210809 |
| 400368.1234 | *Nyctalus leisleri* | BCC-CSM2-MR_ssp245_2061-2080 | 2061-2080 | ssp245 | BCC-CSM2-MR | 7413692 | 0.400368 |
| 1110718.113 | *Plecotus auritus* | BCC-CSM2-MR_ssp245_2061-2080 | 2061-2080 | ssp245 | BCC-CSM2-MR | 9355015 | 1.110718 |
| 939959.8261 | *Rhinolophus affinis* | BCC-CSM2-MR_ssp245_2061-2080 | 2061-2080 | ssp245 | BCC-CSM2-MR | 5133788 | 0.93996 |
| 558271.5203 | *Rhinolophus blasii* | BCC-CSM2-MR_ssp245_2061-2080 | 2061-2080 | ssp245 | BCC-CSM2-MR | 4751972 | 0.558272 |
| 214324.416 | *Rhinolophus euryale* | BCC-CSM2-MR_ssp245_2061-2080 | 2061-2080 | ssp245 | BCC-CSM2-MR | 4353786 | 0.214324 |
| 1234700.718 | *Rhinolophus ferrumequinum* | BCC-CSM2-MR_ssp245_2061-2080 | 2061-2080 | ssp245 | BCC-CSM2-MR | 12254734 | 1.234701 |
| 598149.8379 | *Rhinolophus hipposideros* | BCC-CSM2-MR_ssp245_2061-2080 | 2061-2080 | ssp245 | BCC-CSM2-MR | 6445160 | 0.59815 |
| 598393.5508 | *Rhinolophus macrotis* | BCC-CSM2-MR_ssp245_2061-2080 | 2061-2080 | ssp245 | BCC-CSM2-MR | 3677114 | 0.598394 |
| 373165.7852 | *Rhinolophus malayanus* | BCC-CSM2-MR_ssp245_2061-2080 | 2061-2080 | ssp245 | BCC-CSM2-MR | 1452518 | 0.373166 |
| 57266.61354 | *Rhinolophus mehelyi* | BCC-CSM2-MR_ssp245_2061-2080 | 2061-2080 | ssp245 | BCC-CSM2-MR | 3479949 | 0.057267 |
| 365558.9677 | *Rhinolophus pearsonii* | BCC-CSM2-MR_ssp245_2061-2080 | 2061-2080 | ssp245 | BCC-CSM2-MR | 2947268 | 0.365559 |

| | | | | | | | |
|---|---|---|---|---|---|---|---|
| 604391.3202 | *Rhinolophus pusillus* | BCC-CSM2-MR_ssp245_2061-2080 | 2061-2080 | ssp245 | BCC-CSM2-MR | 4716619 | 0.604391 |
| 196593.4038 | *Rhinolophus sinicus* | BCC-CSM2-MR_ssp245_2061-2080 | 2061-2080 | ssp245 | BCC-CSM2-MR | 2907524 | 0.196593 |
| 379873.5732 | *Rhinolophus thomasi* | BCC-CSM2-MR_ssp245_2061-2080 | 2061-2080 | ssp245 | BCC-CSM2-MR | 2802593 | 0.379874 |
| 135735.1343 | *Tadarida teniotis* | BCC-CSM2-MR_ssp245_2061-2080 | 2061-2080 | ssp245 | BCC-CSM2-MR | 7994697 | 0.135735 |
| 1360481.411 | *Aselliscus stoliczkanus* | BCC-CSM2-MR_ssp245_2081-2100 | 2081-2100 | ssp245 | BCC-CSM2-MR | 2476390 | 1.360481 |
| 1369702.846 | *Chaerephon plicatus* | BCC-CSM2-MR_ssp245_2081-2100 | 2081-2100 | ssp245 | BCC-CSM2-MR | 2825780 | 1.369703 |
| 1639022.969 | *Hipposideros armiger* | BCC-CSM2-MR_ssp245_2081-2100 | 2081-2100 | ssp245 | BCC-CSM2-MR | 4137359 | 1.639023 |
| 860848.8141 | *Hipposideros galeritus* | BCC-CSM2-MR_ssp245_2081-2100 | 2081-2100 | ssp245 | BCC-CSM2-MR | 2227205 | 0.860849 |
| 832310.1943 | *Hipposideros larvatus* | BCC-CSM2-MR_ssp245_2081-2100 | 2081-2100 | ssp245 | BCC-CSM2-MR | 3315892 | 0.83231 |
| 535044.2082 | *Hipposideros pomona* | BCC-CSM2-MR_ssp245_2081-2100 | 2081-2100 | ssp245 | BCC-CSM2-MR | 3100912 | 0.535044 |
| 4330393.505 | *Hipposideros ruber* | BCC-CSM2-MR_ssp245_2081-2100 | 2081-2100 | ssp245 | BCC-CSM2-MR | 8487199 | 4.330394 |
| 129159.9788 | *Miniopterus schreibersii* | BCC-CSM2-MR_ssp245_2081-2100 | 2081-2100 | ssp245 | BCC-CSM2-MR | 2816421 | 0.12916 |

| | | | | | | | |
|---|---|---|---|---|---|---|---|
| 365373.3467 | *Nyctalus leisleri* | BCC-CSM2-MR_ssp245_2081-2100 | 2081-2100 | ssp245 | BCC-CSM2-MR | 7413692 | 0.365373 |
| 1206608.018 | *Plecotus auritus* | BCC-CSM2-MR_ssp245_2081-2100 | 2081-2100 | ssp245 | BCC-CSM2-MR | 9355015 | 1.206608 |
| 955821.4864 | *Rhinolophus affinis* | BCC-CSM2-MR_ssp245_2081-2100 | 2081-2100 | ssp245 | BCC-CSM2-MR | 5133788 | 0.955821 |
| 495957.9146 | *Rhinolophus blasii* | BCC-CSM2-MR_ssp245_2081-2100 | 2081-2100 | ssp245 | BCC-CSM2-MR | 4751972 | 0.495958 |
| 141671.4336 | *Rhinolophus euryale* | BCC-CSM2-MR_ssp245_2081-2100 | 2081-2100 | ssp245 | BCC-CSM2-MR | 4353786 | 0.141671 |
| 1185782.664 | *Rhinolophus ferrumequinum* | BCC-CSM2-MR_ssp245_2081-2100 | 2081-2100 | ssp245 | BCC-CSM2-MR | 12254734 | 1.185783 |
| 507249.9237 | *Rhinolophus hipposideros* | BCC-CSM2-MR_ssp245_2081-2100 | 2081-2100 | ssp245 | BCC-CSM2-MR | 6445160 | 0.50725 |
| 624512.9791 | *Rhinolophus macrotis* | BCC-CSM2-MR_ssp245_2081-2100 | 2081-2100 | ssp245 | BCC-CSM2-MR | 3677114 | 0.624513 |
| 333958.9788 | *Rhinolophus malayanus* | BCC-CSM2-MR_ssp245_2081-2100 | 2081-2100 | ssp245 | BCC-CSM2-MR | 1452518 | 0.333959 |
| 38414.1052 | *Rhinolophus mehelyi* | BCC-CSM2-MR_ssp245_2081-2100 | 2081-2100 | ssp245 | BCC-CSM2-MR | 3479949 | 0.038414 |
| 368905.1746 | *Rhinolophus pearsonii* | BCC-CSM2-MR_ssp245_2081-2100 | 2081-2100 | ssp245 | BCC-CSM2-MR | 2947268 | 0.368905 |
| 665682.748 | *Rhinolophus pusillus* | BCC-CSM2-MR_ssp245_2081-2100 | 2081-2100 | ssp245 | BCC-CSM2-MR | 4716619 | 0.665683 |

| | | | | | | | |
|---|---|---|---|---|---|---|---|
| 233653.6436 | *Rhinolophus sinicus* | BCC-CSM2-MR_ssp245_2081-2100 | 2081-2100 | ssp245 | BCC-CSM2-MR | 2907524 | 0.233654 |
| 351413.5405 | *Rhinolophus thomasi* | BCC-CSM2-MR_ssp245_2081-2100 | 2081-2100 | ssp245 | BCC-CSM2-MR | 2802593 | 0.351414 |
| 89358.4283 | *Tadarida teniotis* | BCC-CSM2-MR_ssp245_2081-2100 | 2081-2100 | ssp245 | BCC-CSM2-MR | 7994697 | 0.089358 |
| 1197727.429 | *Aselliscus stoliczkanus* | BCC-CSM2-MR_ssp585_2021-2040 | 2021-2040 | ssp585 | BCC-CSM2-MR | 2476390 | 1.197727 |
| 1234186.238 | *Chaerephon plicatus* | BCC-CSM2-MR_ssp585_2021-2040 | 2021-2040 | ssp585 | BCC-CSM2-MR | 2825780 | 1.234186 |
| 1445308.967 | *Hipposideros armiger* | BCC-CSM2-MR_ssp585_2021-2040 | 2021-2040 | ssp585 | BCC-CSM2-MR | 4137359 | 1.445309 |
| 819764.5974 | *Hipposideros galeritus* | BCC-CSM2-MR_ssp585_2021-2040 | 2021-2040 | ssp585 | BCC-CSM2-MR | 2227205 | 0.819765 |
| 772763.4799 | *Hipposideros larvatus* | BCC-CSM2-MR_ssp585_2021-2040 | 2021-2040 | ssp585 | BCC-CSM2-MR | 3315892 | 0.772763 |
| 569203.2687 | *Hipposideros pomona* | BCC-CSM2-MR_ssp585_2021-2040 | 2021-2040 | ssp585 | BCC-CSM2-MR | 3100912 | 0.569203 |
| 4135629.367 | *Hipposideros ruber* | BCC-CSM2-MR_ssp585_2021-2040 | 2021-2040 | ssp585 | BCC-CSM2-MR | 8487199 | 4.135629 |
| 536601.6236 | *Miniopterus schreibersii* | BCC-CSM2-MR_ssp585_2021-2040 | 2021-2040 | ssp585 | BCC-CSM2-MR | 2816421 | 0.536602 |
| 747603.5199 | *Nyctalus leisleri* | BCC-CSM2-MR_ssp585_2021-2040 | 2021-2040 | ssp585 | BCC-CSM2-MR | 7413692 | 0.747604 |

| | | | | | | | |
|---|---|---|---|---|---|---|---|
| 1381048.203 | *Plecotus auritus* | BCC-CSM2-MR_ssp585_2021-2040 | 2021-2040 | ssp585 | BCC-CSM2-MR | 9355015 | 1.381048 |
| 922385.0091 | *Rhinolophus affinis* | BCC-CSM2-MR_ssp585_2021-2040 | 2021-2040 | ssp585 | BCC-CSM2-MR | 5133788 | 0.922385 |
| 995624.6105 | *Rhinolophus blasii* | BCC-CSM2-MR_ssp585_2021-2040 | 2021-2040 | ssp585 | BCC-CSM2-MR | 4751972 | 0.995625 |
| 568168.5082 | *Rhinolophus euryale* | BCC-CSM2-MR_ssp585_2021-2040 | 2021-2040 | ssp585 | BCC-CSM2-MR | 4353786 | 0.568169 |
| 1667942.644 | *Rhinolophus ferrumequinum* | BCC-CSM2-MR_ssp585_2021-2040 | 2021-2040 | ssp585 | BCC-CSM2-MR | 12254734 | 1.667943 |
| 1001724.676 | *Rhinolophus hipposideros* | BCC-CSM2-MR_ssp585_2021-2040 | 2021-2040 | ssp585 | BCC-CSM2-MR | 6445160 | 1.001725 |
| 584738.8701 | *Rhinolophus macrotis* | BCC-CSM2-MR_ssp585_2021-2040 | 2021-2040 | ssp585 | BCC-CSM2-MR | 3677114 | 0.584739 |
| 350610.0967 | *Rhinolophus malayanus* | BCC-CSM2-MR_ssp585_2021-2040 | 2021-2040 | ssp585 | BCC-CSM2-MR | 1452518 | 0.35061 |
| 187588.6355 | *Rhinolophus mehelyi* | BCC-CSM2-MR_ssp585_2021-2040 | 2021-2040 | ssp585 | BCC-CSM2-MR | 3479949 | 0.187589 |
| 342239.1605 | *Rhinolophus pearsonii* | BCC-CSM2-MR_ssp585_2021-2040 | 2021-2040 | ssp585 | BCC-CSM2-MR | 2947268 | 0.342239 |
| 608199.168 | *Rhinolophus pusillus* | BCC-CSM2-MR_ssp585_2021-2040 | 2021-2040 | ssp585 | BCC-CSM2-MR | 4716619 | 0.608199 |
| 191633.8932 | *Rhinolophus sinicus* | BCC-CSM2-MR_ssp585_2021-2040 | 2021-2040 | ssp585 | BCC-CSM2-MR | 2907524 | 0.191634 |

| | | | | | | | |
|---|---|---|---|---|---|---|---|
| 344973.5921 | *Rhinolophus thomasi* | BCC-CSM2-MR_ssp585_2021-2040 | 2021-2040 | ssp585 | BCC-CSM2-MR | 2802593 | 0.344974 |
| 408595.9767 | *Tadarida teniotis* | BCC-CSM2-MR_ssp585_2021-2040 | 2021-2040 | ssp585 | BCC-CSM2-MR | 7994697 | 0.408596 |
| 1302219.505 | *Aselliscus stoliczkanus* | BCC-CSM2-MR_ssp585_2041-2060 | 2041-2060 | ssp585 | BCC-CSM2-MR | 2476390 | 1.30222 |
| 1255517.697 | *Chaerephon plicatus* | BCC-CSM2-MR_ssp585_2041-2060 | 2041-2060 | ssp585 | BCC-CSM2-MR | 2825780 | 1.255518 |
| 1528493.098 | *Hipposideros armiger* | BCC-CSM2-MR_ssp585_2041-2060 | 2041-2060 | ssp585 | BCC-CSM2-MR | 4137359 | 1.528493 |
| 849398.5758 | *Hipposideros galeritus* | BCC-CSM2-MR_ssp585_2041-2060 | 2041-2060 | ssp585 | BCC-CSM2-MR | 2227205 | 0.849399 |
| 851343.8892 | *Hipposideros larvatus* | BCC-CSM2-MR_ssp585_2041-2060 | 2041-2060 | ssp585 | BCC-CSM2-MR | 3315892 | 0.851344 |
| 585115.7725 | *Hipposideros pomona* | BCC-CSM2-MR_ssp585_2041-2060 | 2041-2060 | ssp585 | BCC-CSM2-MR | 3100912 | 0.585116 |
| 4197939.538 | *Hipposideros ruber* | BCC-CSM2-MR_ssp585_2041-2060 | 2041-2060 | ssp585 | BCC-CSM2-MR | 8487199 | 4.19794 |
| 169194.0909 | *Miniopterus schreibersii* | BCC-CSM2-MR_ssp585_2041-2060 | 2041-2060 | ssp585 | BCC-CSM2-MR | 2816421 | 0.169194 |
| 430435.5403 | *Nyctalus leisleri* | BCC-CSM2-MR_ssp585_2041-2060 | 2041-2060 | ssp585 | BCC-CSM2-MR | 7413692 | 0.430436 |
| 1179877.143 | *Plecotus auritus* | BCC-CSM2-MR_ssp585_2041-2060 | 2041-2060 | ssp585 | BCC-CSM2-MR | 9355015 | 1.179877 |

| | | | | | | | |
|---|---|---|---|---|---|---|---|
| 921496.6706 | *Rhinolophus affinis* | BCC-CSM2-MR_ssp585_2041-2060 | 2041-2060 | ssp585 | BCC-CSM2-MR | 5133788 | 0.921497 |
| 694689.7772 | *Rhinolophus blasii* | BCC-CSM2-MR_ssp585_2041-2060 | 2041-2060 | ssp585 | BCC-CSM2-MR | 4751972 | 0.69469 |
| 202372.2556 | *Rhinolophus euryale* | BCC-CSM2-MR_ssp585_2041-2060 | 2041-2060 | ssp585 | BCC-CSM2-MR | 4353786 | 0.202372 |
| 1189439.238 | *Rhinolophus ferrumequinum* | BCC-CSM2-MR_ssp585_2041-2060 | 2041-2060 | ssp585 | BCC-CSM2-MR | 12254734 | 1.189439 |
| 561271.6149 | *Rhinolophus hipposideros* | BCC-CSM2-MR_ssp585_2041-2060 | 2041-2060 | ssp585 | BCC-CSM2-MR | 6445160 | 0.561272 |
| 577948.406 | *Rhinolophus macrotis* | BCC-CSM2-MR_ssp585_2041-2060 | 2041-2060 | ssp585 | BCC-CSM2-MR | 3677114 | 0.577948 |
| 366516.9544 | *Rhinolophus malayanus* | BCC-CSM2-MR_ssp585_2041-2060 | 2041-2060 | ssp585 | BCC-CSM2-MR | 1452518 | 0.366517 |
| 56301.26034 | *Rhinolophus mehelyi* | BCC-CSM2-MR_ssp585_2041-2060 | 2041-2060 | ssp585 | BCC-CSM2-MR | 3479949 | 0.056301 |
| 342326.6633 | *Rhinolophus pearsonii* | BCC-CSM2-MR_ssp585_2041-2060 | 2041-2060 | ssp585 | BCC-CSM2-MR | 2947268 | 0.342327 |
| 611357.0672 | *Rhinolophus pusillus* | BCC-CSM2-MR_ssp585_2041-2060 | 2041-2060 | ssp585 | BCC-CSM2-MR | 4716619 | 0.611357 |
| 189740.224 | *Rhinolophus sinicus* | BCC-CSM2-MR_ssp585_2041-2060 | 2041-2060 | ssp585 | BCC-CSM2-MR | 2907524 | 0.18974 |
| 347668.6552 | *Rhinolophus thomasi* | BCC-CSM2-MR_ssp585_2041-2060 | 2041-2060 | ssp585 | BCC-CSM2-MR | 2802593 | 0.347669 |

| | | | | | | | |
|---|---|---|---|---|---|---|---|
| 145355.5461 | *Tadarida teniotis* | BCC-CSM2-MR_ssp585_2041-2060 | 2041-2060 | ssp585 | BCC-CSM2-MR | 7994697 | 0.145356 |
| 1311974.792 | *Aselliscus stoliczkanus* | BCC-CSM2-MR_ssp585_2061-2080 | 2061-2080 | ssp585 | BCC-CSM2-MR | 2476390 | 1.311975 |
| 1241583.649 | *Chaerephon plicatus* | BCC-CSM2-MR_ssp585_2061-2080 | 2061-2080 | ssp585 | BCC-CSM2-MR | 2825780 | 1.241584 |
| 1543240.556 | *Hipposideros armiger* | BCC-CSM2-MR_ssp585_2061-2080 | 2061-2080 | ssp585 | BCC-CSM2-MR | 4137359 | 1.543241 |
| 944038.8445 | *Hipposideros galeritus* | BCC-CSM2-MR_ssp585_2061-2080 | 2061-2080 | ssp585 | BCC-CSM2-MR | 2227205 | 0.944039 |
| 977327.3553 | *Hipposideros larvatus* | BCC-CSM2-MR_ssp585_2061-2080 | 2061-2080 | ssp585 | BCC-CSM2-MR | 3315892 | 0.977327 |
| 741458.3197 | *Hipposideros pomona* | BCC-CSM2-MR_ssp585_2061-2080 | 2061-2080 | ssp585 | BCC-CSM2-MR | 3100912 | 0.741458 |
| 4304328.703 | *Hipposideros ruber* | BCC-CSM2-MR_ssp585_2061-2080 | 2061-2080 | ssp585 | BCC-CSM2-MR | 8487199 | 4.304329 |
| 115548.1465 | *Miniopterus schreibersii* | BCC-CSM2-MR_ssp585_2061-2080 | 2061-2080 | ssp585 | BCC-CSM2-MR | 2816421 | 0.115548 |
| 332172.4836 | *Nyctalus leisleri* | BCC-CSM2-MR_ssp585_2061-2080 | 2061-2080 | ssp585 | BCC-CSM2-MR | 7413692 | 0.332172 |
| 1053318.654 | *Plecotus auritus* | BCC-CSM2-MR_ssp585_2061-2080 | 2061-2080 | ssp585 | BCC-CSM2-MR | 9355015 | 1.053319 |
| 983416.8336 | *Rhinolophus affinis* | BCC-CSM2-MR_ssp585_2061-2080 | 2061-2080 | ssp585 | BCC-CSM2-MR | 5133788 | 0.983417 |

| | | | | | | | |
|---|---|---|---|---|---|---|---|
| 781041.1491 | *Rhinolophus blasii* | BCC-CSM2-MR_ssp585_2061-2080 | 2061-2080 | ssp585 | BCC-CSM2-MR | 4751972 | 0.781041 |
| 108243.862 | *Rhinolophus euryale* | BCC-CSM2-MR_ssp585_2061-2080 | 2061-2080 | ssp585 | BCC-CSM2-MR | 4353786 | 0.108244 |
| 1063967.678 | *Rhinolophus ferrumequinum* | BCC-CSM2-MR_ssp585_2061-2080 | 2061-2080 | ssp585 | BCC-CSM2-MR | 12254734 | 1.063968 |
| 461479.6816 | *Rhinolophus hipposideros* | BCC-CSM2-MR_ssp585_2061-2080 | 2061-2080 | ssp585 | BCC-CSM2-MR | 6445160 | 0.46148 |
| 591933.6896 | *Rhinolophus macrotis* | BCC-CSM2-MR_ssp585_2061-2080 | 2061-2080 | ssp585 | BCC-CSM2-MR | 3677114 | 0.591934 |
| 421496.6656 | *Rhinolophus malayanus* | BCC-CSM2-MR_ssp585_2061-2080 | 2061-2080 | ssp585 | BCC-CSM2-MR | 1452518 | 0.421497 |
| 41800.21332 | *Rhinolophus mehelyi* | BCC-CSM2-MR_ssp585_2061-2080 | 2061-2080 | ssp585 | BCC-CSM2-MR | 3479949 | 0.0418 |
| 378226.3159 | *Rhinolophus pearsonii* | BCC-CSM2-MR_ssp585_2061-2080 | 2061-2080 | ssp585 | BCC-CSM2-MR | 2947268 | 0.378226 |
| 630977.1875 | *Rhinolophus pusillus* | BCC-CSM2-MR_ssp585_2061-2080 | 2061-2080 | ssp585 | BCC-CSM2-MR | 4716619 | 0.630977 |
| 231884.0494 | *Rhinolophus sinicus* | BCC-CSM2-MR_ssp585_2061-2080 | 2061-2080 | ssp585 | BCC-CSM2-MR | 2907524 | 0.231884 |
| 420775.1273 | *Rhinolophus thomasi* | BCC-CSM2-MR_ssp585_2061-2080 | 2061-2080 | ssp585 | BCC-CSM2-MR | 2802593 | 0.420775 |
| 92161.9739 | *Tadarida teniotis* | BCC-CSM2-MR_ssp585_2061-2080 | 2061-2080 | ssp585 | BCC-CSM2-MR | 7994697 | 0.092162 |

| | | | | | | | |
|---|---|---|---|---|---|---|---|
| 1354674.224 | *Aselliscus stoliczkanus* | BCC-CSM2-MR_ssp585_2081-2100 | 2081-2100 | ssp585 | BCC-CSM2-MR | 2476390 | 1.354674 |
| 1244615.167 | *Chaerephon plicatus* | BCC-CSM2-MR_ssp585_2081-2100 | 2081-2100 | ssp585 | BCC-CSM2-MR | 2825780 | 1.244615 |
| 1575768.918 | *Hipposideros armiger* | BCC-CSM2-MR_ssp585_2081-2100 | 2081-2100 | ssp585 | BCC-CSM2-MR | 4137359 | 1.575769 |
| 947935.4661 | *Hipposideros galeritus* | BCC-CSM2-MR_ssp585_2081-2100 | 2081-2100 | ssp585 | BCC-CSM2-MR | 2227205 | 0.947935 |
| 866014.1056 | *Hipposideros larvatus* | BCC-CSM2-MR_ssp585_2081-2100 | 2081-2100 | ssp585 | BCC-CSM2-MR | 3315892 | 0.866014 |
| 588897.5363 | *Hipposideros pomona* | BCC-CSM2-MR_ssp585_2081-2100 | 2081-2100 | ssp585 | BCC-CSM2-MR | 3100912 | 0.588898 |
| 4380332.978 | *Hipposideros ruber* | BCC-CSM2-MR_ssp585_2081-2100 | 2081-2100 | ssp585 | BCC-CSM2-MR | 8487199 | 4.380333 |
| 25114.56252 | *Miniopterus schreibersii* | BCC-CSM2-MR_ssp585_2081-2100 | 2081-2100 | ssp585 | BCC-CSM2-MR | 2816421 | 0.025115 |
| 267786.8967 | *Nyctalus leisleri* | BCC-CSM2-MR_ssp585_2081-2100 | 2081-2100 | ssp585 | BCC-CSM2-MR | 7413692 | 0.267787 |
| 1153915.414 | *Plecotus auritus* | BCC-CSM2-MR_ssp585_2081-2100 | 2081-2100 | ssp585 | BCC-CSM2-MR | 9355015 | 1.153915 |
| 774616.0758 | *Rhinolophus affinis* | BCC-CSM2-MR_ssp585_2081-2100 | 2081-2100 | ssp585 | BCC-CSM2-MR | 5133788 | 0.774616 |
| 740211.7192 | *Rhinolophus blasii* | BCC-CSM2-MR_ssp585_2081-2100 | 2081-2100 | ssp585 | BCC-CSM2-MR | 4751972 | 0.740212 |

| | | | | | | | |
|---|---|---|---|---|---|---|---|
| 26370.68138 | *Rhinolophus euryale* | BCC-CSM2-MR_ssp585_2081-2100 | 2081-2100 | ssp585 | BCC-CSM2-MR | 4353786 | 0.026371 |
| 879230.5287 | *Rhinolophus ferrumequinum* | BCC-CSM2-MR_ssp585_2081-2100 | 2081-2100 | ssp585 | BCC-CSM2-MR | 12254734 | 0.879231 |
| 362834.7095 | *Rhinolophus hipposideros* | BCC-CSM2-MR_ssp585_2081-2100 | 2081-2100 | ssp585 | BCC-CSM2-MR | 6445160 | 0.362835 |
| 514795.318 | *Rhinolophus macrotis* | BCC-CSM2-MR_ssp585_2081-2100 | 2081-2100 | ssp585 | BCC-CSM2-MR | 3677114 | 0.514795 |
| 351169.8495 | *Rhinolophus malayanus* | BCC-CSM2-MR_ssp585_2081-2100 | 2081-2100 | ssp585 | BCC-CSM2-MR | 1452518 | 0.35117 |
| 17856.66212 | *Rhinolophus mehelyi* | BCC-CSM2-MR_ssp585_2081-2100 | 2081-2100 | ssp585 | BCC-CSM2-MR | 3479949 | 0.017857 |
| 299161.9401 | *Rhinolophus pearsonii* | BCC-CSM2-MR_ssp585_2081-2100 | 2081-2100 | ssp585 | BCC-CSM2-MR | 2947268 | 0.299162 |
| 534844.0998 | *Rhinolophus pusillus* | BCC-CSM2-MR_ssp585_2081-2100 | 2081-2100 | ssp585 | BCC-CSM2-MR | 4716619 | 0.534844 |
| 176958.9014 | *Rhinolophus sinicus* | BCC-CSM2-MR_ssp585_2081-2100 | 2081-2100 | ssp585 | BCC-CSM2-MR | 2907524 | 0.176959 |
| 328172.4746 | *Rhinolophus thomasi* | BCC-CSM2-MR_ssp585_2081-2100 | 2081-2100 | ssp585 | BCC-CSM2-MR | 2802593 | 0.328172 |
| 36796.23412 | *Tadarida teniotis* | BCC-CSM2-MR_ssp585_2081-2100 | 2081-2100 | ssp585 | BCC-CSM2-MR | 7994697 | 0.036796 |
| 1219697.883 | *Aselliscus stoliczkanus* | CanESM5_ssp245_2021-2040 | 2021-2040 | ssp245 | CanESM5 | 2476390 | 1.219698 |
| 1268194.172 | *Chaerephon plicatus* | CanESM5_ssp245_2021-2040 | 2021-2040 | ssp245 | CanESM5 | 2825780 | 1.268194 |

| | | | | | | | |
|---|---|---|---|---|---|---|---|
| 1518397.099 | *Hipposideros armiger* | CanESM5_ssp245_2021-2040 | 2021-2040 | ssp245 | CanESM5 | 4137359 | 1.518397 |
| 875613.1822 | *Hipposideros galeritus* | CanESM5_ssp245_2021-2040 | 2021-2040 | ssp245 | CanESM5 | 2227205 | 0.875613 |
| 885492.9036 | *Hipposideros larvatus* | CanESM5_ssp245_2021-2040 | 2021-2040 | ssp245 | CanESM5 | 3315892 | 0.885493 |
| 574749.2458 | *Hipposideros pomona* | CanESM5_ssp245_2021-2040 | 2021-2040 | ssp245 | CanESM5 | 3100912 | 0.574749 |
| 4779078.168 | *Hipposideros ruber* | CanESM5_ssp245_2021-2040 | 2021-2040 | ssp245 | CanESM5 | 8487199 | 4.779078 |
| 382002.2729 | *Miniopterus schreibersii* | CanESM5_ssp245_2021-2040 | 2021-2040 | ssp245 | CanESM5 | 2816421 | 0.382002 |
| 631472.7607 | *Nyctalus leisleri* | CanESM5_ssp245_2021-2040 | 2021-2040 | ssp245 | CanESM5 | 7413692 | 0.631473 |
| 1365689.522 | *Plecotus auritus* | CanESM5_ssp245_2021-2040 | 2021-2040 | ssp245 | CanESM5 | 9355015 | 1.36569 |
| 1043424.537 | *Rhinolophus affinis* | CanESM5_ssp245_2021-2040 | 2021-2040 | ssp245 | CanESM5 | 5133788 | 1.043425 |
| 1148848.679 | *Rhinolophus blasii* | CanESM5_ssp245_2021-2040 | 2021-2040 | ssp245 | CanESM5 | 4751972 | 1.148849 |
| 468368.3742 | *Rhinolophus euryale* | CanESM5_ssp245_2021-2040 | 2021-2040 | ssp245 | CanESM5 | 4353786 | 0.468368 |
| 1681468.626 | *Rhinolophus ferrumequinum* | CanESM5_ssp245_2021-2040 | 2021-2040 | ssp245 | CanESM5 | 12254734 | 1.681469 |
| 926414.8336 | *Rhinolophus hipposideros* | CanESM5_ssp245_2021-2040 | 2021-2040 | ssp245 | CanESM5 | 6445160 | 0.926415 |
| 617538.4346 | *Rhinolophus macrotis* | CanESM5_ssp245_2021-2040 | 2021-2040 | ssp245 | CanESM5 | 3677114 | 0.617538 |
| 356040.7847 | *Rhinolophus malayanus* | CanESM5_ssp245_2021-2040 | 2021-2040 | ssp245 | CanESM5 | 1452518 | 0.356041 |
| 150537.2831 | *Rhinolophus mehelyi* | CanESM5_ssp245_2021-2040 | 2021-2040 | ssp245 | CanESM5 | 3479949 | 0.150537 |

| | | | | | | | |
|---|---|---|---|---|---|---|---|
| 352559.3836 | *Rhinolophus pearsonii* | CanESM5_ssp245_2021-2040 | 2021-2040 | ssp245 | CanESM5 | 2947268 | 0.352559 |
| 642718.1473 | *Rhinolophus pusillus* | CanESM5_ssp245_2021-2040 | 2021-2040 | ssp245 | CanESM5 | 4716619 | 0.642718 |
| 203681.5755 | *Rhinolophus sinicus* | CanESM5_ssp245_2021-2040 | 2021-2040 | ssp245 | CanESM5 | 2907524 | 0.203682 |
| 361616.5306 | *Rhinolophus thomasi* | CanESM5_ssp245_2021-2040 | 2021-2040 | ssp245 | CanESM5 | 2802593 | 0.361617 |
| 279274.8496 | *Tadarida teniotis* | CanESM5_ssp245_2021-2040 | 2021-2040 | ssp245 | CanESM5 | 7994697 | 0.279275 |
| 1297465.56 | *Aselliscus stoliczkanus* | CanESM5_ssp245_2041-2060 | 2041-2060 | ssp245 | CanESM5 | 2476390 | 1.297466 |
| 1447344.846 | *Chaerephon plicatus* | CanESM5_ssp245_2041-2060 | 2041-2060 | ssp245 | CanESM5 | 2825780 | 1.447345 |
| 1677396.905 | *Hipposideros armiger* | CanESM5_ssp245_2041-2060 | 2041-2060 | ssp245 | CanESM5 | 4137359 | 1.677397 |
| 970301.199 | *Hipposideros galeritus* | CanESM5_ssp245_2041-2060 | 2041-2060 | ssp245 | CanESM5 | 2227205 | 0.970301 |
| 891664.8094 | *Hipposideros larvatus* | CanESM5_ssp245_2041-2060 | 2041-2060 | ssp245 | CanESM5 | 3315892 | 0.891665 |
| 557004.1157 | *Hipposideros pomona* | CanESM5_ssp245_2041-2060 | 2041-2060 | ssp245 | CanESM5 | 3100912 | 0.557004 |
| 4395131.903 | *Hipposideros ruber* | CanESM5_ssp245_2041-2060 | 2041-2060 | ssp245 | CanESM5 | 8487199 | 4.395132 |
| 1492525.317 | *Miniopterus schreibersii* | CanESM5_ssp245_2041-2060 | 2041-2060 | ssp245 | CanESM5 | 2816421 | 1.492525 |
| 1381563.2 | *Nyctalus leisleri* | CanESM5_ssp245_2041-2060 | 2041-2060 | ssp245 | CanESM5 | 7413692 | 1.381563 |
| 2132642.907 | *Plecotus auritus* | CanESM5_ssp245_2041-2060 | 2041-2060 | ssp245 | CanESM5 | 9355015 | 2.132643 |
| 1017546.181 | *Rhinolophus affinis* | CanESM5_ssp245_2041-2060 | 2041-2060 | ssp245 | CanESM5 | 5133788 | 1.017546 |
| 1404586.288 | *Rhinolophus blasii* | CanESM5_ssp245_2041-2060 | 2041-2060 | ssp245 | CanESM5 | 4751972 | 1.404586 |

| | | | | | | | |
|---|---|---|---|---|---|---|---|
| 1527094.852 | *Rhinolophus euryale* | CanESM5_ssp245_2041-2060 | 2041-2060 | ssp245 | CanESM5 | 4353786 | 1.527095 |
| 2843317.061 | *Rhinolophus ferrumequinum* | CanESM5_ssp245_2041-2060 | 2041-2060 | ssp245 | CanESM5 | 12254734 | 2.843317 |
| 1804505.016 | *Rhinolophus hipposideros* | CanESM5_ssp245_2041-2060 | 2041-2060 | ssp245 | CanESM5 | 6445160 | 1.804505 |
| 678818.9402 | *Rhinolophus macrotis* | CanESM5_ssp245_2041-2060 | 2041-2060 | ssp245 | CanESM5 | 3677114 | 0.678819 |
| 344962.0968 | *Rhinolophus malayanus* | CanESM5_ssp245_2041-2060 | 2041-2060 | ssp245 | CanESM5 | 1452518 | 0.344962 |
| 407209.9769 | *Rhinolophus mehelyi* | CanESM5_ssp245_2041-2060 | 2041-2060 | ssp245 | CanESM5 | 3479949 | 0.40721 |
| 394872.1023 | *Rhinolophus pearsonii* | CanESM5_ssp245_2041-2060 | 2041-2060 | ssp245 | CanESM5 | 2947268 | 0.394872 |
| 685156.2702 | *Rhinolophus pusillus* | CanESM5_ssp245_2041-2060 | 2041-2060 | ssp245 | CanESM5 | 4716619 | 0.685156 |
| 237025.5635 | *Rhinolophus sinicus* | CanESM5_ssp245_2041-2060 | 2041-2060 | ssp245 | CanESM5 | 2907524 | 0.237026 |
| 320791.212 | *Rhinolophus thomasi* | CanESM5_ssp245_2041-2060 | 2041-2060 | ssp245 | CanESM5 | 2802593 | 0.320791 |
| 708286.8066 | *Tadarida teniotis* | CanESM5_ssp245_2041-2060 | 2041-2060 | ssp245 | CanESM5 | 7994697 | 0.708287 |
| 1285422.477 | *Asellia stoliczkanus* | CanESM5_ssp245_2061-2080 | 2061-2080 | ssp245 | CanESM5 | 2476390 | 1.285422 |
| 1295570.226 | *Chaerephon plicatus* | CanESM5_ssp245_2061-2080 | 2061-2080 | ssp245 | CanESM5 | 2825780 | 1.29557 |
| 1531495.367 | *Hipposideros armiger* | CanESM5_ssp245_2061-2080 | 2061-2080 | ssp245 | CanESM5 | 4137359 | 1.531495 |
| 935456.4588 | *Hipposideros galeritus* | CanESM5_ssp245_2061-2080 | 2061-2080 | ssp245 | CanESM5 | 2227205 | 0.935456 |
| 901731.2482 | *Hipposideros larvatus* | CanESM5_ssp245_2061-2080 | 2061-2080 | ssp245 | CanESM5 | 3315892 | 0.901731 |

| | | | | | | | |
|---|---|---|---|---|---|---|---|
| 492441.3015 | *Hipposideros pomona* | CanESM5_ssp245_2061-2080 | 2061-2080 | ssp245 | CanESM5 | 3100912 | 0.492441 |
| 4917596.015 | *Hipposideros ruber* | CanESM5_ssp245_2061-2080 | 2061-2080 | ssp245 | CanESM5 | 8487199 | 4.917596 |
| 114524.7637 | *Miniopterus schreibersii* | CanESM5_ssp245_2061-2080 | 2061-2080 | ssp245 | CanESM5 | 2816421 | 0.114525 |
| 347285.8202 | *Nyctalus leisleri* | CanESM5_ssp245_2061-2080 | 2061-2080 | ssp245 | CanESM5 | 7413692 | 0.347286 |
| 1202033.56 | *Plecotus auritus* | CanESM5_ssp245_2061-2080 | 2061-2080 | ssp245 | CanESM5 | 9355015 | 1.202034 |
| 893557.1192 | *Rhinolophus affinis* | CanESM5_ssp245_2061-2080 | 2061-2080 | ssp245 | CanESM5 | 5133788 | 0.893557 |
| 739641.1602 | *Rhinolophus blasii* | CanESM5_ssp245_2061-2080 | 2061-2080 | ssp245 | CanESM5 | 4751972 | 0.739641 |
| 152105.0999 | *Rhinolophus euryale* | CanESM5_ssp245_2061-2080 | 2061-2080 | ssp245 | CanESM5 | 4353786 | 0.152105 |
| 1221522.707 | *Rhinolophus ferrumequinum* | CanESM5_ssp245_2061-2080 | 2061-2080 | ssp245 | CanESM5 | 12254734 | 1.221523 |
| 538440.6991 | *Rhinolophus hipposideros* | CanESM5_ssp245_2061-2080 | 2061-2080 | ssp245 | CanESM5 | 6445160 | 0.538441 |
| 604845.133 | *Rhinolophus macrotis* | CanESM5_ssp245_2061-2080 | 2061-2080 | ssp245 | CanESM5 | 3677114 | 0.604845 |
| 312869.1301 | *Rhinolophus malayanus* | CanESM5_ssp245_2061-2080 | 2061-2080 | ssp245 | CanESM5 | 1452518 | 0.312869 |
| 39963.69572 | *Rhinolophus mehelyi* | CanESM5_ssp245_2061-2080 | 2061-2080 | ssp245 | CanESM5 | 3479949 | 0.039964 |
| 340789.5675 | *Rhinolophus pearsonii* | CanESM5_ssp245_2061-2080 | 2061-2080 | ssp245 | CanESM5 | 2947268 | 0.34079 |
| 602311.3996 | *Rhinolophus pusillus* | CanESM5_ssp245_2061-2080 | 2061-2080 | ssp245 | CanESM5 | 4716619 | 0.602311 |
| 202221.1354 | *Rhinolophus sinicus* | CanESM5_ssp245_2061-2080 | 2061-2080 | ssp245 | CanESM5 | 2907524 | 0.202221 |

| | | | | | | | |
|---|---|---|---|---|---|---|---|
| 325162.2657 | *Rhinolophus thomasi* | CanESM5_ssp245_2061-2080 | 2061-2080 | ssp245 | CanESM5 | 2802593 | 0.325162 |
| 77652.39625 | *Tadarida teniotis* | CanESM5_ssp245_2061-2080 | 2061-2080 | ssp245 | CanESM5 | 7994697 | 0.077652 |
| 1283542.136 | *Aselliscus stoliczkanus* | CanESM5_ssp245_2081-2100 | 2081-2100 | ssp245 | CanESM5 | 2476390 | 1.283542 |
| 1341929.684 | *Chaerephon plicatus* | CanESM5_ssp245_2081-2100 | 2081-2100 | ssp245 | CanESM5 | 2825780 | 1.34193 |
| 1525905.664 | *Hipposideros armiger* | CanESM5_ssp245_2081-2100 | 2081-2100 | ssp245 | CanESM5 | 4137359 | 1.525906 |
| 941644.6284 | *Hipposideros galeritus* | CanESM5_ssp245_2081-2100 | 2081-2100 | ssp245 | CanESM5 | 2227205 | 0.941645 |
| 877330.8462 | *Hipposideros larvatus* | CanESM5_ssp245_2081-2100 | 2081-2100 | ssp245 | CanESM5 | 3315892 | 0.877331 |
| 420415.3906 | *Hipposideros pomona* | CanESM5_ssp245_2081-2100 | 2081-2100 | ssp245 | CanESM5 | 3100912 | 0.420415 |
| 4913953.613 | *Hipposideros ruber* | CanESM5_ssp245_2081-2100 | 2081-2100 | ssp245 | CanESM5 | 8487199 | 4.913954 |
| 77594.52834 | *Miniopterus schreibersii* | CanESM5_ssp245_2081-2100 | 2081-2100 | ssp245 | CanESM5 | 2816421 | 0.077595 |
| 317614.441 | *Nyctalus leisleri* | CanESM5_ssp245_2081-2100 | 2081-2100 | ssp245 | CanESM5 | 7413692 | 0.317614 |
| 1140266.655 | *Plecotus auritus* | CanESM5_ssp245_2081-2100 | 2081-2100 | ssp245 | CanESM5 | 9355015 | 1.140267 |
| 807864.7882 | *Rhinolophus affinis* | CanESM5_ssp245_2081-2100 | 2081-2100 | ssp245 | CanESM5 | 5133788 | 0.807865 |
| 712200.7953 | *Rhinolophus blasii* | CanESM5_ssp245_2081-2100 | 2081-2100 | ssp245 | CanESM5 | 4751972 | 0.712201 |
| 108303.3673 | *Rhinolophus euryale* | CanESM5_ssp245_2081-2100 | 2081-2100 | ssp245 | CanESM5 | 4353786 | 0.108303 |
| 1132127.684 | *Rhinolophus ferrumequinum* | CanESM5_ssp245_2081-2100 | 2081-2100 | ssp245 | CanESM5 | 12254734 | 1.132128 |

| | | | | | | | |
|---|---|---|---|---|---|---|---|
| 489143.2517 | *Rhinolophus hipposideros* | CanESM5_ssp245_2081-2100 | 2081-2100 | ssp245 | CanESM5 | 6445160 | 0.489143 |
| 577271.3416 | *Rhinolophus macrotis* | CanESM5_ssp245_2081-2100 | 2081-2100 | ssp245 | CanESM5 | 3677114 | 0.577271 |
| 273415.4665 | *Rhinolophus malayanus* | CanESM5_ssp245_2081-2100 | 2081-2100 | ssp245 | CanESM5 | 1452518 | 0.273415 |
| 26531.81814 | *Rhinolophus mehelyi* | CanESM5_ssp245_2081-2100 | 2081-2100 | ssp245 | CanESM5 | 3479949 | 0.026532 |
| 321332.7436 | *Rhinolophus pearsonii* | CanESM5_ssp245_2081-2100 | 2081-2100 | ssp245 | CanESM5 | 2947268 | 0.321333 |
| 568117.6959 | *Rhinolophus pusillus* | CanESM5_ssp245_2081-2100 | 2081-2100 | ssp245 | CanESM5 | 4716619 | 0.568118 |
| 197286.583 | *Rhinolophus sinicus* | CanESM5_ssp245_2081-2100 | 2081-2100 | ssp245 | CanESM5 | 2907524 | 0.197287 |
| 301548.6653 | *Rhinolophus thomasi* | CanESM5_ssp245_2081-2100 | 2081-2100 | ssp245 | CanESM5 | 2802593 | 0.301549 |
| 53172.8112 | *Tadarida teniotis* | CanESM5_ssp245_2081-2100 | 2081-2100 | ssp245 | CanESM5 | 7994697 | 0.053173 |
| 1244707.644 | *Aselliscus stoliczkanus* | CanESM5_ssp585_2021-2040 | 2021-2040 | ssp585 | CanESM5 | 2476390 | 1.244708 |
| 1250130.076 | *Chaerephon plicatus* | CanESM5_ssp585_2021-2040 | 2021-2040 | ssp585 | CanESM5 | 2825780 | 1.25013 |
| 1534834.683 | *Hipposideros armiger* | CanESM5_ssp585_2021-2040 | 2021-2040 | ssp585 | CanESM5 | 4137359 | 1.534835 |
| 901730.5344 | *Hipposideros galeritus* | CanESM5_ssp585_2021-2040 | 2021-2040 | ssp585 | CanESM5 | 2227205 | 0.901731 |
| 858669.2752 | *Hipposideros larvatus* | CanESM5_ssp585_2021-2040 | 2021-2040 | ssp585 | CanESM5 | 3315892 | 0.858669 |
| 546729.9369 | *Hipposideros pomona* | CanESM5_ssp585_2021-2040 | 2021-2040 | ssp585 | CanESM5 | 3100912 | 0.54673 |
| 4833172.232 | *Hipposideros ruber* | CanESM5_ssp585_2021-2040 | 2021-2040 | ssp585 | CanESM5 | 8487199 | 4.833172 |
| 339015.8242 | *Miniopterus schreibersii* | CanESM5_ssp585_2021-2040 | 2021-2040 | ssp585 | CanESM5 | 2816421 | 0.339016 |

| | | | | | | | |
|---|---|---|---|---|---|---|---|
| 580221.0646 | *Nyctalus leisleri* | CanESM5_ssp585_2021-2040 | 2021-2040 | ssp585 | CanESM5 | 7413692 | 0.580221 |
| 1356887.963 | *Plecotus auritus* | CanESM5_ssp585_2021-2040 | 2021-2040 | ssp585 | CanESM5 | 9355015 | 1.356888 |
| 987437.8587 | *Rhinolophus affinis* | CanESM5_ssp585_2021-2040 | 2021-2040 | ssp585 | CanESM5 | 5133788 | 0.987438 |
| 1045897.436 | *Rhinolophus blasii* | CanESM5_ssp585_2021-2040 | 2021-2040 | ssp585 | CanESM5 | 4751972 | 1.045897 |
| 417232.7274 | *Rhinolophus euryale* | CanESM5_ssp585_2021-2040 | 2021-2040 | ssp585 | CanESM5 | 4353786 | 0.417233 |
| 1634591.773 | *Rhinolophus ferrumequinum* | CanESM5_ssp585_2021-2040 | 2021-2040 | ssp585 | CanESM5 | 12254734 | 1.634592 |
| 869129.3962 | *Rhinolophus hipposideros* | CanESM5_ssp585_2021-2040 | 2021-2040 | ssp585 | CanESM5 | 6445160 | 0.869129 |
| 632521.8547 | *Rhinolophus macrotis* | CanESM5_ssp585_2021-2040 | 2021-2040 | ssp585 | CanESM5 | 3677114 | 0.632522 |
| 350844.2127 | *Rhinolophus malayanus* | CanESM5_ssp585_2021-2040 | 2021-2040 | ssp585 | CanESM5 | 1452518 | 0.350844 |
| 122031.0701 | *Rhinolophus mehelyi* | CanESM5_ssp585_2021-2040 | 2021-2040 | ssp585 | CanESM5 | 3479949 | 0.122031 |
| 346944.2757 | *Rhinolophus pearsonii* | CanESM5_ssp585_2021-2040 | 2021-2040 | ssp585 | CanESM5 | 2947268 | 0.346944 |
| 626404.8843 | *Rhinolophus pusillus* | CanESM5_ssp585_2021-2040 | 2021-2040 | ssp585 | CanESM5 | 4716619 | 0.626405 |
| 215480.0761 | *Rhinolophus sinicus* | CanESM5_ssp585_2021-2040 | 2021-2040 | ssp585 | CanESM5 | 2907524 | 0.21548 |
| 354485.657 | *Rhinolophus thomasi* | CanESM5_ssp585_2021-2040 | 2021-2040 | ssp585 | CanESM5 | 2802593 | 0.354486 |
| 246445.3172 | *Tadarida teniotis* | CanESM5_ssp585_2021-2040 | 2021-2040 | ssp585 | CanESM5 | 7994697 | 0.246445 |
| 1266654.849 | *Aselliscus stoliczkanus* | CanESM5_ssp585_2041-2060 | 2041-2060 | ssp585 | CanESM5 | 2476390 | 1.266655 |

| | | | | | | | |
|---|---|---|---|---|---|---|---|
| 1257403.514 | *Chaerephon plicatus* | CanESM5_ssp585_2041-2060 | 2041-2060 | ssp585 | CanESM5 | 2825780 | 1.257404 |
| 1533872.041 | *Hipposideros armiger* | CanESM5_ssp585_2041-2060 | 2041-2060 | ssp585 | CanESM5 | 4137359 | 1.533872 |
| 937663.6306 | *Hipposideros galeritus* | CanESM5_ssp585_2041-2060 | 2041-2060 | ssp585 | CanESM5 | 2227205 | 0.937664 |
| 904891.836 | *Hipposideros larvatus* | CanESM5_ssp585_2041-2060 | 2041-2060 | ssp585 | CanESM5 | 3315892 | 0.904892 |
| 481429.6006 | *Hipposideros pomona* | CanESM5_ssp585_2041-2060 | 2041-2060 | ssp585 | CanESM5 | 3100912 | 0.48143 |
| 4951266.561 | *Hipposideros ruber* | CanESM5_ssp585_2041-2060 | 2041-2060 | ssp585 | CanESM5 | 8487199 | 4.951267 |
| 131745.5175 | *Miniopterus schreibersii* | CanESM5_ssp585_2041-2060 | 2041-2060 | ssp585 | CanESM5 | 2816421 | 0.131746 |
| 351237.0023 | *Nyctalus leisleri* | CanESM5_ssp585_2041-2060 | 2041-2060 | ssp585 | CanESM5 | 7413692 | 0.351237 |
| 1137500.721 | *Plecotus auritus* | CanESM5_ssp585_2041-2060 | 2041-2060 | ssp585 | CanESM5 | 9355015 | 1.137501 |
| 879465.1591 | *Rhinolophus affinis* | CanESM5_ssp585_2041-2060 | 2041-2060 | ssp585 | CanESM5 | 5133788 | 0.879465 |
| 808167.1904 | *Rhinolophus blasii* | CanESM5_ssp585_2041-2060 | 2041-2060 | ssp585 | CanESM5 | 4751972 | 0.808167 |
| 170927.6309 | *Rhinolophus euryale* | CanESM5_ssp585_2041-2060 | 2041-2060 | ssp585 | CanESM5 | 4353786 | 0.170928 |
| 1269916.328 | *Rhinolophus ferrumequinum* | CanESM5_ssp585_2041-2060 | 2041-2060 | ssp585 | CanESM5 | 12254734 | 1.269916 |
| 554153.1205 | *Rhinolophus hipposideros* | CanESM5_ssp585_2041-2060 | 2041-2060 | ssp585 | CanESM5 | 6445160 | 0.554153 |
| 607342.7487 | *Rhinolophus macrotis* | CanESM5_ssp585_2041-2060 | 2041-2060 | ssp585 | CanESM5 | 3677114 | 0.607343 |
| 323938.5753 | *Rhinolophus malayanus* | CanESM5_ssp585_2041-2060 | 2041-2060 | ssp585 | CanESM5 | 1452518 | 0.323939 |

| | | | | | | | |
|---|---|---|---|---|---|---|---|
| 49739.77798 | *Rhinolophus mehelyi* | CanESM5_ssp585_2041-2060 | 2041-2060 | ssp585 | CanESM5 | 3479949 | 0.04974 |
| 339566.7717 | *Rhinolophus pearsonii* | CanESM5_ssp585_2041-2060 | 2041-2060 | ssp585 | CanESM5 | 2947268 | 0.339567 |
| 604268.2393 | *Rhinolophus pusillus* | CanESM5_ssp585_2041-2060 | 2041-2060 | ssp585 | CanESM5 | 4716619 | 0.604268 |
| 200901.9292 | *Rhinolophus sinicus* | CanESM5_ssp585_2041-2060 | 2041-2060 | ssp585 | CanESM5 | 2907524 | 0.200902 |
| 331072.3529 | *Rhinolophus thomasi* | CanESM5_ssp585_2041-2060 | 2041-2060 | ssp585 | CanESM5 | 2802593 | 0.331072 |
| 88612.22731 | *Tadarida teniotis* | CanESM5_ssp585_2041-2060 | 2041-2060 | ssp585 | CanESM5 | 7994697 | 0.088612 |
| 1332497.549 | *Aselliscus stoliczkanus* | CanESM5_ssp585_2061-2080 | 2061-2080 | ssp585 | CanESM5 | 2476390 | 1.332498 |
| 1233483.287 | *Chaerephon plicatus* | CanESM5_ssp585_2061-2080 | 2061-2080 | ssp585 | CanESM5 | 2825780 | 1.233483 |
| 1546197.114 | *Hipposideros armiger* | CanESM5_ssp585_2061-2080 | 2061-2080 | ssp585 | CanESM5 | 4137359 | 1.546197 |
| 950719.6428 | *Hipposideros galeritus* | CanESM5_ssp585_2061-2080 | 2061-2080 | ssp585 | CanESM5 | 2227205 | 0.95072 |
| 913367.8736 | *Hipposideros larvatus* | CanESM5_ssp585_2061-2080 | 2061-2080 | ssp585 | CanESM5 | 3315892 | 0.913368 |
| 400985.715 | *Hipposideros pomona* | CanESM5_ssp585_2061-2080 | 2061-2080 | ssp585 | CanESM5 | 3100912 | 0.400986 |
| 4994794.138 | *Hipposideros ruber* | CanESM5_ssp585_2061-2080 | 2061-2080 | ssp585 | CanESM5 | 8487199 | 4.994794 |
| 22375.07951 | *Miniopterus schreibersii* | CanESM5_ssp585_2061-2080 | 2061-2080 | ssp585 | CanESM5 | 2816421 | 0.022375 |
| 256272.148 | *Nyctalus leisleri* | CanESM5_ssp585_2061-2080 | 2061-2080 | ssp585 | CanESM5 | 7413692 | 0.256272 |
| 1157516.677 | *Plecotus auritus* | CanESM5_ssp585_2061-2080 | 2061-2080 | ssp585 | CanESM5 | 9355015 | 1.157517 |
| 891376.9869 | *Rhinolophus affinis* | CanESM5_ssp585_2061-2080 | 2061-2080 | ssp585 | CanESM5 | 5133788 | 0.891377 |

| | | | | | | | |
|---|---|---|---|---|---|---|---|
| 794303.8346 | *Rhinolophus blasii* | CanESM5_ssp585_2061-2080 | 2061-2080 | ssp585 | CanESM5 | 4751972 | 0.794304 |
| 60756.77425 | *Rhinolophus euryale* | CanESM5_ssp585_2061-2080 | 2061-2080 | ssp585 | CanESM5 | 4353786 | 0.060757 |
| 1125033.98 | *Rhinolophus ferrumequinum* | CanESM5_ssp585_2061-2080 | 2061-2080 | ssp585 | CanESM5 | 12254734 | 1.125034 |
| 398732.2681 | *Rhinolophus hipposideros* | CanESM5_ssp585_2061-2080 | 2061-2080 | ssp585 | CanESM5 | 6445160 | 0.398732 |
| 612112.8641 | *Rhinolophus macrotis* | CanESM5_ssp585_2061-2080 | 2061-2080 | ssp585 | CanESM5 | 3677114 | 0.612113 |
| 270329.0041 | *Rhinolophus malayanus* | CanESM5_ssp585_2061-2080 | 2061-2080 | ssp585 | CanESM5 | 1452518 | 0.270329 |
| 24594.94639 | *Rhinolophus mehelyi* | CanESM5_ssp585_2061-2080 | 2061-2080 | ssp585 | CanESM5 | 3479949 | 0.024595 |
| 314468.5093 | *Rhinolophus pearsonii* | CanESM5_ssp585_2061-2080 | 2061-2080 | ssp585 | CanESM5 | 2947268 | 0.314469 |
| 624294.3305 | *Rhinolophus pusillus* | CanESM5_ssp585_2061-2080 | 2061-2080 | ssp585 | CanESM5 | 4716619 | 0.624294 |
| 229360.2595 | *Rhinolophus sinicus* | CanESM5_ssp585_2061-2080 | 2061-2080 | ssp585 | CanESM5 | 2907524 | 0.22936 |
| 299235.5518 | *Rhinolophus thomasi* | CanESM5_ssp585_2061-2080 | 2061-2080 | ssp585 | CanESM5 | 2802593 | 0.299236 |
| 30614.35087 | *Tadarida teniotis* | CanESM5_ssp585_2061-2080 | 2061-2080 | ssp585 | CanESM5 | 7994697 | 0.030614 |
| 1427152.143 | *Aselliscus stoliczkanus* | CanESM5_ssp585_2081-2100 | 2081-2100 | ssp585 | CanESM5 | 2476390 | 1.427152 |
| 1212542.796 | *Chaerephon plicatus* | CanESM5_ssp585_2081-2100 | 2081-2100 | ssp585 | CanESM5 | 2825780 | 1.212543 |
| 1602918.654 | *Hipposideros armiger* | CanESM5_ssp585_2081-2100 | 2081-2100 | ssp585 | CanESM5 | 4137359 | 1.602919 |
| 959137.289 | *Hipposideros galeritus* | CanESM5_ssp585_2081-2100 | 2081-2100 | ssp585 | CanESM5 | 2227205 | 0.959137 |

| | | | | | | | |
|---|---|---|---|---|---|---|---|
| 872580.7 497 | *Hipposideros larvatus* | CanESM5_ssp5 85_2081-2100 | 2081-2 100 | ssp5 85 | CanE SM5 | 33158 92 | 0.872 581 |
| 316813.6 334 | *Hipposideros pomona* | CanESM5_ssp5 85_2081-2100 | 2081-2 100 | ssp5 85 | CanE SM5 | 31009 12 | 0.316 814 |
| 4971903. 893 | *Hipposideros ruber* | CanESM5_ssp5 85_2081-2100 | 2081-2 100 | ssp5 85 | CanE SM5 | 84871 99 | 4.971 904 |
| 9931.838 768 | *Miniopterus schreibersii* | CanESM5_ssp5 85_2081-2100 | 2081-2 100 | ssp5 85 | CanE SM5 | 28164 21 | 0.009 932 |
| 145887.0 314 | *Nyctalus leisleri* | CanESM5_ssp5 85_2081-2100 | 2081-2 100 | ssp5 85 | CanE SM5 | 74136 92 | 0.145 887 |
| 999590.0 653 | *Plecotus auritus* | CanESM5_ssp5 85_2081-2100 | 2081-2 100 | ssp5 85 | CanE SM5 | 93550 15 | 0.999 59 |
| 914873.4 07 | *Rhinolophus affinis* | CanESM5_ssp5 85_2081-2100 | 2081-2 100 | ssp5 85 | CanE SM5 | 51337 88 | 0.914 873 |
| 726816.4 688 | *Rhinolophus blasii* | CanESM5_ssp5 85_2081-2100 | 2081-2 100 | ssp5 85 | CanE SM5 | 47519 72 | 0.726 816 |
| 7255.995 395 | *Rhinolophus euryale* | CanESM5_ssp5 85_2081-2100 | 2081-2 100 | ssp5 85 | CanE SM5 | 43537 86 | 0.007 256 |
| 997923.6 407 | *Rhinolophus ferrumequinu m* | CanESM5_ssp5 85_2081-2100 | 2081-2 100 | ssp5 85 | CanE SM5 | 12254 734 | 0.997 924 |
| 249321.4 992 | *Rhinolophus hipposideros* | CanESM5_ssp5 85_2081-2100 | 2081-2 100 | ssp5 85 | CanE SM5 | 64451 60 | 0.249 321 |
| 638687.9 421 | *Rhinolophus macrotis* | CanESM5_ssp5 85_2081-2100 | 2081-2 100 | ssp5 85 | CanE SM5 | 36771 14 | 0.638 688 |
| 193161.5 365 | *Rhinolophus malayanus* | CanESM5_ssp5 85_2081-2100 | 2081-2 100 | ssp5 85 | CanE SM5 | 14525 18 | 0.193 162 |
| 4659.059 48 | *Rhinolophus mehelyi* | CanESM5_ssp5 85_2081-2100 | 2081-2 100 | ssp5 85 | CanE SM5 | 34799 49 | 0.004 659 |
| 298357.1 419 | *Rhinolophus pearsonii* | CanESM5_ssp5 85_2081-2100 | 2081-2 100 | ssp5 85 | CanE SM5 | 29472 68 | 0.298 357 |
| 626759.0 574 | *Rhinolophus pusillus* | CanESM5_ssp5 85_2081-2100 | 2081-2 100 | ssp5 85 | CanE SM5 | 47166 19 | 0.626 759 |

| | | | | | | | |
|---|---|---|---|---|---|---|---|
| 249762.3981 | *Rhinolophus sinicus* | CanESM5_ssp585_2081-2100 | 2081-2100 | ssp585 | CanESM5 | 2907524 | 0.249762 |
| 253966.5328 | *Rhinolophus thomasi* | CanESM5_ssp585_2081-2100 | 2081-2100 | ssp585 | CanESM5 | 2802593 | 0.253967 |
| 13831.06697 | *Tadarida teniotis* | CanESM5_ssp585_2081-2100 | 2081-2100 | ssp585 | CanESM5 | 7994697 | 0.013831 |

**Table S7.** Population trends (IUCN Red List, access in 28/11/2021) ordered by decreasing to increasing trends. * indicates not enough data to be modelled.

| Species | Population trend |
|---|---|
| *Miniopterus schreibersii* | Decreasing |
| *Rhinolophus blasii* | Decreasing |
| *Rhinolophus creaghi* | Decreasing |
| *Rhinolophus euryale* | Decreasing |
| *Rhinolophus ferrumequinum* | Decreasing |
| *Rhinolophus hipposideros* | Decreasing |
| *Rhinolophus mehelyi* | Decreasing |
| *Rhinolophus rex** | Decreasing |
| *Plecotus auritus* | Stable |
| *Rhinolophus affinis* | Stable |
| *Rhinolophus macrotis* | Stable |
| *Rhinolophus malayanus* | Stable |
| *Rhinolophus pusillus* | Stable |
| *Rhinolophus shameli** | Stable |
| *Rhinolophus stheno** | Stable |
| *Aselliscus stoliczkanus* | Unknown |
| *Chaerephon plicatus* | Unknown |
| *Hipposideros armiger* | Unknown |
| *Hipposideros galeritus* | Unknown |
| *Hipposideros larvatus* | Unknown |
| *Hipposideros pomona (gentilis)** | Unknown |
| *Hipposideros pratti** | Unknown |

| | |
|---|---|
| *Hipposideros ruber* | Unknown |
| *Nyctalus leisleri* | Unknown |
| *Rhinolophus acuminatus** | Unknown |
| *Rhinolophus blythi* | Unknown |
| *Rhinolophus cornutus** | Unknown |
| *Rhinolophus luctus* | Unknown |
| *Rhinolophus marshalli** | Unknown |
| *Rhinolophus monoceros** | Unknown |
| *Rhinolophus pearsonii* | Unknown |
| *Rhinolophus siamensis* | Unknown |
| *Rhinolophus sinicus* | Unknown |
| *Rhinolophus thomasi* | Unknown |
| *Tadarida teniotis* | Unknown |
